# Genomic and genetic insights into speciation and pigment pattern diversification in *Danio* fishes

**DOI:** 10.1101/2025.10.17.682988

**Authors:** Jianguo Lu, Marco Podobnik, Junrou Huang, Braedan M. McCluskey, Shane A. McCarthy, Jonathan Wood, Joanna Collins, James Torrance, Ying Sims, Dong Gao, Jing Huang, Jia Liu, Wenyu Fang, Peilin Huang, Chunlei Ma, David Parichy, Uwe Irion, Jian Liu, Kerstin Howe, John H. Postlethwait

## Abstract

The Danioninae subfamily of teleost fishes boasts up to four hundred distinct species that have evolved to display a stunning diversity of morphological forms. Here we use newly assembled genome sequences of four laboratory and wild zebrafish strains as well as eleven species of the *Danio* and *Danionella* genera to explore their phylogenetic history and the genetic basis of pigment pattern diversification. Phylogenomic analyses uncover extensive introgression and incomplete lineage sorting that have obscured phylogenetic relationships within *Danio* and corroborate an ancient hybrid origin of zebrafish. Whereas *D. rerio* inherited ancestral horizontal stripes, relatives repeatedly evolved spots and vertical bars. Interspecific complementation tests reveal functional divergence of the adhesion molecule gene *igsf11* and the gap junction gene *gja5b* between the striped zebrafish and *Danio* species with divergent patterns. Comparative genomic and transcriptomic analyses suggest that protein and regulatory evolution have accompanied pigment pattern diversification. Our analyses elucidate complex genetic changes underlying the phylogenetic history and morphological diversification in the *Danio* genus. Resolved phylogenetic relationships, available genome assemblies, transcriptomes, and genetic tractability establish *Danio* fish species as excellent models for biomedical research in vertebrates.

## Introduction

Small freshwater fish species of the genus *Danio* are becoming important model organisms for evolutionary developmental biology given their long-established biomedical model zebrafish, *Danio rerio*^1^. Recently demonstrated complexity in phylogenetic relationships within *Danio* suggest that increasing the level of resolution of historic and current relationships among species could provide new insights into evolutionary changes as the *Danio* clade diversified^2,3^. One of the most variable traits within the genus is pigmentation, with different species exhibiting a diversity of pigment patterns, including horizontal stripes (*D. rerio, D. kyathit*), vertical bars (*D. aesculapii, D. choprae, D. erythromicron*) and spots (*D. tinwini, D. margaritatus*). Other species have a mix of stripes and spots (*D. nigrofasciatus*, *D. dangila*) or have a nearly uniform pattern (*D. albolineatus*).

Pigment patterns play key roles in camouflage, kin recognition, and mate choice. They are targets for natural and sexual selection, and therefore of high evolutionary significance^4,5^. The *Danio* model genus offers an excellent opportunity to unravel the genetic mechanisms underlying pigment pattern evolution^6,7^. An ancestral state reconstruction suggests that the most plausible scenario for pattern diversification in *Danio* is repeated evolution from an inferred ancestral pattern of horizontal stripes reminiscent of zebrafish towards the various alternative pattern states now evident^8^. Analysis of hybrids between *Danio* species further suggests a repeated and independent evolution of vertical bars^9^. In zebrafish, interactions between the three major pigment cell types – black melanophores, orange/yellow xanthophores and silvery iridophores – are required for stripe formation. Four genes – *kcnj13*, *igsf11*, *gja4* and *gja5b* – encode membrane proteins contributing to cell-cell interactions among the different pigment cell types that influence their shapes and locations. The *kcnj13* gene encodes a potassium channel required in melanophores for interactions with xanthophores and iridophores, regulating the shapes of all three pigment cell types^10–13^. This gene has functionally diverged so as to promote the formation of stripes in zebrafish but bars in *D. aesculapii*^9^. Igsf11 is an immunoglobulin superfamily adhesion protein required for migration and survival of melanophores^14^. Gja4 and Gja5b are gap junction proteins that are required autonomously by melanophores and xanthophores for homotypic and heterotypic interactions, i.e., interactions between the same or different pigment cell types, respectively^15,16^. The *igsf11*, *gja4* and *gja5b* genes are essential for pigment patterning in *D. aesculapii* as in zebrafish but do not contribute to the patterning differences between these species^9^. Whether these genes contribute to the formation and evolution of divergent patterns in other *Danio* species is not known. Likewise, the identities of other genes that contribute to pattern variation through alterations in structure and function of their products or changes in their expression remain largely unknown^17,18^.

Here, we first analyzed relationships among phylogeny, biogeography and population dynamics for several *Danio* species. Further demonstrating their utility for comparative evolutionary studies, we used genus-wide interspecific complementation tests in hybrids between *D. rerio* and nine other *Danio* species to identify genes that have functionally diverged to contribute to pigment patterning differences between species. Our findings suggest that divergence in *igsf11* and *gja5b* have contributed to pattern diversification between zebrafish and other *Danio* species with divergent patterns. Leveraging the newly available genome sequences and transcriptomes of other *Danio* species identified signs of selection in these genes as well as changes to inferred protein sequences. Using transcriptomic analyses of interspecific hybrids, we further identify genes likely to have undergone cis regulatory changes affecting their levels of expression in pigment cells and skin cells. Together, our findings provide new insights into *Danio* genomes and how they have evolved, and identify new candidate genes that may have contributed to differences in pigmentation among these species. The genomic resources this project provides will enable research into the evolution of additional species-specific traits in this group of fishes.

## Results

### 1. Genome assembly and gene annotation

We produced *de novo* genome assemblies of three *D. rerio* strains and eleven species within the genus *Danio* and in the closely related genus of dwarf fishes *Danionella*^19^. Apart from the *D. rerio* reference genome (GRCz11), our collection includes the laboratory *D*. *rerio* strains AB, Nadia (NA), and Cooch Behar (CB)^20^, as well as eight *Danio* species^20–22^ and two *Danionella* species (*D*. *cerebrum*^23^ and *D*. *dracula*^24^). The *Danio* species can be grouped into two clades: six species belong to the Rerio clade (*D*. *rerio*, *D*. *aesculapii*, *D*. *kyathit*, *D*. *tinwini*, *D*. *nigrofasciatus*, *D*. *albolineatus*), and four belong to the Choprae clade (*D*. *jaintianensis*, *D*. *choprae*, *D*. *margaritatus*, *D*. *erythromicron*) (Fig. 1).

**Fig. 1:**
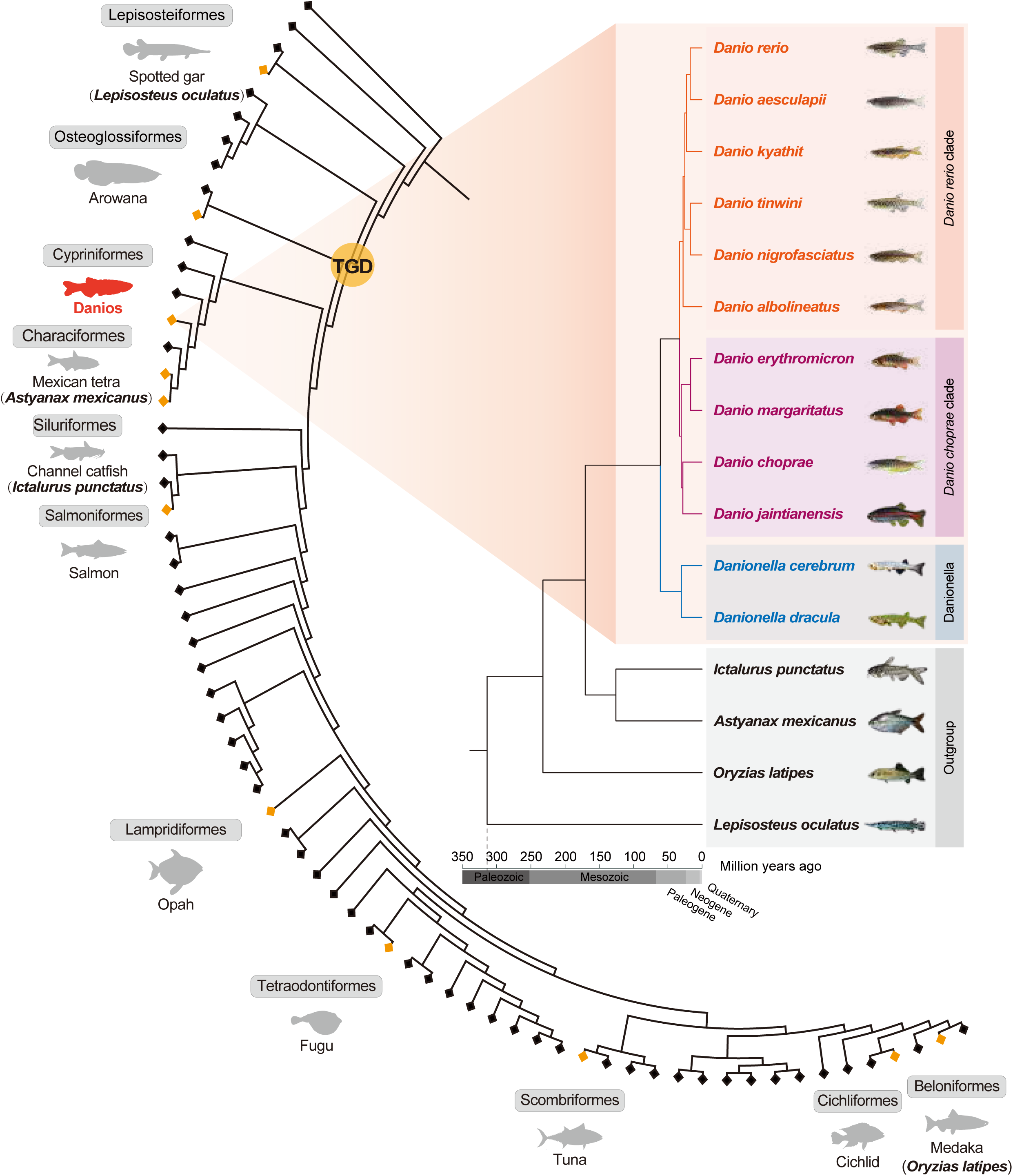
Phylogeny of *Danio* and *Danionella* genera within ray-finned fishes. Zebrafish and its relatives within the *Danio* genus can be grouped into two clades represented by Rerio clade (*D. rerio*, *D. aesculapii*, *D. kyathit*, *D. tinwini*, *D. nigrofasciatus*, *D. albolineatus*) and Choprae clade (*D. choprae*, *D. jaintianensis*, *D. margaritatus*, *D. erythromicron*). The two *Danionella* species (*D. dracula*, *D. cerebrum*) form a distinct genus, with both genera belonging to the cyprinid subfamily Danioninae, which comprise over 100 species. These species are a small subset of teleost fishes with over 30,000 species, which are all ray-finned fishes and make up about half of all vertebrate species^117^. The two most basal diverging lineages in the tree are, successively, Polypteriformes and Acipenseriformes. Orange diamonds mark the location of selected orders for orientation, with each label providing the order name and an example species. After two rounds of vertebrate-specific genome duplications (VGD, vertebrate genome duplication)^118^, one additional genome duplication occurred at the base of the teleost lineage (TGD, teleost genome duplication)^119,120^ after the split from non-teleost ray-finned fishes (represented by the spotted gar, *Lepisosteus aculatus*). The tree inference was conducted based on 1- and 2-phase sites of single-copy gene families by the Mrbayes software with GTR + Gamma; divergence times were inferred by MCMCTREE (see Methods).

We generated high-quality genome assemblies using combinations of five sequencing technologies, including Pacific Biosciences (PacBio) CLR long reads, 10x Genomics linked reads, Illumina short reads, Bionano optical maps, and Hi-C data (see Methods for detailed assembly strategies). The assembled genomes have lengths ranging from 1.4 - 1.7 Gbp for species in the Rerio clade, 1.0 - 1.1 Gbp for those in the Choprae clade, and around 0.7 Gbp for the two *Danionella* species, all with relatively high continuity and completeness.

Chromosome-level assemblies were achieved for the *D. rerio* strains (AB, NA, and CB), as well as for *D. aesculapii* and *D. kyathit*. The assemblies of *D. aesculapii* and *D. kyathit* were generated using an integrated approach that incorporated PacBio long reads, 10X Genomics Chromium, BioNano, and Hi-C data, resulting in high-quality genome assemblies as evidenced by outstanding scaffold N50, near-complete chromosomal assignment, and high Benchmarking Universal Single-Copy Orthologs (BUSCO) completeness scores. The *D. rerio* strains were originally sequenced using short-read technologies and subsequently scaffolded to chromosome level using the Sanger AB Tübingen map (SATmap)^67^. This process resulted in high scaffold continuity (Scaffold N50 = 49,100 - 51,269 kb) but intermediate BUSCO completeness scores (72.9 - 79.4 %), indicating potential residual errors or gaps despite successful chromosomal assignment.

The genome of *D. dracula*, also assembled from PacBio long reads, exhibited excellent contiguity, with a Contig N50 of 2,300 kb and a Scaffold N50 of 10,288 kb, along with high BUSCO completeness (90.3 %).

Assemblies for *D. nigrofasciatus*, *D. margaritatus*, and *D. erythromicron* were constructed using Platanus, primarily based on short-read data. These assemblies are structurally fragmented (Scaffold N50 = 152 - 659 kb; scaffold count >1.38 million), yet exhibit moderate BUSCO completeness (79.7 - 84.9 %), indicating adequate coverage of conserved gene regions. The assemblies of *D. tinwini*, *D. jaintianensis*, *D. albolineatus*, *D. choprae*, and *D. cerebrum* were generated using 10X Genomics data. Although these assemblies show limited contiguity (Scaffold N50 = 4,632 - 18,166 kb), they achieve reasonably complete gene space coverage, with BUSCO scores ranging from 78 to 91.9 %. The scaffold N50 of *Danionella cerebrum* was 6.5 Mb, 19-fold larger than a previous draft genome^25^ with a similar genome size (Table 1).

**Table 1.**
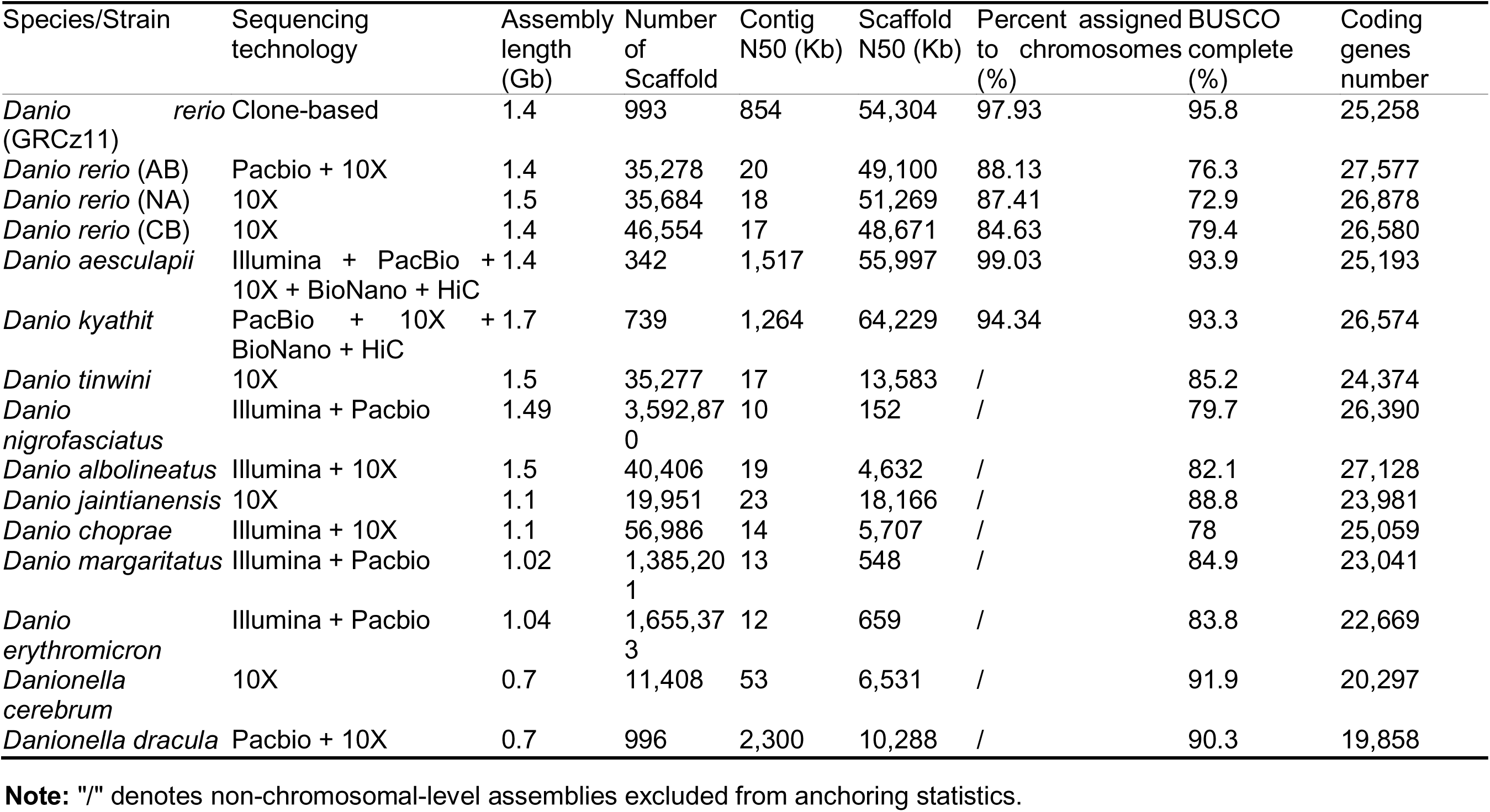
Summary of Assembly and Annotation Results of 14 Danioninae Genomes.

| Species/Strain | Sequencing technology | Assembly length (Gb) | Number of Scaffold | Contig N50 (Kb) | Scaffold N50 (Kb) | Percent assigned to chromosomes (%) | BUSCO complete (%) | Coding genes number |
| --- | --- | --- | --- | --- | --- | --- | --- | --- |
| *Danio rerio* (GRCz11) | Clone-based | 1.4 | 993 | 854 | 54,304 | 97.93 | 95.8 | 25,258 |
| *Danio rerio* (AB) | Pacbio + 10X | 1.4 | 35,278 | 20 | 49,100 | 88.13 | 76.3 | 27,577 |
| *Danio rerio* (NA) | 10X | 1.5 | 35,684 | 18 | 51,269 | 87.41 | 72.9 | 26,878 |
| *Danio rerio* (CB) | 10X | 1.4 | 46,554 | 17 | 48,671 | 84.63 | 79.4 | 26,580 |
| *Danio aesculapii* | Illumina + PacBio + 10X + BioNano + HiC | 1.4 | 342 | 1,517 | 55,997 | 99.03 | 93.9 | 25,193 |
| *Danio kyathit* | PacBio + 10X + BioNano + HiC | 1.7 | 739 | 1,264 | 64,229 | 94.34 | 93.3 | 26,574 |
| *Danio tinwini* | 10X | 1.5 | 35,277 | 17 | 13,583 | / | 85.2 | 24,374 |
| *Danio nigrofasciatus* | Illumina + Pacbio | 1.49 | 3,592,870 | 10 | 152 | / | 79.7 | 26,390 |
| *Danio albolineatus* | Illumina + 10X | 1.5 | 40,406 | 19 | 4,632 | / | 82.1 | 27,128 |
| *Danio jaintianensis* | 10X | 1.1 | 19,951 | 23 | 18,166 | / | 88.8 | 23,981 |
| *Danio choprae* | Illumina + 10X | 1.1 | 56,986 | 14 | 5,707 | / | 78 | 25,059 |
| *Danio margaritatus* | Illumina + Pacbio | 1.02 | 1,385,201 | 13 | 548 | / | 84.9 | 23,041 |
| *Danio erythromicron* | Illumina + Pacbio | 1.04 | 1,655,373 | 12 | 659 | / | 83.8 | 22,669 |
| *Danionella cerebrum* | 10X | 0.7 | 11,408 | 53 | 6,531 | / | 91.9 | 20,297 |
| *Danionella dracula* | Pacbio + 10X | 0.7 | 996 | 2,300 | 10,288 | / | 90.3 | 19,858 |

1590 **Note:** "/" denotes non-chromosomal-level assemblies excluded from anchoring statistics.

The final annotated gene numbers ranged from 19,858 to 27,577 between different species using a combination of *de novo* and homology-based gene prediction approaches, with the available protein sequences of *D. rerio*, *Oryzias latipes*, *Takifugu rubripes*, *Tetraodon nigroviridis*, and *Gasterosteus aculeatus* as references (Fig. S1-S3). There was a positive correlation between gene number and genome size.

In summary, the integration of long-read technologies with multi-platform scaffolding is essential for producing chromosome-level assemblies. Although assemblies relying on short reads or linked reads exhibit lower contiguity, they still achieve satisfactory gene-space completeness. Despite methodological variations influencing assembly structure, each approach was tailored to available data to produce high-quality genomic resources. The comprehensive datasets for species of the *Danio* genus provide a valuable foundation for comparative genomics.

### 2. Variation

Syntenies were largely conserved between *D. rerio*, *D. aesculapii* and *D. kyathit*. Nevertheless, chromosome 4 of *D. rerio* (Dre4) appears substantially elongated compared to its counterparts in the other two species. The right arm of Dre4 exhibited extensive repetitive matches to the other two genome sequences, yet harbored few orthologs or paralogs on any chromosome of *D. aesculapii* or *D. kyathit*. Instead, genes from this region of the *D. rerio* genome predominantly aligned to unanchored scaffolds from those genomes (Fig. S4). These findings suggest that the right arm of Dre4 may have undergone substantial rearrangement and expansion after the divergence of the *D. rerio* lineage from those of *D. aesculapii* and *D. kyathit*. This interpretation, however, must be considered in light of the challenging nature of this genomic region, which is notoriously difficult to assemble accurately. Nevertheless, if the assembled architecture reflects biological reality rather than an artifact, it may be associated with the acquisition of sex determination loci^26,27^ or evolution of the maternal-to-zygotic-transition block of genes in this region^28^, as well as extensive heterochromatinization^29^, thereby giving rise to a distinct sex determination mechanism in *D. rerio*^30^ relative to the other two species.

We identified 162,500,303 SNPs, 39,523,974 small insertions and deletions (indels) and 531,349 presence or absent variations (PAVs) among the 14 genome assemblies (Fig. S5). The number and distribution of SNPs varied among different *Danio* species (Fig. 2). Using the *Danio rerio* (GRCz11) reference genome, the CB strain exhibited 37 % more SNPs compared to the AB strain, and 30 % more SNPs compared to the NA strain, consistent with greater degrees of inbreeding in the latter strains. We also observed different SNP distributions among the syntenic regions of *D. rerio* strains, *D. aesculapii* and *D. kyathit*. In most cases, the SNP profiles of *Danio rerio* strains exhibited similar genomic distributions, whereas those of *D. aesculapii* and *D. kyathit* showed pronounced divergence in both SNP density and localization. Consistent with the phylogenetic relationships, the cumulative sizes of PAVs from the *Danio rerio* (GRCz11) reference were smaller among the three *D. rerio* strains (ranging between approximately 215 - 310 Mb) than the other *Danio* species. More specifically, the *D. rerio* PAVs were generally shorter in length, while the number of PAVs was almost equal to or greater than that of the other *Danio* species (Table S1). It is noteworthy that the sensitivity and specificity of PAV detection can be influenced by the quality and continuity of the genome assemblies used for comparison.

**Fig. 2:**
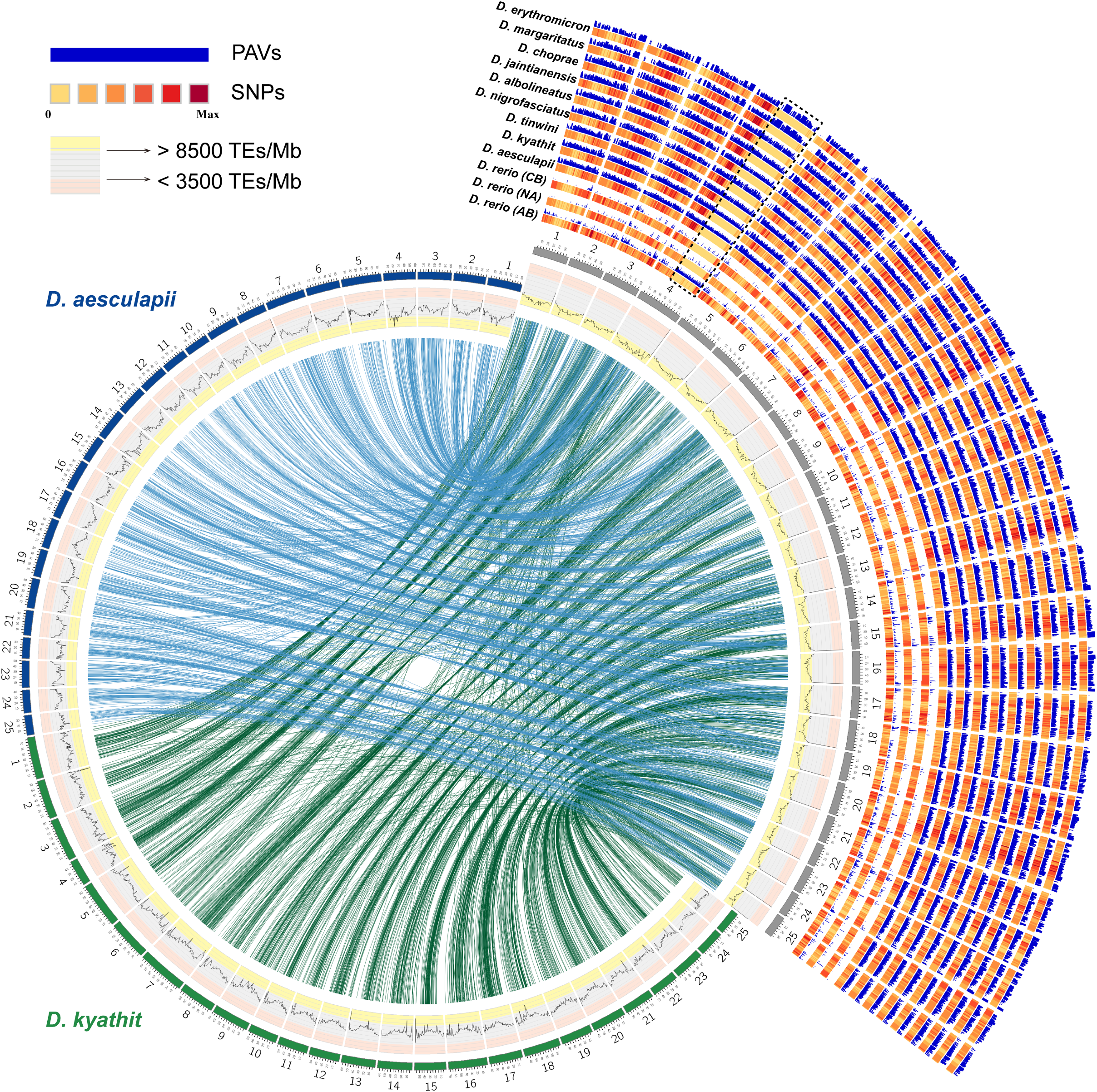
An overview of variants called from genome assemblies of zebrafish strains and other *Danio* species relative to the zebrafish reference strain AB. The circumference represents each of the 25 chromosomes in genomes of *D. rerio* (grey, right), *D. aesculapii* (blue, left) and *D. kyathit* (green, bottom) shown in a circle, with their syntenic relationships displayed in the center. The innermost track displays line graphs representing the relative density of transposable elements (TE number/Mb), binned into 1 Mb regions. To better distinguish TE abundance: regions with > 8500 TEs/Mb have a yellow background, regions with 3500–8500 TEs/Mb have a grey background, and regions with <3500 TEs/Mb have a pink background. The tracks on the right show SNPs and PAVs for three zebrafish strains and nine *Danio* species, using the zebrafish genome as the reference. The highly repetitive long arm of chr4 (boxed by dashed black lines) was not assembled in some samples, so this region is excluded from our analyses.

### 3. Phylogenetic relationships

Previous studies using reduced representation sequencing yielded conflicting models of the evolutionary history of *Danio*, with one study suggesting limited gene flow^2^ and another study supporting extensive gene flow including a hybrid origin of *D. rerio*^31^. The present genomic approach with far more data, denser taxon sampling, and improved phylogenomic methods allows us to understand the history of this group in much greater detail. For a phylogenomic analysis of *Danio* and *Danionella* species, we included outgroups of distantly related ray-finned fishes, including *Lepisosteus oculatus*, whose lineage split from teleosts before the teleost genome duplication (TGD), and *Ictalurus punctatus, Astyanax mexicanus,* and *Oryzias latipes*, whose lineages split from the *Danio* lineage at various times after the TGD. To explore phylogenies, we generated five data sets: 1) concatenated whole genome alignments (WGAs) (Fig. S6A), 2) first codon position of single-copy orthologs (Fig. S7), 3) first and second codon position of single-copy orthologs (Fig. 1, Fig. S8), 4) SNPs (Fig. S6B, S9), and 5) four-fold degenerate (4d) sites of single-copy orthologs (Fig. S6C). We reconstructed phylogenies for each data set using three methods: maximum likelihood, Bayesian and multispecies coalescent.

All five data sets supported a division of *Danio* species into two different major clades, represented by *D. rerio* and *D. choprae*, respectively (Fig. 1). In this study, the Choprae clade is represented by four species (*D. choprae, D. erythromicron, D. jaintianensis,* and *D. margaritatus*), while the Rerio clade is represented by six species (*D. rerio, D. aesculapii, D. albolineatus, D. kyathit, D. nigrofasciatus,* and *D. tinwini*). Within the Rerio clade, however, the genome-wide data sets supported three different phylogenetic topologies, differing even in their designation of the sister species of *D. rerio* (Fig. S6). In the WGA, first codon nucleotide, and first+second codon nucleotide trees, *D. kyathit* was inferred as the sister clade to (*D. rerio*, *D. aesculapii*), but distinguished from (*D. tinwini*, *D. nigrofasciatus*). The SNP tree, however, recovered *D. kyathit* as the sister species to (*D. tinwini*, *D. nigrofasciatus*), while the 4d sites tree recovered *D. kyathit* as the sister species of *D. rerio*.

The difficulty finding a single, unambiguous phylogeny for this group is consistent with a complex evolutionary history of the Rerio clade involving a hybrid origin of *D. rerio*, high rates of incomplete lineage sorting, and speciation with gene flow as suggested previously^2,31^. To investigate this history further, we generated non-overlapping 10-kb windows throughout each genome and inferred maximum-likelihood trees for each window (window-based gene trees, WGTs). The topology of the ASTRAL tree^32^ generated with WGTs by coalescent-based phylogenetic method (Fig. S10) was found to be identical to the 4d-sites tree (Fig. S6).

We constructed a DensiTree to visualize the frequency of different gene trees (Fig. 3A)^33^ and calculated quartet frequencies for the internal branches in the ASTRAL tree (Fig. 3B)^32,34^. Topological conflicts among WGTs were widely observed as visualized by DensiTree (Fig. 3A). A major discordance was the relationships among *D. rerio*, *D. aesculapii* and *D. kyathit* (Fig. 3B). The quartet probability of reconstructing the three species as sister species were very close to one third (random), with less than 5 % difference in relative frequency (Fig. 3B). The high level of phylogenetic incompatibility around the first branch might be caused by a combination of high true incompatibility and gene tree estimation errors.

**Fig. 3:**
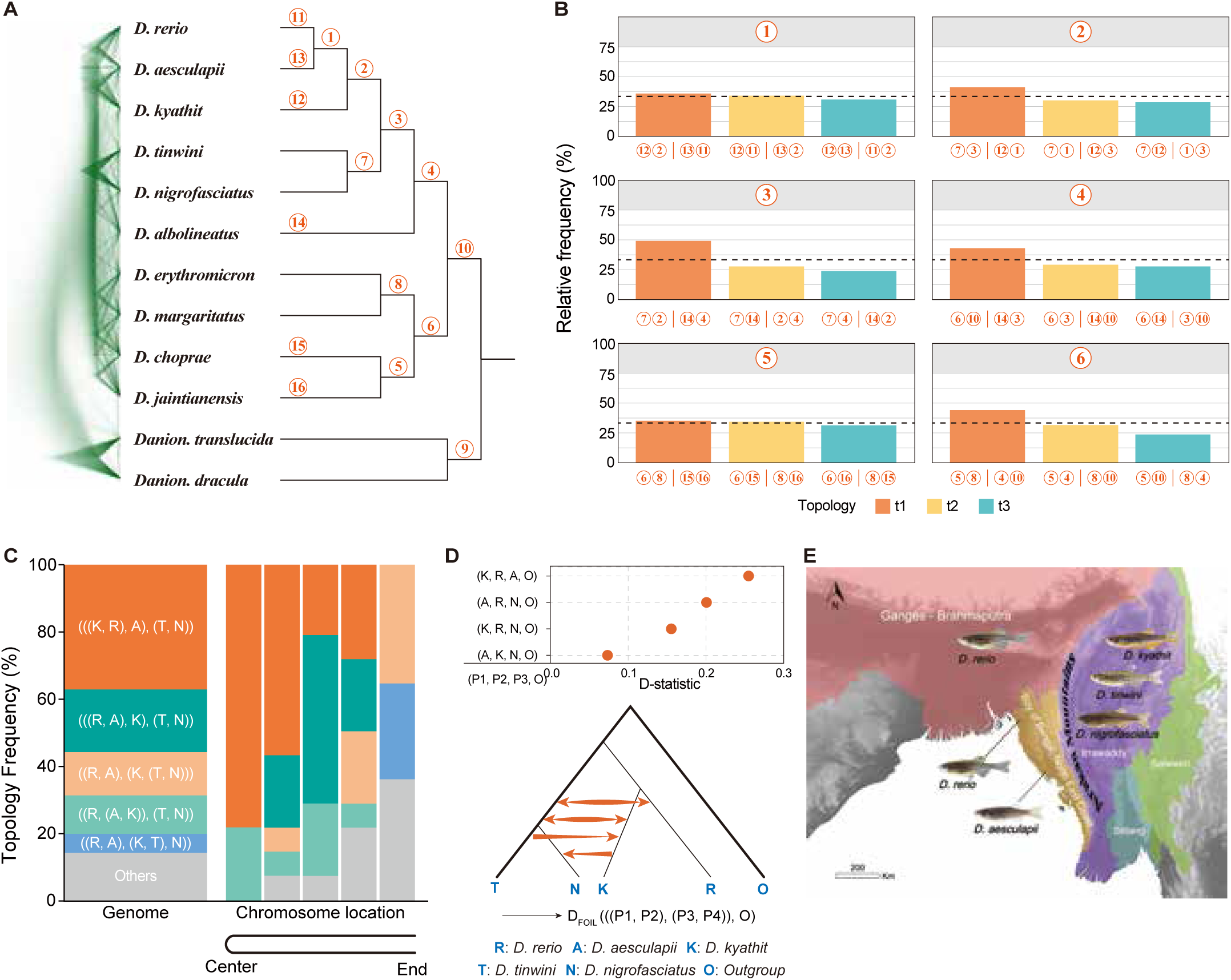
Gene-tree variability and gene flow within *Danio*. (A) Windows-based gene trees (WGTs) in a consensus DensiTree plot. The DensiTree shows phylogenetic conflicts in WGTs (left). Each of the internal branches is given a number. (B) Quartet frequencies of branches based on WGTs. Each internal branch with four neighboring branches would present three possible topologies. The frequency of the three topologies around focal internal branches of ASTRAL trees were computed using DiscoVista from the WGTs. The frequency of the most frequent topology (found in the ASTRAL tree) is t1 (Orange), and the other two alternative topologies are shown with Bisque- or CadetBlue-colored bars. The dotted line indicates the one-third threshold expected at random. Each circled number indicates the number labeled on the corresponding branch on the tree in (A). The x-axis provides the exact definition of each quartet topology using the neighboring branch labels separated by a bar “|”. Each internal branch has four neighboring branches that could be used to represent quartet topologies. Branches are collapsed at the species level and marked with numbers (A). (C) Topology frequencies inferred from genome-wide windows (left). Topology frequencies inferred from five partitioned chromosomal regions (right), where chromosomes, which are metacentric, are folded in two so that both ends of each chromosome are represented in the right-most of five columns and each chromosome’s middle is represented in the left-most column. (D) Inferred gene flow among *Danio* species using *D. albolineatus* as the outgroup. (Top) D-statistics results for the phylogeny (((P1, P2), P3), O), with each dot indicating a significant signal (p < 0.05) involving P2 and P3. (Bottom) Gene flow signals inferred by D_FOIL_ for the phylogeny (((P1, P2), (P3, P4)), O). Gene flow signals and direction are shown as orange arrows. R, *D. rerio;* K*, D. kyathit;* A*, D. aesculapii;* T*, D. tinwini;* N*, D. nigrofasciatus.* (E) Phylogeography of *Danio* species. *D. rerio* and *D. aesculapii* occupy basins West of the Arakan Mountains of Myanmar. *D. kyathit*, *D. tinwini and D. nigrofasciatus* are found in the Irrawaddy basin East of the Arakan Mountains of Myanmar.

To assess how genome structure shaped evolutionary history in Danioninae, we partitioned each chromosome into five non-overlapping regions (each representing 20 % of the sequence from telomere to centromere). Using the WGTs within each region, we inferred phylogenetic trees via ASTRAL (Table S2-S4). After excluding chromosomes with fewer than 20 WGTs, our final dataset comprised 14 chromosomes (Chr1 and Chr10-22), yielding 70 regional trees. These trees revealed strong positional effects and that topology frequencies were significantly different at different locations along chromosomes (Fig. 3C). Globally, 37.1 % of regional trees supported *D. kyathit* as sister to *D. rerio*, while 18.6 % of them supported *D. aesculapii* as sister to *D. rerio*. The dominant topologies transitioned from centromere to telomere: 1) near the centromere region, the dominant tree was (((*D. rerio*, *D. kyathit*), *D*. *aesculapii*), (*D*. *tinwini*, *D*. *nigrofasciatus*)); 2) for the intermediate region: (((*D*. *rerio*, *D*. *aesculapii*), *D*. *kyathit*), (*D*. *tinwini*, *D*. *nigrofasciatus*)); and 3) for the telomere region: ((*D*. *rerio*, *D*. *aesculapii*), (*D*. *kyathit*, (*D*. *tinwini*, *D*. *nigrofasciatus*))).

Phylogenetic relationships inferred by the telomere region of chromosomes were concordant with the present geographical distributions of *Danio* species. The Arakan Mountains separate the Ganges/Brahmaputra basin from the Irrawaddy basin. *D. rerio*^35,36^ and *D. aesculapii* ^37^ can be found in several basins that are west of the Arakan Mountains, with some basins containing both species, whereas *D. kyathit*, *D. tinwini* and *D. nigrofasciatus* inhabit the Irrawaddy basin East of the Arakan Mountains (Fig. 3E, Fig. S17).

We also investigated the possibility of gene flow contributing to the observed topological discordances. D-statistics (four-taxon)^38^ suggested possible gene flow between *D. rerio* and *D. aesculapii, D. rerio* and *D. nigrofasciatus*, and also *D. kyathit* and *D. nigrofasciatus* (Fig. 3D). Further, *D*_FOIL_ statistics (five-taxon)^39^ could be used for distinguishing incomplete lineage sorting from gene flow and determining the direction of detected introgressions. The *D*_FOIL_ statistics (Fig. 3D, Table S5, S10) revealed extensive gene flow among the Rerio clade. This evidence suggested introgression of alleles from *D. tinwini* into *D. kyathit*, and from *D. kyathit* into *D. nigrofasciatus* (Table S5, Fig. 3D, Fig. S12). Bidirectional gene flow signals were detected between (*D. tinwini*, *D. nigrofasciatus*) and *D. rerio*, also between *D. kyathit* and *D. aesculapii* (Fig. 3D, Fig. S12). *D*_FOIL_ statistics (Fig. S12) also provided evidence for gene flow among *D. rerio*, *D. kyathit* and *D. aesculapii*.

### 4. Population dynamics

Having corroborated the complex phylogenetic relationships of *Danio* species, we sought to investigate population dynamics as another means to infer the evolutionary history of these species. Different species can be divided into clades based on changes of their effective population size (*N_e_*)^40^. Using the pairwise sequentially Markovian coalescent (PSMC) method, we estimated the *N_e_* to reconstruct the demographic histories of different Danio lineages. This analysis revealed five distinct clades with divergent *N_e_* changes (Fig. S16).

The first clade included ((*D. rerio*, *D. aesculapii*), *D. kyathit*). The *N_e_* of *D. rerio* reached its peak in the Late Pleistocene interglacial stage (about 126-115 thousand years ago (kya)) and began to decrease sharply from the last glacial period (about 115 - 11.7 kya). Similarly, *D. aesculapii* and *D. kyathit* populations exhibited expansion-contraction dynamics, but their contraction phases began later than *D. rerio*’s, at 40 - 30 kya and 20 kya, respectively. The second clade included (*D. tinwini*, *D. nigrofasciatus*). Populations of these two species began to expand 20 kya, slightly in *D. tinwini*, while sharply in *D. nigrofasciatus*. The third clade (*D. albolineatus*)*: D. albolineatus* experienced population expansion during the Upper Pleistocene (126 - 11.7 kya), during which the population was relatively stable in size from 50 - 20 kya. The fourth clade (*D. erythromicron*, *D. margaritatus*): *D. erythromicron* population expanded during 200-50 kya (spanning the Late Pleistocene interglacial to early Last Glacial periods) before contracting at 50 - 40 kya, whereas *D. margaritatus* population expanded later (100 - 20 kya) with contraction initiating at 20 kya. The fifth clade, including (*D. choprae*, *D. jaintianensis*), experienced continuous population expansion during the Upper Pleistocene period (126 - 11.7 kya). These demographic patterns suggest that major evolutionary signals of the Danio species align with known climatic events.

Combining our phylogenomic analysis with geographical distribution revealed a complex evolutionary history shaped by both isolation and gene flow. The detected gene flow signals between *D. rerio* and *D. aesculapii*, and among *D. kyathit*, *D. tinwini* and *D. nigrofasciatus* are consistent with the current geographic distribution of these *Danio* species (Fig. 3E). The Arakan Mountains form the southern segment of Indo-Burman Ranges (IBR), and the uplift time of IBR may be late Oligocene (27.82 - 23.03 mya) or Oligocene-Miocene transition (23 mya)^41^. This uplift time overlaps with the divergence time of Rerio clade species, and the divergence of *D. choprae* and D. *jaintianensis* (Fig. S13-S15). The tree topologies ((*D. rerio*, *D. aesculapii*), (*D. kyathit*, (*D. tinwini*, *D. nigrofasciatus*)) and ((*D. rerio*, *D. aesculapii*), ((*D. kyathit*, *D. tinwini*), *D. nigrofasciatus*)) are both consistent with the existing distribution of the *Danio* species and are mainly recovered in regions of high recombination near the ends of *Danio* chromosomes (Fig. 3C and E). These results corroborate earlier phylogenomic analyses based on exon sequencing across 10 *Danio* species, which provided evidence supporting the hybrid origin hypothesis of *D. rerio* through ancient introgression between *D. aesculapii* and *D. kyathit* lineages^3^.

*D. jaintianensis* appeared as the sister clade to *D. choprae* in all trees, while their distribution basins were separated by the Arakan Mountains, making genetic material exchange unlikely (Fig. S17). Gene flow between the now geographically separated species may have occurred before speciation or geographical isolation. Geographical isolation may have accelerated the divergence of *D. choprae* and *D. jaintianensis*, after which gene flow occurred among species in nearby basins, respectively (Table S6). In addition, ILS caused by rapid speciation may be the cause of extensive phylogenetic incongruence among *Danio* species (Fig. S11).

### 5. Gene family expansion and contraction and positive selection

To understand how genome evolution paralleled morphological diversification, we focused on pigment patterns, one of the most divergent traits among *Danio* species. For these analyses, we restricted our investigation to eight *Danio* species exhibiting prominent black bar/stripe/spot patterns, omitting species that lacked relevant pigmentation traits. We then analyzed signatures of selection in pigment patterning genes as well as expansions and contractions of related gene families. Using protein sequences from eight *Danio* species together with the outgroup *Ictalurus punctatus*, we clustered a total of 22,444 orthogroups. The outgroup was selected for its chromosome-level genome assembly and because, as a fellow otophysan, it represents an appropriate evolutionary distance for this analysis (Fig. 1). From these orthogroups, we identified 7,460 high-confidence single-copy orthologues (SGs), which were then used to calculate dN/dS ratios across all ancestral nodes and extant species leaves to identify genes under selection pressure (Fig. S19-S22). A total of 3,578 orthologues were ultimately identified as positively selected genes (PSGs) at 7,403 nodes/leaves (Fig. S23, S24). Among the 7,460 SGs, we identified 253 pigment-related genes^42^. Of these, 174 pigment-related genes underwent positive selection at 345 of the nodes / leaves. The proportion of pigment-related genes showing positive selection ranged from 2.3 to 6.0 % depending on taxon. The dN/dS density distribution curve of the pigment-related PSGs showed a second peak above three for all nodes (Fig. 4B). The dN/dS values of pigment-related PSGs were notably higher than those of other PSGs at node 1 and also the ancestor node of Choprae clade (node 2) but were lower at the ancestor node of Rerio clade (node 6) (Fig. 4C).

**Fig. 4:**
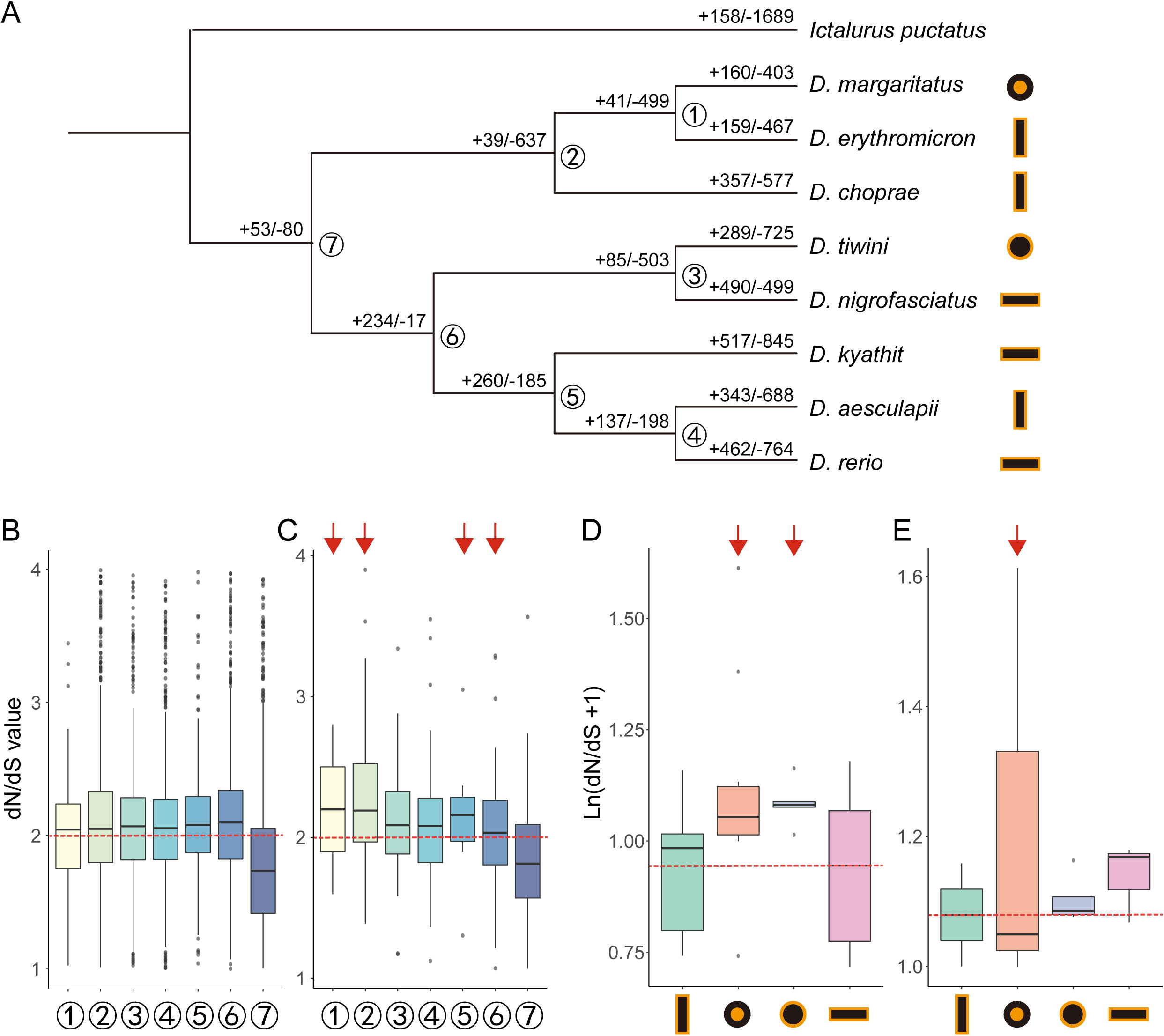
Expansion, contraction and positive selection of the orthogroups in *Danio*. (A) The number of expanded (+) and contracted (-) gene families at each node. (B) Distribution of dN/dS ratios for all positively selected genes (PSGs) across nodes (node numbering corresponds to panel A). Dashed line marks dN/dS = 2. (C) dN/dS ratios of the pigment-related PSGs at each node (node numbering corresponds to panel A). Red arrows mark specific nodes on the phylogenetic tree that are associated with shifts in dN/dS ratios of pigment-related PSGs. Specifically, the values were significantly higher at Node 1 (ancestral node) and the ancestor of the Choprae clade (node 2), but lower at the ancestor of the Rerio clade (node 6) compared to other PSGs. (D) dN/dS ratios of all pigment-related genes stratified by pigment pattern. The columns represent the pattern categories in the following order (from left to right): vertical bars (*D. aesculapii*, *D. choprae*, *D. erythromicron*), black body with yellow spots (*D. margaritatus*), yellow body with black spots (*D. tinwini*), horizontal stripes (*D. rerio*, *D. kyathit*, *D. nigrofasciatus*). (E) dN/dS ratios of pigment-related PSGs, shown by pigment pattern (categories as in D).

Of 253 pigment-related SGs, the average dN/dS value was lowest in the horizontally striped species while spotted species had higher values (Fig. 4D). The highest average dN/dS value for pigment-related PSGs was observed in *D. margaritatus*, with ectodysplasin A receptor (*edar*, dN/dS = 19.9) and engrailed homeobox 1b (*en1b*, dN/dS = 4.0) showing relatively high dN/dS values. Additionally, pigment-related PSGs in striped species showed higher dN/dS values compared to those in barred or spotted species, but since only a few such genes were detected on each leaf, the value of a single gene may have caused a disproportionate impact on the overall average (Fig. 4E).

Positive selection of *igsf11* was detected at the ancestor node of the Choprae clade as well. The *p*-value of *kcnj13* and *gja4* were all less than 0.05 at nodes 2/4/5/6, but *kcnj13* underwent positive selection at nodes 4/5/6, while *gja4* at nodes 2/4/5 (Table S8). These results indicated that pigment-related genes, i.e., including those regulating direct cell-cell interactions between pigment cells, exhibited signs of positive selection, and these genes evolved more rapidly in some species than others.

In addition, among the 22,444 orthogroups, 621 were identified as pigment-related. Beyond the 253 pigment-related SGs analyzed above, the remaining 368 orthogroups likely contain multiple gene copies or may even be completely absent in one or more species, potentially resulting from lineage-specific duplications or losses such as those mediated by the teleost-specific whole-genome duplication (TGD). Six of these 621 orthogroups show evidence of gene family expansion and contraction within *Danio* as detected by CAFÉ^43^. The histone H3f3a is essential for pigment cell development by regulating the expression of key neural crest specifier genes, such as *sox10*, through chromatin remodeling. We discovered that the *h3f3a* gene family contracted at the ancestor node of Choprae clade, resulting in fewer paralogs (Fig. 4A, Fig. S18, Table S7)^44^.

### 6. Interspecific complementation tests reveal genes involved in pigment pattern diversification

Functional tests can identify the roles these rapidly evolving genes play in development, but inducing targeted mutations in *Danio* species other than *D. rerio* becomes straightforward only with the genome sequence assemblies we provide here. Interspecific hybrids and complementation tests have provided insights into pigment pattern evolution in *Danio*, revealing for example that *kcnj13* has likely functionally diverged several times independently across lineages owing to cis regulatory changes^9,13,45^. With the availability of new genomic information, we extended such complementation testing to three additional genes, *gja4*, *gja5b* and *igsf11*, all known to regulate interactions between pigment cells during stripe formation in zebrafish ^9–16^. We crossed *D. rerio* (Tübingen/GRCz11 reference strain) heterozygous for recessive CRISPR/Cas9-induced loss-of-function alleles with each of nine other *Danio* species (Fig. 5A), including the large-bodied *D. dangila*, of potential interest for future sequencing. Wild-type hybrids between *D. rerio* and other *Danio* species generally display a pattern of horizontal stripes^6,9,46,47^, similar to the *D. rerio* pattern, with stripe aberrations sometimes evident likely depending on !genetic backgrounds of different species isolates, as depicted here for *D. tinwini* and *D. albolineatus* hybrids (Fig. 5A). In combination with previously published results, we compared altogether nine wild-type hybrids with 36 hemizygous hybrids carrying loss-of-function alleles in one of four *D. rerio* genes, *kcnj13*, *gja4*, *gja5b* and *igsf11*. In most cases (25 out of 36, 69.4 %), we observed no differences between wild-type and hemizygous test hybrids, suggesting that functions of these genes are largely conserved between *D. rerio* and these other species. Of the remaining combinations, six (16.7 % of total) showed variant stripe pattern (circles with dashed white outlines in Fig. 5A), but with enough variability that phenotypes could also overlap with those of control hybrids, suggesting genetic polymorphisms in modifier loci within species^48^. In five cases (13.9 % of total), hemizygous hybrids developed meandering patterns and broken stripes distinct from control hybrids (circles with solid white outlines in Fig. 5A). These results suggest that wild-type alleles from these other *Danio* species cannot fully complement the loss of function, and otherwise recessive, *D. rerio* alleles, and are thus likely to be hypomorphic relative to wild-type *D. rerio* alleles. Particularly notable were the phenotypes of *gja5b* and *igsf11* mutant *D. rerio* with the normally spotted species, *D. margaritatus* (Fig. 5B and C). Together, these results identify potential functional differences in *kcnj13*, *gja4*, *gja5b* and *igsf11* in contributing to pattern differences between *D. rerio* and other species, and support prior inferences of functional differences in *kcnj13* between *D. rerio* and *D. aesculapii*^9^. Our finding that the observed non-complementation phenotypes of hemizygous hybrids were not as severe as those of homozygous *D. rerio* mutants (Fig. 5A, bottom) further implies that these loci have some activity in pigment cells of the other species and that functions of these genes in pigment pattern predate the origin of the *Danio* genus.

**Fig. 5:**
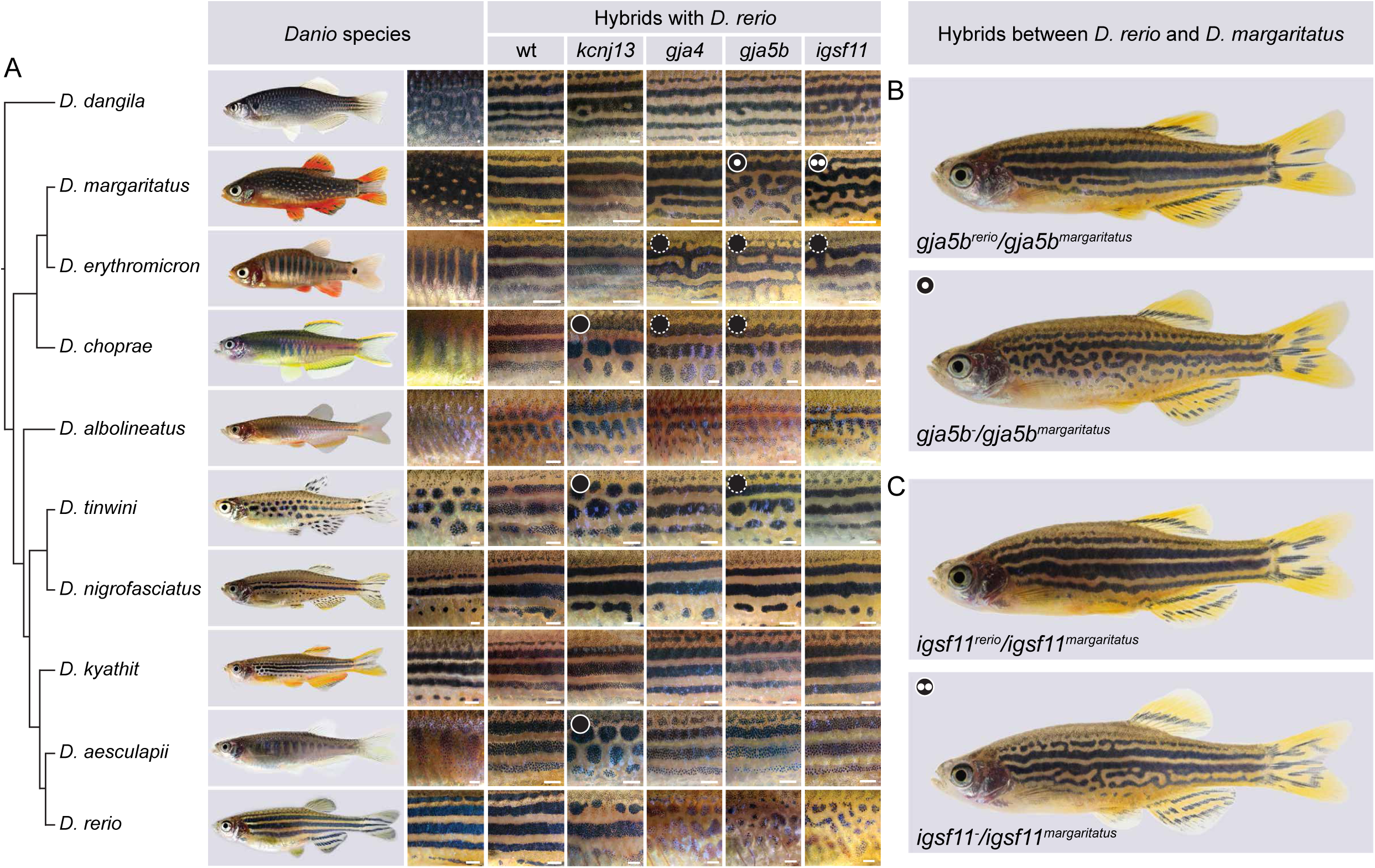
Genus-wide one-way complementation tests suggest divergence in *gja5b* and *igsf11* between *D. rerio* and *D. margaritatus*. (A) On the left, the phylogenetic tree depicts the relationships between the nine *Danio* species tested; pigment patterns of each species are shown in the first column of square boxes. The second column shows pigment patterns of hybrids between wild-type *D. rerio* and wild-type individuals of the other nine species. The lowest row shows pigment patterns of *D. rerio* that are homozygous for the mutant alleles used in this study. In addition to pattern variations observed in hybrids between *D. rerio kcnj13* mutants and *D. aesculapii*, *D. tinwini* and *D. choprae* wild types, we find pattern defects in two more cases (marked with black dots with solid outline): hybrids between *D. margaritatus* and (B) *D. rerio gja5b* (n=7) and (C) *igsf11* mutants (n=14), respectively. In six cases, patterns in hemizygous hybrids differed less clearly from wild-type hybrids (marked with black dots with dashed outline: *gja4^-/choprae^*, n=6; *gja4^-/erythromicron^*, n=14; *gja5b^-/tinwini^*, n=15; *gja5b^-/choprae^*, n=5; *gja5b^-/erythromicron^*, n=11; *igsf11^-/erythromicron^*, n=7). In all other 16 cases the patterns of the hemizygous hybrids did not differ from wild-type hybrids (*gja4^-/kyathit^*, n=5; *gja4^-/nigrofasciatus^*, n=7; *gja4^-/tinwini^*, n=12; *gja4^-/albolineatus^*, n=13; *gja4^-/margaritatus^*, n=6; *gja4^-/dangila^*, n=5; *gja5b^-/kyathit^*, n=4; *gja5b^-/nigrofasciatus^*, n=28; *gja5b^-/albolineatus^*, n=15; *gja5b^-/dangila^*, n=3; *igsf11^-/kyathit^*, n=21; *igsf11^-/nigrofasciatus^*, n=21; *igsf11^-/tinwini^*, n=21; *igsf11^-/albolineatus^*, n=3; *igsf11^-/choprae^*, n=24; *igsf11^-/dangila^*, n=6). Hybrids between the four *D. rerio* mutants and *D. aesculapii* were tested previously^9^. All pictures show representative examples of the corresponding species/hybrids/genotypes. Scale bars correspond to approximately 1 mm.

### 7. Insights into protein evolution relative to pigment pattern diversification

Two *Danio* species have independently evolved spotted patterns, *D. tinwini* with spots of black melanophores and *D. margaritatus* with spots of iridophores (Fig. 5A). Hybrids between these species develop a pattern different from either parental pattern indicating differences in underlying mechanisms^9^. As our interspecific complementation tests point towards *gja5b* as a hub for spot formation in *Danio*, with non-complementation phenotypes evident for both of these species, we asked whether protein sequence alterations could underlie the functional divergence. To this end, we reconstructed ancestral Gja5b protein sequences within *Danio* and assessed the accumulation of non-synonymous substitutions at either species or ancestral nodes. We found that the transmembrane helix domains are highly conserved, whereas the intracellular loop and the C-terminus, both of which are disordered regions, were more variable among species (Fig. 6A and B). These regions experienced the highest number of non-synonymous substitutions, with the greatest extent of change observed within the Choprae clade. *D. tinwini* exhibited non-synonymous changes solely within the intracellular loop and the C-terminus.

**Fig. 6:**
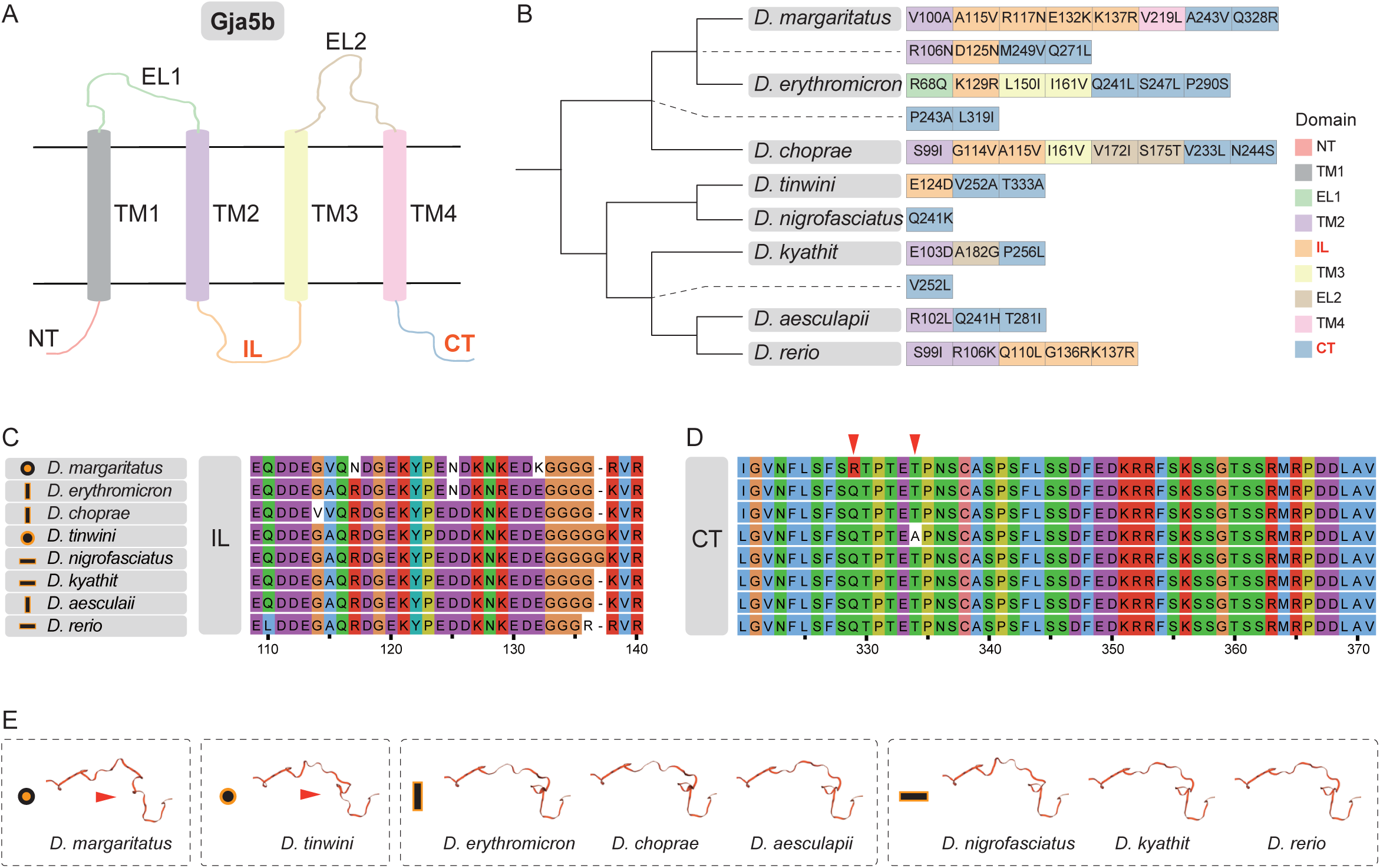
Divergence in Gja5b proteins in *Danio*. (A) Structure and domains of Gja5b protein. The two horizontal lines demarcate the predicted cell membrane region, with TM1 - TM4 representing transmembrane domains (TM). (B) Amino acid substitutions in each species or node. The box colors indicate domains where amino acid substitutions were located. (C) Multiple sequence alignment of the intracellular (IL) domain amino acids. (D) Multiple sequence alignment of the C-terminal (CT) domain amino acids. Red triangles mark specific amino acid substitution sites found in the CT domain of two spotted-pattern species (*D. margaritatus* and *D. tinwini*). (E) Protein structure prediction of the C-terminus of Gja5b. Red triangles mark unique features predicted for the CT domain in two spotted-pattern species (*D. margaritatus* and *D. tinwini*). To highlight the structural similarities within pigment pattern categories, the predicted structures are grouped and arranged by pattern (black body with yellow spots, yellow body with black spots, vertical bars, horizontal stripes), as delineated by dashed boxes. z

The intracellular loop of Gja5b exhibited numerous clustered amino acid variations (Fig. 6C). *D. margaritatus* and *D. tinwini* each carry one unique amino acid residue in the C-terminus. Structural modeling based on protein sequences suggests that this domain may adopt a similar folded conformation in both species, whereas it appears to be structurally conserved across other species (Fig. 6D and E).

These considerations coupled with the results of the complementation tests (Fig. 5) thus raise the intriguing possibility that Gja5b structure and function has independently converged in *D. tinwini* and *D. margaritatus*, perhaps facilitating the evolution of a different spotted pattern in each species. Indeed, changes in *gja5b* as well as *gja4* may contribute to differences more broadly between the Choprae clade and *D. rerio*, given additional non-complementation phenotypes involving these loci (Fig. 5A) and changes in average dN/dS ratios that suggest dynamic changes in *gja4* in the common ancestor of the Choprae clade (Table S8). Given that the common ancestor of *Danio* likely had horizontal stripes presumably dependent on gap junctions between and amongst melanophores and xanthophores involving Gja4 and Gja5b as in *D. rerio*^15,16^, diminished gap junction communication between pigment cells may have been a pre-requisite for the evolution of vertical bars and spots in the Choprae clade.

Because the interspecific complementation tests suggested a divergence in *igsf11* function between *D. rerio* and *D. margaritatus* as well as *D. erythromicron* (Fig. 5a), we also assessed sequence evolution at this locus. We identified positive selection of *igsf11* at the ancestral node of the Choprae clade (Table S8) and found that *D. margaritatus* uniquely exhibits four amino acid substitutions and ten consecutive amino acid deletions at the C-terminal region of the protein relative to all other *Danio* species (Fig. S25). The *D. margaritatus*-specific Ser390Ala amino acid substitution aligns with the polar residues (Ser375) of the mouse Igsf11 protein, which may affect protein interaction functions. These findings raise the hypothesis that functional divergence in *igsf11* has contributed to patterning differences between *D. rerio* and *D. margaritatus*. The new genome sequences we have produced now make testing this hypothesis possible.

### 8. Divergence in gene expression by regulatory evolution relative to pigment pattern diversification

Beyond protein coding sequences, the availability of assembled and annotated genomes allowed us to screen for alterations in gene expression of potential functional significance between species. To identify genes with expression differences attributable to cis-regulatory alterations, we again used hybrids to generate a common trans-regulatory environment for *D. rerio* and each of four other species, *D. aesculapii*, *D. kyathit*, and *D. nigrofasciatus*, a close relative of *D. tinwini*, as well as the more distantly related *D. albolineatus*, species chosen because earlier aspects of their development and differences from *D. rerio* have been studied in other contexts^17,18,46–48^.

To identify genes exhibiting cis-regulatory variation of potential relevance to pigmentation, we used in these hybrids zebrafish transgenic for fluorescent reporters of melanophores (*tyrp1b:mem-mCherry*), xanthophores (*aox5:mem-EGFP*)^49^ or iridophores (*pnp4a:mem-mCherry*)^18^ and we enriched for pigment cells by fluorescence activated cell sorting. To distinguish alleles in hybrids, we mapped RNA-seq libraries to both parental genomes and retained reads that had a single best match, separating the mapped reads into one library for each species in a given hybrid sample (Fig. 7A). We then remapped reads to the *D. rerio* genome to ensure a consistent set of gene annotations across species. The reporter genes were upregulated in the expected cell types, verifying enrichment of hybrid cells (Fig. S26A). To validate the sort for skin, we used *col1a2*, expressed by non-pigment skin cells involved in pattern formation^50,51^. IGV traces showed that distinct alleles with SNPs matching the two parental genomes were assigned to the appropriate species (Fig. S26B). Across all libraries, many genes displayed a slight bias for the *D. rerio* allele, which may reflect some non-*D. rerio* reads failing to map to the *D. rerio* genome, while reads from mitochondrial genes mapped almost exclusively to the maternal *D. rerio* genome as anticipated (Fig. S26C). Transcriptome-wide analysis of normalized expression values separated samples according to cell type and species (Fig. 7B). The separation of the *D. rerio* libraries from the other species in the first two principal components likely reflects the high expression of maternal *D. rerio* mitochondrial genes.

**Fig. 7:**
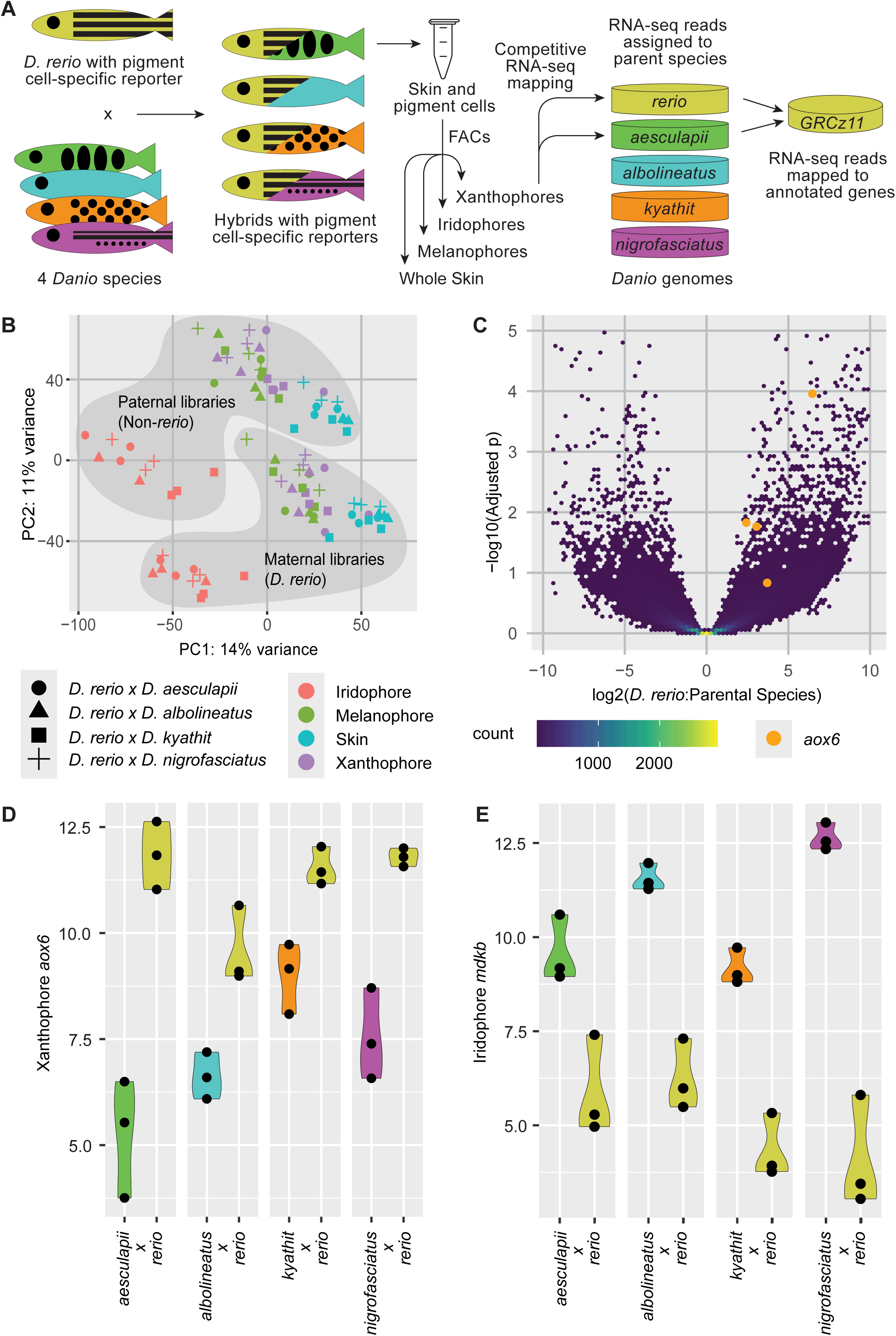
Allele-specific RNA-seq across five *Danio* species and four cell types. A) Experiment overview. B) Principal components plot of all 96 sublibraries. C) Volcano density plot showing allele-specific expression differences in xanthophores, with *aox6* highlighted in orange. D) Violin plots showing allele-specific expression of xanthophore *aox6* and E) iridophore *mdkb* (log2 normalized counts) in hybrids between *D. rerio* and *D. aesculapii, D. kyathit, D. nigrofasciatus* and *D. albolineatus*.

We used DEseq2 to identify genes with allele-specific differences between the *D. rerio* allele and the allele from the other species in each library. Across all 16 treatments (4 cell type-enrichments and 4 hybrid types) we identified 12,107 instances of allele-specific expression (adjusted *p*-value < 0.05) corresponding to 3.04 % of comparisons. To focus on loci that were recently changed in zebrafish, we identified 778 nuclear genes where the *D. rerio* allele was differentially expressed in the same direction in the same cell type across multiple species comparisons. This list was enriched for gene ontology terms related to chemokine-directed cell migration and potassium ion transport; two processes important for proper pigment pattern development (Supplemental Material 2). Of particular interest, the *D. rerio* allele of *aldehyde oxidase* 6 (*aox6*, a tandem duplicate of *aox5*, which affects yellow pigmentation), is expressed significantly higher in xanthophores relative to alleles from other species (Fig. 7C and D). This finding suggests that a *cis*-acting regulatory change has increased the expression of *aox6* in *D. rerio* and affected xanthophore pigment intensity. Conversely, the *D. rerio mdkb* allele was expressed lower relative to all other species in iridophores of hybrids (Fig. 7E), suggesting another recent *cis*-acting regulatory change in *D. rerio*. Several other genes previously associated with pigmentation are of potential interest as well (e.g., expressed in *D. rerio* at lower levels: *magoh*, *snap23.1*; at higher levels: *bloc1s6*, *lrmda*, *myg1*, *thrab*).

To further assess the utility of this dataset, we screened a small number of additional genes for expression in *D. rerio* by *in situ* hybridization during the larva-to-adult transformation. Of 34 genes examined, 26 had distinct domains of expression, with most identifying cells in presumptive epidermal or dermal compartments of the skin (Fig. S27A). In some instances, genes were clearly expressed in epidermal cells, dermal scales, or other cell types, consistent with expression at lower levels in pigment cells, or indicating “by-catch” of non-pigment cells during FACS isolation. Expression patterns of other genes in this set were suggestive of xanthophores (*asip2b*, *mc5r*) and iridophores (*mdkb*).

Indeed, comparison of *mdkb* expression between *D. rerio* and *D. albolineatus* revealed markedly greater staining in the latter species (Fig. S27C and D), consistent with allelic differences by RNA-seq (log2FC = –3.1 to –4.5, Dre < Dal) (Fig. 7E). Roles for *asip2b* have been found in cichlid stripe patterning^52^ as well as fin leucophores of *Danio rerio*^53^. The other tested loci have not been associated with pigmentation in *Danio*, but these analyses suggest a new set of high priority candidate genes that may have contributed to species-specific pigmentation, or differences in skin or scale development^50^.

## Discussion

The 14 *Danio* genome sequences presented here offer an opportunity to reconstruct the evolutionary history of genes and pathways and relate genomic changes to phenotypic shifts, epitomized here in pigment pattern evolution. The phylogenomic analyses point to a complex history of incomplete lineage sorting, introgression, and possible genomic rearrangements contributing to topological discordances between *Danio* species. Integrating this sequence information with biogeographical data provides clues about how geological events like mountain formation led to variance and secondary contact. Based on all the aforementioned analyses, we speculate that the speciation of *D. rerio*, *D. kyathit* and *D. aesculapii* was accompanied by geographical isolation and gene flow with other *Danio* species after the divergence of the most recent common ancestor (MRCA) of *D. rerio*, *D. aesculapii* and *D. kyathit* and the MRCA of *D. tinwini* and *D. nigrofasciatus*. This complex speciation history resulted in the genetic material of *D. rerio* and *D. kyathit*, originally more closely related, being now more closely related to their sympatric *Danio* species.

Beyond the complex patterns of speciation and gene flow, our demographic analyses provide insights into *Danio* population history. Closely related Danios shared similar *N_e_* trends, and the *N_e_* of *Danio* species did not decrease at the beginning of the last glaciation, except for *D. rerio*. Even during the last glacial maximum (about 26.5-19 KYA), only the *N_e_* of *D. aesculapii* and *D. erythromicron* decreased sharply, while most of the *Danio* species were not strongly affected. Our results suggested no significant correlation between the *N_e_* of *Danio* species (except *D. rerio*) and climate change during the glacial period. In contrast to the historical population dynamics, current reports indicate the beautiful *D. margaritatus* as endangered, likely caused by restructuring of their natural environment by humans.

Pigment pattern formation is a self-organizing process based on interactions between pigment cell types, as shown for example in *D. rerio*. Genes encoding transmembrane proteins regulate some of these interactions and serve as candidates for genes that diverged between species. Based on our cross-genus complementation tests in hybrids between *D. rerio* mutants and nine other *Danio* species, we demonstrated a complex genetic basis of pigment pattern diversification. In particular, these complementation tests suggest functional divergence in *igsf11* between *D. rerio* and *D. margaritatus*, but less clearly between *D. rerio* and *D. erythromicron* or *D. choprae*, the closest relatives of *D. margaritatus*. Our analyses of selection lend additional support to potential roles for *igsf11* in allowing for the evolution of the unique spotted pattern of *D. margaritatus*. Complementation tests and structural considerations also support the possibility that convergent changes in *gja5b* within *D. margaritatus* and *D. tinwini* contributed to these patterns.

Our results also suggest that *gja5b* may have functionally converged in *D. margaritatus* and *D. tinwini*, which are situated in different clades within *Danio*, and that this convergence might have mediated changes in Gja5b proteins resulting in the convergent evolution of their spotted patterns. Gap junctions containing Gja5b are critical for stripe patterning in *D. rerio* with mutants developing spots similar to the wild-type pattern of *D. tinwini*^15,16,54^. The C-terminal domain of Gja5b mediates anchoring and localization by stabilizing the gap junction plaque and facilitating interactions with regulatory proteins like ZO-1, and also modulates channel gating^55–57^; the intracellular loop domain of Gja5b may regulate channel gating and trafficking, and other interactions^58^. Functional divergence at these sites in *D. margaritatus* and *D. tinwini* could impede gap junction formation or gating among melanophores and xanthophores, as compared to the situation in *D. rerio*. Thus, it seems likely that both *igsf11* and *gja5b* have been involved in pigment pattern divergence; delineating those roles more fully will require additional mutants in multiple species associated with functional analyses, experiments facilitated by resources reported here.

Our results, together with previous research^8^, provide evidence that the ancestral pigmentation pattern of the *Danio* genus is likely horizontal stripes, while vertical bars and spotted patterns recurrently evolved in different *Danio* lineages over the course of evolution. Our findings suggest a model in which *D. rerio*, *D. kyathit*, and *D. nigrofasciatus* retained ancestral striped patterns, whereas structural variants in the C-terminus of Gja5b impaired normal intercellular interactions between melanophores and xanthophores, disrupting horizontal stripes and leading to spots in *D. tinwini*. Mutations in *kcnj13* and associated signaling interrupted horizontal striping by affecting homotypic (between the same cell type) and heterotypic (between different cell types) interactions, allowing for vertical bars and spots. The common ancestor of the Choprae clade had already evolved vertical bars. Here, amino acid changes in the intracellular loop of Gja5b may have led to reduced function and discontinuous stripes, with divergence in *kcnj13* of *D. choprae* contributing to reduced contrast between dark bars and light intervening regions, similar to *D. aesculapii*. Finally, divergence of both *gja5b* and *igsf11* led to the formation of small irregular spots in *D. margaritatus*. Additional functional analyses will allow testing roles hypothesized for these genes, as well as other genes identified here, as having cis-regulatory variation in expression among species, or that may be identified through future unbiased mutation screens. The genome sequences produced in our study provide essential resources for such endeavors and highlight the *Danio* genus as a model for studying pigment pattern formation and evolution, as well as the many other traits that vary within the genus, like size, various morphologies, ageing, and other traits. The integration of genomic and genetic analyses provides exciting possibilities to investigate the molecular and complex genetic basis of morphological diversification in a vertebrate system.

## Methods

### 1. Genome Assembly and Annotation

#### 1.1 Genome assembly

The assembly of *D. aesculapii* and *D. kyathit* were based on PacBio data, 10X Genomics Chromium data, BioNano data and Dovetail Hi-C data. The original assembly of PacBio long reads was performed using Falcon-unzip^59^. Then the assembly was purged from alternative haplotigs using purge_haplotigs^60^. Next, scaffolding was performed using three datasets: 10X based scaffolding with scaff10x, BioNano hybrid-scaffolding with Solve, Hi-C based scaffolding with SALSA2^61^. PacBio CLR reads were mapped to the assembly using PBALIGN^62^ and one round polishing was implemented using Arrow^59^. 10X reads were mapped to the assembly using Longranger^63^ and another two round of polishing were performed by Freebayes^64^. The assembly of *D. margaritatus*, *D. erythromicron* and *D. nigrofasciatus* were performed using Platanus^65^. Next, gaps were closed using paired-end information to retrieve read pairs in which one end mapped to a unique contig and the other was located in the gap region by GapCloser^66^. The genome assembly of *D. rerio* (AB), *D. rerio* (NA), and *D. rerio* (CB) was scaffolded to the chromosome level using the Sanger AB Tübingen map (SATmap) data^67^. For *D. tinwini*, *D. jaintianensis*, *D. albolineatus*, *D. choprae*, and *D. cerebrum*, the assembly was performed based on 10X Genomics Chromium data. The original assembly of 10X reads was performed using Supernova 2.0.1^68^, and haplotig identification with Purge Haplotigs^69^. The genome of *D. dracula* was also assembled using PacBio long reads data. All the assemblies were finally analysed and manually improved using gEVAL^70^.

#### 1.2 Repeat annotation

Firstly, LTR_FINDER^71^ and RepeatModeler (version 1.0.4) (Smit, Hubley, & Green, 2015) were used to find repeats. Next, RepeatMasker (version 4.0.5)^72^ was used (with the parameters: -nolow -no_is -norna -parallel 1) to search for known and novel transposable elements (TEs) by mapping sequences against the *de novo* repeat library and Repbase TE library (version 16.02)^73^. Subsequently, tandem repeats were annotated using Tandem Repeat Finder ^74^ (version 4.07b; with following settings: 2 7 7 80 10 50 2000 -d -h). In addition, we used the RepeatProteinMask software (version open-4.0.6, with parameters: -no LowSimple -p value 0.0001) to identify TE-relevant proteins (Fig. S1, S2).

#### 1.3 Gene annotation

For de novo gene prediction, we utilized SNAP (version 2006-07-28), GENSCAN (version 1.0)^75^, GlimmerHMM (version 3.0.3)^76^, and AUGUSTUS (version 2.5.5)^77^ to analyze the repeat-masked genome. For homology-based predictions, the protein sequences of *Danio rerio* (GRCz11) were used as templates for homology-based gene prediction for all of the newly assembled *Danio* genomes. First, we aligned protein sequences of the reference gene set to each genome by TBLASTN^78^ with an E-value cut-off of 1E-5. We filtered the candidate loci for which the homologous block length was shorter than 30 % of the length of the query protein. Then, we extracted genomic sequences of candidate gene loci, including the intronic regions and 2,000 bp upstream/downstream sequences. The retrieved sequences were subjected to more precise alignment performed with GeneWise (version 2.2.0)^79^. The output of GeneWise includes the predicted gene models in the genome. Then, we translated the predicted coding regions into protein sequences. We filtered out predicted proteins that had a length of < 30 amino acids (aa) or percent identity of < 25 %. EVidenceModeler software (EVM, version 1.1.1)^80^ was used to integrate genes predicted by the homology and de novo approaches and generate a consensus gene set. Short-length (< 50 aa) and prematurely terminating genes were removed from the consensus gene set, and the final gene set was produced (Fig. S1, S3).

### 2. Identification of genomic variation

#### 2.1 SNPs and Indels analysis

Reads were aligned to the *Danio rerio* genomic sequence version 11 (Genome Reference Consortium zebrafish build 11, GRCz11) using the bwa - mem v.0.7.17 algorithm with default options^81^. Next, we converted the aligned results to bam files using SAMtools (version 1.3.1)^82^. We sorted the bam files with commands: samtools sort and duplicate reads were marked using the GATK v4.1.0.0^83^ MarkDuplicates tool. Then, all marked bam files were validated by GATK ValidateSamFiles tool for calling SNPs and Indels.

#### 2.2 Variant calling, filtering, and genotype refinement

Briefly, SNP and short intel variants against the GRCz11 reference were called with GATK HaplotypeCaller. Variant filtering was then performed on the GATK VariantFiltration using hard filters based on variant quality by unfiltered depth (QD), root mean square mapping quality (MQ), u-based z-approximation from the Rank Sum Test for mapping qualities (MQRankSum), u-based z-approximation from the Rank Sum Test for site position within reads (ReadsPosRankSum), Fisher Stand (FS).

#### 2.3 Presence-absence variation analysis

To explore the presence/absence variations (PAVs) for all available assemblies, we compared the reference genome (GRCz11) with the new genome assemblies of other *Danio* species. We searched PAV sequences that were present in the reference genome but absent from other genome assemblies using scanPAV^84^ and generated a list of one-to-one correspondences.

### 3. Evolutionary analysis

#### 3.1 Phylogeny

##### 3.1.1 Whole genome tree

Whole genome alignments (WGAs) are critical for comparative analyses, and we generated multiple genome alignments for all 15 *Danio* and *Danionella* genomes as well as four outgroup fishes. First, pairwise alignments for each pair of genomes were produced by the LAST version 982 package^85^, using the *Danio rerio* (GRCz11)^86^ genome as reference. Each genome was aligned to the reference using the “lastal” command with the parameter -E0.05. Then, we used the “maf-swap” command to change the order of sequences in the MAF-format alignments and obtained the best pairwise aligned blocks. Lastly, we used MULTIZ version 11.2^87^ to merge the pairwise alignments into multiple genome alignments. Approximately 153 Mb of conserved syntenic sequences shared by all the *Danio*, *Danionella* and outgroup species were obtained in the final alignment. WGAs of 19 fish genomes were used to construct a phylogenetic tree rooted by the *Lepisosteus oculatus* genome sequence. Syntenic blocks were concatenated using bespoke Python scripts, and a FASTA-formatted alignment file was then generated. We used IQtree (version 1.7.6)^88^ to estimate the model (using the ModelFinder^89^ function in IQtree), the tree, and 100 standard bootstraps (command: iqtree -s <alignment> -m MFP -b 100). Finally, we obtained a maximum likelihood (ML) tree with bootstrap supports on each node.

##### 3.1.2 Single copy orthologous gene trees

The genome and annotation data for *Lepisosteus oculatus*, *Oryzias latipes*, *Ictalurus punctatus*, *Astyanax mexicanus* and *Danio rerio* were downloaded from Ensembl (release 92). The longest predicted translation product was chosen to represent each gene, and gene models with an open reading frame <150 bp in the genomes were removed. Next, these protein sets were pooled, and self-to-self BLASTP was conducted for all of the aforementioned protein sequences with an E-value of 1e–5. Hits with identity values less than 30 % and coverage less than 30 % were removed. Then, based on the filtered BLASTP results, orthologous groups were constructed by ORTHOMCL v2.0.9^90^. Phylogenetic tree inference was conducted based on 1&2 phase sites of single-copy gene families in series by the Mrbayes^91^ software with GTR+Gamma model and set mcmc ngen=100000 printfreq=100 samplefreq=100 nchains=4 savebrlens=yes. We also used OrthoFinder (version 2.3.8)^92^ software to identify single copy families and construct the phylogenetic tree in multiple processes, which showed the same phylogenetic structure. Divergence time estimation was performed using the MCMCTREE in the PAML4.7 package^93^.

##### 3.1.3 SNP trees

We converted from MAF to FASTA using a bespoke Perl script and removed all gap sites using trimAl. Then, we converted FASTA to VCF using snp-sites^94^, and finally filtered the snps-site output using VCFtools (version 0.1.16)^95^ with parameters “--min-alleles 2 --max-alleles 2 --thin 100”. Sites were included if they were present in all species (i.e., no missing data or gaps) as a single copy, bi-allelic SNPs. This process resulted in a set of 51,633 SNPs.

We applied a multispecies coalescent method that attempted to reconstruct the species tree based on the SNPs dataset. We used SVDquartets^96^ as implemented in PAUP* (v4.0a, build 166)^97^. We prepared the data into the NEXUS format, using Python script. Then we ran SVDquartets in PAUP* setting outgroup to *L. oculatus* and then executing evalQuartets=all with 100 standard bootstraps.

Next, we also used SNAPP^98^ as implemented in BEAST version 2.6.2^99^ to construct a phylogenetic tree based on the SNP dataset. The “forward” and “backward” mutation rate parameters u and v were calculated directly from the data by SNAPP using the “Calc mutation” rates option. The default value 10 was used for the “Coalescent rate” parameter and the value of the parameter was sampled (estimated in the Markov chain Monte Carlo (MCMC) chain). The prior for the ancestral population sizes was chosen to be a relatively broad gamma distribution with default parameters. We ran a 1,000,000,000-iteration chain with sampling every 50,000 iterations. Due to time limitations, the program was terminated early, and finally 48,600,000 iterations were carried out.

##### 3.1.4 Window-based gene trees

To investigate the phylogenetic discordance across genomic regions, we segmented the WGA sequences in 10-kbp non overlapping windows. After excluding those windows that included repeat sequences and sequence sizes less than 150 bp, a total of 9101windows remained. We then constructed the window-based gene tree (WGT) for each window using IQtree with 100 standard bootstraps (command: iqtree -s <align.fa> -o <outgroup> -b 100 -m MFP). Subsequently, we applied ASTRAL to reconstruct the species tree from WGTs using the default parameters (for the detail, see “Visualization of gene-tree discordance” below).

##### 3.1.5 Visualization of gene-tree discordance

We filtered WGTs so that they included (*Danio rerio* Tübingen, *Danio rerio* AB, *Danio rerio* NA, *Danio rerio* CB) in a monophyletic clade. Then, we applied ASTRAL to reconstruct the species tree based on WGTs using the default parameters, respectively. We visualized phylogenetic conflicts by superimposing WGTs in a DensiTree (version 2.2.5)^33^ plot. Prior to superimposition, we rendered the trees ultrametric using the R package Phybase (v2.0)^100^. Furthermore, we used DiscoVista (Discordance Visualization Tool, version 1.0)^34^ to analyze gene-tree compatibilities with parameters: “-m 5”, using WGTs (Quartet frequencies of the internal branches in the species tree were calculated using ASTRAL).

#### 3.2 Gene flow

##### 3.2.1 D-Statistics

We filtered SNP sites (for details of SNP calling, see “SNP and Indels analysis”) using BCFtools version 1.8 with parameters “bcftools view -e ‘AC==0 || AC==AN || F_MISSING > 0.2’ -m2 -M2”. This process resulted in a set of 93043 SNPs. We calculated the D-statistic for all possible trios of Rerio clade species (P1, P2, P3) without assuming a known species tree topology, using Dsuite software^38^ (command: ./Build/Dsuite Dtrios <SNPs> <samples>) with *D. albolineatus* as outgroup. To get a better overview of introgression patterns supported by D-statistics, we used ggplot2 version 3.3.2 to visualize these in the form of a bubble chart in which a circle in the bubble chart indicates the most significant D-statistic found with P2 and P3 (*p*<0.05).

##### 3.2.2 D_FOIL_ statistics

Pease and Hahn^101^ proposed a five-taxon test to distinguish ILS from gene flow (the D_FOIL_ statistics). D_FOIL_ analyses assume a symmetrical five-taxon topology: (((P1, P2), (P3, P4)), O), which can determine the direction of any detected introgression phylogeny. We used *L. oculatus* as outgroup, and (*D. tinwini*, *D. nigrofasciatus*), (*D. erythromicron*, *D. margaritatus*) or (*D. choprae*, *D. jaintianensis*) as (P3, P4) with other *Danio* species as (P1, P2) to calculate the D_FOIL_ statistics for all possible four fitted topologies based on WGA sequences (excluding repeat sites) using D_FOIL_^39^. We also used *D. albolineatus* as an outgroup to detect geneflow between species for the *D. rerio* subgroup. We tested five fitted topologies: (1) (((*D. rerio*, *D. aesculapii*), (*D. kyathit*, *D. tinwini*)), *D. albolineatus*); (2) (((*D. rerio*, *D.aesculapii*),(*D. kyathit*, *D. nigrofasciatus*)), *D. albolineatus*); (3) (((*D. rerio*, *D. aesculapii*), (*D. tinwini*, *D. nigrofasciatus*)), *D. albolineatus*); (4)(((*D. rerio*, *D. kyathit*), (*D. tinwini*, *D. nigrofasciatus*)), *D. albolineatus*); (5) (((*D. aesculapii*, *D. kyathit*), (*D. tinwini*, *D. nigrofasciatus*)), *D. albolineatus*). The 10,175 10-kbp windows (excluding repeat sites and sequence lengths no less than 150 bp), were used to analyze gene flow using D_FOIL_.

##### 3.2.3 Divergence time calibration

Divergence time estimation was performed using the MCMCTREE in PAML4.7 package^93^. The upper and lower limit of the divergence time found on the TIMETREE website (http://www.timetree.org/) was used as the calibration time: *D. rerio* - *D. erythromicron* (27 - 32 mya); *D. rerio* - *Danionella cerebrum* (36 - 68 mya); *A. mexicanus* - *I. punctatus* (109 - 157 mya); *D. rerio* - *O. latipes* (206 - 252 mya); *D. rerio* - *L. oculatus* (295 - 334 mya). Then, we used MCMCTREE in the PAML4.7 package to estimate divergence times based on three different topological evolution trees (1 and 2 codon sites BI tree, 4d sites BI tree and SNP tree constructed by SNAPP) with calibration time and the corresponding multiple sequence alignment files. We first calculated the substitution rate using baseml in PAML, then set usedata=3 and clock=2 to generate out.BV using MCMCTREE, and finally set usedata=2 and clock=2 to calculate the divergence time using MCMCTREE.

#### 3.3 Identification of positively selected genes (PSGs)

We used a conserved genome synteny methodology to establish a high-confidence orthologous gene set that included *Ictalurus punctatus*, *Astyanax mexicanus*, *Oryzias latipes*, *Lepisosteus oculatus*, and these *Danio* species. Briefly, pairwise WGAs were constructed for relevant genomes using LAST, with the GRCz11 *D. rerio* sequence serving as the reference genome. To minimize the effect of annotation, sequencing, and assembly errors, pseudogenes, non-orthologous alignments, and non-conserved gene structures on subsequent evolutionary rate analyses, a series of rigorous filtering criteria was adopted: (1) genes mapped to the reference genome via a single chain of sequence alignments including at least 80 % of the gene’s coding sequence (CDS), and met the alignment length/score thresholds required for inclusion in the MULTIZ alignments; (2) frame-shift indels in CDSs were prohibited; (3) CDSs with premature stop codons were excluded and (4) genes with Ks values (synonymous substitutions per synonymous site) between each species and GRCz11 larger than two were excluded.

Based on the filtered orthologous gene set, we estimated the lineage-specific evolutionary rate for each branch. The Codeml program in the PAML package (version 4.8) with the free-ratio model (model=1) was run for each ortholog. Positive selection signals on genes along specific lineages were detected using the optimized branch-site model following the author’s recommendation^102^. A likelihood ratio test (LRT) was conducted to compare a model that allowed sites to be under positive selection on the foreground branch with the null model in which sites could evolve either neutrally or under purifying selection. The *p*-values were computed based on Chi-square statistics, and genes with *p*-value less than 0.05 were treated as candidates that underwent positive selection.

In order to identify candidate genes contributing to pigment pattern variations, we defined a set of 253 single-copy genes that overlapped with a curated list of 650 pigmentation-associated genes^103^, and calculated their dN/dS ratios for each branch under the two-ratio branch model (model 2) of PAML. Background dN/dS ratios were calculated under the one-ratio branch model (model=1) of PAML. The distribution densities of the dN/dS values were shown (Fig. S19, S21, S23). We further identified genes with elevated dN/dS ratios in each branch by comparing with background values of dN/dS under Student’s t test of *p*-value < 0.01.

#### 3.4 Demographic history reconstruction

We inferred the demographic history for Danio species by applying the pairwise sequentially Markovian coalescent model (PSMC)^82^. We used aligned reads and consensus sequences to conduct the PSMC (version 0.6.5-r67) analysis. Population size histories were inferred by PSMC (with the parameters: psmc -N25 -t15 -r5 -p “4+25*2+4+6”). The generation time (g) of fish in the *Danio* genus was obtained from previous studies^104^. Per-year mutation rates were estimated by r8s. The per-generation mutation rate is estimated by multiplying the per-year mutation rates by the generation time.

### 4. Genus-wide one-way complementation tests

#### 4.1 Fish husbandry

Zebrafish, *D. rerio*, were maintained as previously described^105^. If not newly generated, the following lines were used for experiments: Wild-type *D. rerio* Tübingen strain, *gja4^t37ui^*^15^ and *igsf11^t35ui^* ^9^. Wild-type Tübingen strains of *D. aesculapii*, *D. nigrofasciatus* and *D. albolineatus* were maintained identical to *D. rerio*. For the other *Danio* species, *D. kyathit*, *D. tinwini*, *D. choprae*, *D. margaritatus*, *D. erythromicron* and *D. dangila*, individual pair matings were not successful. Therefore, those species of fish were kept in groups in tanks containing boxes lightly covered with Java moss (*Taxiphyllum barbieri*), which resulted in sporadic matings and allowed us to collect fertilized eggs.

Interspecific hybrids were either obtained by natural mating or by in vitro fertilization. Hemizygous mutant hybrids were identified by PCR and sequence analysis using zebrafish-specific primer pairs for *gja4* (Tü838_for: 5’-TGCCTCTAGGAACATGATTGGG-3’ and Tü975_rev: 5’-GGTCATCTTCGTCTCAACTCCG-3’), *gja5b* (MP335_for: 5’-CAGGCTCCTCTGAATAGGCA-3’ and MP336_rev: 5’-GTGTAGACACGAACACGATCTG-3’) and *igsf11* (Tü1449_for: 5’-TCATCTACCAGAGTGGTCAG-3’ and Tü1450_rev: 5’-CCTAAACTTTTGCAGCACAG-3’). Phenotypic variability observed between hybrids of different genotypes could be caused by the influence of novel genetic backgrounds from specific species pairs rather than by a functional divergence of the tested genes. Initially, we used the established *D. rerio* strain *leo^t1^*, carrying a nonsense mutation in *gja5b*, for the complementation tests. However, we frequently observed variable phenotypes unrelated to the genotypes of the resulting hybrids. Therefore, we repeated these experiments with a new CRISPR/Cas9-generated loss-of-function allele, *gja5b^t21mp^*, induced in the same genetic wild-type background as our other mutants. The genetic background in which this allele was generated reduced the observed phenotypic variability in hybrids. Consequently, we used CRISPR alleles for all other interspecific complementation tests.

All species were staged according to the normal table of *D. rerio* development^106^. All animal experiments were performed in accordance with the rules of the State of Baden-Württemberg, Germany, and approved by the Regierungspräsidium Tübingen. Individuals used for genome sequencing were maintained and handled in accordance with protocols approved by the institutional animal care and use committee (IACUC) at the University of Washington, Seattle.

#### 4.2 CRISPR/Cas9-mediated knock-out

The CRISPR/Cas9 system was applied to generate loss-of-function mutations in *gja5b* as previously described^107^. Briefly, the oligonucleotides Tü1037_for (5’-TAGGCTGCTGAATCCTCGTGGG-3’) and Tü1038_rev (5’-AAACCCCACGAGGATTCAGCAG-3’) were cloned into pDR274 to generate the sgRNA vector. sgRNAs were transcribed from the linearised vector using the MEGAscript T7 Transcription Kit (Invitrogen). The sgRNA was injected as ribonucleoprotein complex with Cas9 protein into one-cell stage embryos. The efficiency of indel generation was tested on eight larvae at 1 dpf by PCR using *gja5b*-specific the primer pairs MP335_for and MP336_rev (see 4.1 above) and by sequence analysis as described previously^108^. The remaining larvae were raised to adulthood. Mature F0 fish carrying indels were outcrossed. Recessive loss-of-function alleles in heterozygous F1 fish (c.16_25delCTGCTGGGGA, p.Lys6ThrfsX2) were selected to establish the homozygous mutant line *gja5b^t21mp^* developing the characteristic mutant phenotype similar to *gja5b^t1^* ^109,110^.

#### 4.3 Image acquisition and processing

Anesthesia of adult fish was performed as described previously^111^. Bright field images of adult fish were obtained using a Canon 5D Mk II camera. Fish with different pigment patterns vary considerably in contrast, thus requiring different settings for aperture and exposure time, which can result in slightly different color representations in the pictures. Images were processed using Adobe Photoshop and Adobe Illustrator CS6.

### 5. Hybrid RNA-sequencing analysis and in situ hybridization

#### 5.1 Experimental Design and Sample Collection

To measure cis-regulatory changes in gene expression across Danio species, we generated interspecies hybrids by crossing males from four species: *D. aesculapii*, *D. kyathit*, *D. albolineatus*, and *D. nigrofasciatus* to female *D. rerio* carrying cell-type specific fluorescent reporters for melanophores (*tyrp1b:mem-mCherry*), xanthophores (*aox5:mem-EGFP*)^49^ or iridophores (*pnp4a:mem-mCherry*)^18^. We dissociated and FACS sorted adult hybrid skin to enrich for iridophores, melanophores, or xanthophores based on reporter expression, yielding 36 cell-enriched biological samples (3 cell types x 3 replicates x 4 paternal species) and 12 whole skin control samples (3 replicates x 4 paternal species).

#### 5.2 RNA-seq and Read Processing

Full-length RNA-seq libraries were prepared using SMART-Seq2 (Takara Bio) and sequenced on an Illumina NextSeq with PE 150 reads. RNA-seq in hybrids requires discerning species of origin as well as gene of origin for each read. To avoid any issues with differing annotation quality, we used a two-step mapping strategy. First, to distinguish parental alleles in hybrid samples, we mapped each of the 48 samples against combined references containing both parental genomes. The best mapping location of each read was used to assign species of origin and split each of the 48 sequencing libraries into two species-specific sublibraries (96 in total). We then remapped each sublibrary to the *D. rerio* genome to utilize the available high quality GRCz11 gene annotations. For both steps, reads were mapped with BBMap (v38.57)^112^ with parameters: k=13, maxindel=100000, minid=0.76, ambiguous=best. The “ambiguous” parameter addresses any reads that map equally well to two locations in the combined, two-species references. The very low minimum identity accounts for sequence divergence between species when remapping to GRCz11.

#### 5.3 Differential Expression Analysis

Gene expression was quantified using featureCounts (Rsubread v2.18)^113^. To mitigate the possibility of mapping bias toward the *D. rerio* reference in regions with poor sequence conservation, we excluded reads mapping to 3’ and 5’ untranslated regions. Statistical analysis was performed using DESeq2 (v1.44.0)^114^, filtering genes with less than 1 normalized count per sample (retaining 24,672 of 29,005 genes). Allele-specific expression was tested based on species of origin to compare *D. rerio* versus non-*D. rerio* alleles within each hybrid background for each cell type enrichment. Differential expression was assessed using default parameters: Wald test with adjusted *p*-values using Benjamini-Hochberg correction and FDR < 0.05. Gene set enrichment analysis of GOTERM_BP_DIRECT terms was performed using DAVID (Knowledgebase v2023q4)^115^ with default parameters on the 778 genes that showed allele-specific differential expression in the same direction and cell type across multiple species comparisons, indicating *D. rerio*-specific *cis*-regulatory changes.

#### 5.4 In situ hybridization

*In situ* hybridization and imaging was performed as previously described^116^, with larvae sampled at standard lengths ranging from 6.2–8.5 mm (approx. 14 to 28 days post fertilization) and images captured on Zeiss Axiocam cameras.

## Acknowledgements

We thank Roberta Occhinegro for excellent technical assistance and Lauren Saunders for help with hybrid RNA-Seq library preparation. This work was supported by an ERC Advanced Grant “DanioPattern” (694289), the Max Planck Society, and NIH R35 GM222471 to DMP, and NIH R35 GM139635 to JHP. The generation of assemblies and analyses were enabled by Wellcome through core funding of the Wellcome Sanger Institute (098051 and 206194). We thank Richard Durbin for his invaluable advice and support throughout the project.

## Author contributions

Jianguo Lu and Kerstin Howe conceptualized and supervised the study, performed genome assembly, and revised the manuscript. Shane McCarthy, Jonathan Wood and Joana Collins performed genome assembly and assembly curation. Marco Podobnik conducted experiments and participated in manuscript writing and editing. Junrou Huang performed bioinformatics analysis, contributed to methodology development, and participated in manuscript preparation. Braedan M. McCluskey collected biological material for sequencing, carried out bioinformatics analysis and assisted in manuscript writing. James Torrance, Ying Sims, Dong Gao, Jing Huang and Jia Liu conducted bioinformatics analyses, with Jing Huang additionally contributing to manuscript writing. David Parichy and Uwe Irion conducted experiments and provided critical manuscript revisions. Wenyu Fang and Peilin Huang selected and analyzed candidate genes. Chunlei Ma generated geographical mapping data. Jian Liu provided overall oversight of the bioinformatics analyses. J.H. Postlethwait provided overall scientific oversight, critical guidance on the research direction, and final approval of the manuscript.

