## Supplementary Material 1 for "Genomic and genetic insights into speciation and pigment pattern diversification in *Danio* fishes"

^1^School of Marine Sciences, Sun Yat-sen University, Zhuhai, China. ^2^Southern Marine Science and Engineering Guangdong Laboratory (Zhuhai), Zhuhai, China. ^3^Max Planck Institute for Biology, Max-Planck-Ring 5, 72076, Tübingen, Germany. ^4^Present address: Australian Regenerative Medicine Institute, Level 2, 15 Innovation Walk, Monash University, Wellington Road, Clayton VIC 3800, Australia. ^5^Department of Biology and Department of Cell Biology, University of Virginia, Charlottesville, Virginia, United States of America. ^6^Present address: Minnesota Supercomputing Institute, University of Minnesota, Minneapolis, Minnesota, United States of America. ^7^Wellcome Sanger Institute, Cambridge, UK. ^8^State Key Laboratory of Medicinal Chemical Biology, College of Computer Science, Nankai University. ^9^Institute of Neuroscience, University of Oregon, Eugene, OR 97403-1254.

*joint first authors, **corresponding authors

**SM Text 1. Species information**

The final collection of Danioninae genomes used in this project contained 15 species or sub-species (strains), including 14 newly assembled genomes and the published reference genome of *Danio rerio*^1^. The genomic resources used in this project had been released in advance. Tissue samples were collected from Wellcome Sanger Institute. All samples were collected legally and in accordance with the policy of the Animal Care and Use ethics of each institution.

**SM Text 2. Genome assembly benchmarking,** **annotation and variation calling**

**Evaluation of genome assemblies**

We evaluated the genome assembly quality for each species using different strategies (described below), and the best assembly was retained as the final genome assembly for downstream analyses.

***C-value***

The *C-value* is defined as the total amount of DNA contained within a single set of chromosomes. The *C-value* can be easily converted into base pairs of genome size, using 1 pg = 978 Mb. The sizes of the genome assemblies were highly consistent with the C-values.

***K-mer* analysis**

The illumine short reads from the 250 bp and 500bp libraries were used for *k*-mer frequency analyses of genomes. For a diploid genome without severe repetitive elements and heterozygosity, the relationship between *k*-mer frequency and sequence depth follows a Poisson distribution. The frequency of each *k*-mer can be calculated from the genome sequence reads. A *k*-mer is an artificial sequence division where the sequencing reads are divided iteratively into pieces of *k* bases. As the length of each *k*-mer is *k* bp, a read with L bases contains (L – *k* + 1) *k*-mers. *k*-mer analysis can also be used to calculate a genome size (Gbp) by the formula G= *k*_num/ *k*_depth, where *k*_num is the total number of *k*-mers, and *k*_depth is the average depth of the *k*-mers.

**Evaluation of genome completeness with BUSCO**

BUSCO (version 3.0.2)^2^ was used to assess the genome completeness by estimating the percentage of expected single copy conserved orthologs captured in our assemblies, referring to the actinopterygii_odb9 database. 14 genomes with scaffolded assemblies were evaluated.

**Whole genome synteny**

We obtained the synteny blocks of *Danio aesculapii* and *Danio kyathit* by aligning to the reference genome (GRCz11) using Minimap2^3^. Each genome was aligned to the reference genome using the minimap2 with the parameter: -t 12, -X, -x asm5. Then, we filter these aligned results and obtained the best pairwise aligned blocks. The filter standard including: block length ≥ 10 kbp and score = 0.

**Genome annotation**

**Repeat annotation**

Transposable elements in the Danio genome were identified by a combination of homology-based and de novo approaches. Tandem repeats were identified using Tandem Repeat Finder^4^. Interspersed repeats were characterized by homolog-based identification using RepeatMasker open-4.0.3^5^ and the repeat database, Repbase^6^. Repeated proteins were identified using RepeatProteinMask and the transposable elements protein database. De novo identified interspersed repeats were annotated using RepeatModeler^7^, and LTR _FINDER^8^ was used to identify the LTRs; these results were used to generate the de novo repeat libraries, and then RepeatMasker was run once more against the de novo libraries. All repeats identified in this manner were included in the total count of interspersed repeats.

**Gene annotation**

The protein-coding genes were annotated following the use of a combination of homolog gene prediction and *de novo* gene prediction tools. For homolog gene prediction, the protein sequences from *Danio rerio*, *Oryzias latipes*, *Takifugu rubripes*, *Tetraodon nigroviridis*, and *Gasterosteus aculeatus* were mapped to the genome using tBLASTn^9^. Exonerate^10^ was used to predict the gene model based on the alignment results. *De novo* gene prediction was performed using GENSCAN^11^, AUGUSTUS^12^, and GLIMMERHMM^13^ based on the repeat-masked genome. For five of the genomes, *Danio aesculapii*, *Danio albolineatus*, *Danio choprae*, *Danio kyathit*, *Danio rerio* (AB)*,* which have transcriptome data, we used HISAT2^14^ to align transcriptome reads to genome and used StringTie^15^ to assembly the transcripts. Then, MAKER^16^ were applied to integrate the predicted genes. Finally, manual integration and remove the incomplete gene was performed to construct the final gene set. We searched the final gene set against the NCBI nr database, SwissProt^17^, and TrEMBL^17^ protein databases to identify gene functions. The gene motifs and domains were determined using InterProScan^18^ following analysis of public protein databases, including ProDom, PRINTS, PFAM, SMART, PANTHER and PROSITE. All genes were aligned against the KEGG pathway database^19^, and the best match for each gene was identified. The GO IDs for each gene were obtained from the corresponding InterPro entries. We also mapped the final get set to the GRCz11 proteins and identified the best hit gene symbol.

**Identification of genomic variation**

a) **SNP and Indels analysis**

Reads were aligned to the *Danio rerio* genomic sequence version 11 (Genome Reference Consortium zebrafish build 11, GRCz11) using the bwa -men v.0.7.17^20^ algorithm with default options. Next, we converted the aligned results to bam files using SAMtools (version 1.3.1)^21^. We sorted the bam files with commands: samtools sort and duplicate reads were marked using the GATK v4.1.0.0^22^ MarkDuplicates tool. Then, all marked bam files were validated by GATK ValidateSamFiles tool for calling SNPs and Indels.

Briefly, SNP and short intel variants against the GRCz11 reference were called with GATK HaplotypeCaller. Variant filtering was then performed on the GATK VariantFiltration using hard filters based on variant quality by unfiltered depth (QD), root mean square mapping quality (MQ), u-based z-approximation from the Rank Sum Test for mapping qualities (MQRankSum), u-based z-approximation from the Rank Sum Test for site position within reads (ReadsPosRankSum), Fisher Stand (FS).

**Commands/parameters used:**

**Step 1. Data pre-processing:**

bwa index -p <index_file_name> <reference>.fa

bwa mem -M <index_file> <sample_name>R1.fastq <sample_name>R2.fastq -R @RG\tID:<id_name>\tLB:<library_name>\tSM:<sample_name>\tPL:Illumina" -t 24 > <sample_name>.sam

samtools view -bS -@ 12 <sample_name>.sam > <sample_name>.bam

samtools sort -@ 12 <sample_name>.bam -o <sample_name>.sorted.bam

gatk MarkDuplicates -I <sample_name>.sorted.bam -O <sample_name>.repeatmarked.bam -M <sample_name>.repeatmarked.bam.metrics

samtools index -@ 12 <sample_name>.repeatmarked.bam

gatk ValidateSamFile -I <sample_name>.repeatmarked.bam

**Step 2. Variant calling, filtering, and genotype refinement:**

gatk HaplotypeCaller -I <sample_name>.repeatmarked.added.bam --native-pair-hmm-threads 32 -stand-call-conf 30 -O <sample_name>.g.vcf.gz -R <reference>.fa -ERC GVCF

gatk --java-options "-Xmx4g" GenotypeGVCFs -R <reference>.fa –V<sample_name>.g.vcf.gz -O <sample_name>.vcf.gz

gatk SelectVariants –V <sample_name>.vcf –R <reference>.fa –select-type SNP/INDEL –O <sample_name>.raw_snps/indels.vcf

gatk VariantFiltration –R <reference>.fa –V <sample_name>.raw_snp.vcf –filter-expression “QD < 2.0 || FS > 60.0 || MQ < 40.0 || <QRankSum < 12.5 ||ReadPosRankSum < 8.0” –filter-name “my_snp_filter” –O <sample_name>.filtered_snps.vcf

b) **Presence-absence variation analysis**

To explore the presence/absence variations (PAVs) information of all genome assemblies among these *Danio* species, we compared the DNA sequences of reference genome (GRCz11) with other genomes of *Danio* species, respectively. We searched PAV sequences present in reference genome but absent in other genome assemblies through scanPAV ^23^ and generated a list of one-to-one correspondences.

**Commands/parameters used:**

/path/scanPAV –nodes <no(Kiełbasa et al. 2011)des> -align <aligner> -scroe <sw-score> <path/reference>.fasta <path/species1>.fasta <pavs_present_in_reference>.fasta

Notes:

nodes: number of CPUs reseqested [default=30]

sw-score: smith-waterman alignemnt score [default=30]

aligner: sequence aligner: bra or small [default=30]

**SM Text 3. Evolutionary analysis**

**Phylogeny**

**Whole genome tree**

Whole genome alignments (WGAs) are critical for comparative analyses, and we generated multiple genome alignments for all the 13 *Danio* species and 6 outgroup fishes. First, pairwise alignments for each pair of genomes were produced by the LAST (version 982) package^24^, using the *Danio rerio* (GRCz11)^1^ genome as reference. Each genome was aligned to the reference using the “lastal” command with the parameter -E0.05. Then, we used the “maf-swap” command to change the order of the sequences in the MAF-format alignments and obtained the best pairwise aligned blocks. Lastly, we used MULTIZ (version 11.2)^25^ to merge the pairwise alignments into multiple genome alignments.

WGAs of 19 fish genomes were used to construct a phylogenetic tree rooted by the *Lepisosteus oculatus.* Syntenic blocks were concatenated using in-house Python scripts, and a FASTA-formatted alignment file was then generated. We used IQtree (version 1.7.6)^26^ to estimate the model (using the ModelFinder^27^ function in IQtree), the tree, and 100 standard bootstraps (command: iqtree -s <alignment> -m MFP -b 100). Finally, we obtained a maximum likelihood (ML) tree with bootstrap supports on each node.

**Single-copy orthologous gene trees**

The genome and annotation data for *Lepisosteus oculatus*, *Oryzias latipes*, *Ictalurus punctatus*, *Astyanax mexicanus* and *D. rerio* were downloaded from Ensembl (release 92). The longest predicted translation product was chosen to represent each gene, and gene models with an open reading frame <150 bp in the genomes were removed. Next, these protein sets were pooled, and self-to-self BLASTP was conducted for all of the aforementioned protein sequences with an E-value of 1e–5. Hits with identity values less than 30 % and coverage less than 30 % were removed. Then, based on the filtered BLASTP results, orthologous groups were constructed by ORTHOMCL v2.0.9^28^. Multiple sequence alignment (MSA) was made for the amino acid sequences of each gene family using MUSCLE (version 3.8.1551)^29^ and then the amino acid sequences were inversely translated back to the corresponding CDS sequences. Then, the genes were concatenated to generate a supergene sequence. Considering the composition heterogeneity of different codon positions, we further extracted 1st, 2nd codon sites and four-fold degenerate (4d) sites from the orthologous gene sequences using Perl script. Phylogenetic tree inference was conducted based on 1 codon, 1&2 codon and 4d sites of single-copy gene families using Bayesian and ML methods, respectively. Bayesian phylogenetic analysis using Mrbayes (version 3.2.6)^30^ software with GTR+Gamma model and set “mcmc ngen=100000 printfreq=100 samplefreq=100 nchains=4 savebrlens=yes”. ML phylogenetic analysis using PhyML (version 3.1) software^31^ with the command: phyml -i <input.phy> -d nt -b -4 -m HKY85 -a e -c 4 -t e.

**SNPs trees**

We converted from MAF to FASTA using in-house Perl script, and removed all gap sites using trimAl. Then, FASTA converted to VCF using snp-sites^32^, and finally filtered snps site using VCFtools (version 0.1.16)^33^ with parameters “--min-alleles 2 --max-alleles 2 --thin 100”. Sites were included if they were present in all species (i.e., no missing data or gaps) as a single copy, bi-allelic SNPs. This resulted in a set of 51,633 SNPs.

We applied multispecies coalescent method that attempt to reconstruct the species tree based on SNPs dataset. We used SVDquartets^34^ as implemented in PAUP* (v4.0a, build 166)^35^. We prepared the data into the NEXUS format, using Python script. Then we ran SVDquartets in PAUP* setting outgroup to *L. oculatus* and then executing evalQuartets=all with 100 standard bootstraps.

Next, we also used SNAPP (Bryant et al. 2012) as implemented in BEAST (Version 2.6.2)^36^ to constructed phylogenetic tree based on SNPs dataset. The “forward” and “backward” mutation rate parameters u and v were calculated directly from the data by SNAPP (the “Calc mutation” rates option). The default value 10 was used for the “Coalescent rate” parameter and the value of the parameter was sampled (estimated in the Markov chain Monte Carlo (MCMC) chain). The prior for ancestral population sizes was chosen to be a relatively broad gamma distribution with default parameters. We ran a 20,000,000-iteration chain with sampling every 50,000 iteration. Due to time limit, the program was terminated early, and finally 48,600,000-iteration were carried out.

**Phylogenetic discordance across the genome**

**Window-based gene trees**

To investigate the phylogenetic discordance across genomic regions, we segmented the WGA sequences in 10-kbp nonoverlapping windows. After excluding those windows in which included repeats sequences and sequences size less than 150 bp. A total of 9101 windows remains. We then constructed the window-based gene tree (WGT) of each window using IQtree with 100 standard bootstraps (command: iqtree -s <align. fa> -o <outgroup> -b 100 -m MFP). Subsequently, we applied ASTRAL^37^ to reconstruct the species tree from WGTs using the default parameters (for the detail, see “Visualizations of gene-tree discordance”).

**Visualizations of gene-tree discordance**

We filtered WGTs that included (*Danio rerio* (TU), *Danio rerio* (AB), *Danio rerio* (NA), *Danio rerio* (CB)) monophyletic clades. Then, we applied ASTRAL to reconstruct the species tree based on WGTs using the default parameters. Then, we used WGTs (7764 trees) to constructed DensiTree to visualize the phylogenetic conflicts. We first converted species tree to ultrametric trees using the R package Phybase (version 2.0)^38^, and then superimposed using DensiTree (version 2.2.5)^39^. Furthermore, we used DiscoVista (Discordance Visualization Tool, version 1.0)^40^ to analyze the gene-tree compatibility with the parameters: "-m 5", using the WGTs (Quartet frequencies of the internal branches in the species tree were calculated using ASTRAL).

We used all WGTs only included *D. rerio*, *D. aesculapii*, *D. kyathit*, *D. tinwini* and *D. nigrofasciatus* clades to analyze the topological differences among different positions of chromosomes. Firstly, according to the position from the telomere, the chromosomes were divided into five regions from both ends to the center, each region representing 20 % of the entire chromosome sequence. The chromosomes with less than 20 WGTs numbers were filtered out, and we finally got 14 chromosomes (Chr1 and Chr10-22) and 70 chromosome regions. Then, we used the WGTs in each region to do ASTRAL tree inference (the inference method was shown above), resulting in 70 ASTRAL region trees. Finally, we made topology type and frequency statistics on 70 ASTRAL region trees, and selected the five topologies with the highest frequency in the whole genome as the main topologies to do topological frequency statistics on different regions of chromosomes, in order to visualize the distribution preferences of evolutionary history in different locations of chromosomes. After the chromosomes which topology is (*D. rerio*, (*D. aesculapii*, *D. kyathit*)), (*D. tinwini*, *D. nigrofasciatus*)) in the central region were removed (Chr11, Chr17 and Chr19), repeated the above topological statistics work. Finally, 11 chromosomes and 55 ASTRAL region trees were retained, and the preference of chromosome topological distribution was analyzed. We also extracted from WGTs of 11 chromosomes only the topologies of *D. rerio*, *D. aesculapii* and *D. kyathit*, also analyzed the preference of chromosome topological distribution.

**Gene flow**

**D-Statistics**

We filtered SNP sites (for the detail of calling SNP, see “SNP and Indels analysis”) using BCFtools (version 1.8) with parameters “bcftools view -e 'AC==0 || AC==AN || F_MISSING > 0.2' -m2 -M2”. This resulted in a set of 93043 SNPs. We calculated the D-statistic for each trio of *D. rerio* subclade species (P1, P2, P3) without species tree (the outgroup again fixed as *D. albolineatus*), using Dsuite software^41^ (command: ./Build/Dsuite Dtrios <SNPs> <samples>). To get a better overview of introgression patterns supported by D-statistics, we used ggplot2 (version 3.3.2) to visualize these in the form of a bubble chart in which the species in positions P2 and P3 are sorted on the horizontal and vertical axes, and the circle of bubble chart indicates the most significant D-statistic found with these two species, across all possible species in P1. Script modified from <https://github.com/mmatschiner/tutorials/tree/master/analysis_of_introgression_with_snp_data>.

**D_FOIL_ statistics**

Pease and Hahn proposed a five-taxon test to distinguish ILS from gene flow (the D_FOIL_ statistics). D_FOIL_ analyses assume a symmetrical five-taxon topology: (((P1, P2), (P3, P4)), O), which can determine the direction of any detected introgression phylogeny. We used *L. oculatus* as outgroup, and (*D. tinwini*, *D. nigrofasciatus*), (*D. erythromicron*, *D. margaritatus*) or (D. *choprae*, D. jaintianensis) as (P3, P4) with other Danio species as (P1, P2) to calculate the D_FOIL_ statistics for all the possible four fitted topologies based on WGA sequences (excluded repeat sites) using D_FOIL_^42^. We also used *D. albolineatus* as outgroup to detect geneflow between species for *D. rerio* subgroup. We tested five fitted topologies: (1) (((*D. tinwini*, *D. nigrofasciatus*), (*D. rerio*, *D. aesculapii*)), *D. albolineatus*); (2) (((*D. tinwini*, *D. nigrofasciatus*), (*D. rerio*, *D. kyathit*)), *D. albolineatus*); (3) (((*D. tinwini*, *D. nigrofasciatus*), (*D. aesculapii*, *D. kyathit*)), *D. albolineatus*); (4) (((*D. rerio*, *D. aesculapii*), (*D. kyathit*, *D. tinwini*)), *D. albolineatus*); (5) (((*D. rerio*, *D. aesculapii*), (*D. kyathit*, *D. nigrofasciatus*)), *D. albolineatus*). The 10175 10-kbp windows (excluded repeat sites and sequences size no less than 150 bp), were used to analyze gene flow by using D_FOIL._

**Demographic history reconstruction**

**Divergence time calibration**

Divergence time estimation was performed using the MCMCTREE in PAML4.7 package^43^. The upper and lower limit of the divergence time found on the TIMETREE website (<http://www.timetree.org/>) was used as the calibration time: *D. rerio* - *D. erythromicron* (36-68 Mya); *D. rerio* - *Danionella cerebrum* (36-68 Mya); *A. mexicanus* - *I. punctatus* (109-157 Mya); *D. rerio* - *O. latipes* (206-252 Mya); *D. rerio* - *L. oculatus* (295-334 Mya). Then, we used MCMCTREE in PAML4.7 package^43^ to estimate the divergence time based on three different topological evolution trees (1&2 codon sites BI tree, 4d sites BI tree and SNP tree constructed by SNAPP) with calibration time and the corresponding multiple sequence alignment files. We first calculated substitution rate using baseml in PAML, then set usedata=3 and clock=2 to generate out.BV using MCMCTREE, and finally set usedata=2 and clock=2 to calculate the divergence time using MCMCTREE.

**Demographic history reconstruction**

We inferred the demographic history for Danio species by applying the pairwise sequentially Markovian coalescence model (PSMC)^44^. We used aligned reads and consensus sequences to conduct the PSMC (version 0.6.5-r67) analysis. Firstly, we applied bwa (version 0.7.15-r1140) with the default parameters to align sequencing reads to the assembled species genome. Next, we used SAMtools (version 1.10)^21^ to convert the aligned results to bam files (with the parameters: samtools view -S -b) and merged them into one file (with the parameters: samtools merge). Then, we sorted the bam file (with the parameters: samtools sort) and remove PCR duplicate reads (with the parameters: samtools rmdup). In order to obtain a diploid consensus genome sequence, we estimated genotype likelihoods with an adjusted mapping quality greater than 50, then used BCFtools (version 1.3.1) to identify SNPs, and finally utilized "vcfutils.pl" to exclude sites with mapping depths > 1000 or < 1. Specifically, we used the commands:

samtools mpileup -C50 -uf <genome. fa> <species.MERGED.SORTED.rmdup.bam> | bcftools call -c - | vcfutils.pl vcf2fq -d 1 -D 1000 | gzip > <species.fq.gz>

Finally, we used the file to conduct the PSMC analysis. We transformed the format of consensus sequence by using fq2psmcfa (with the parameters: fq2psmcfa -q20). After the transformation, the population size histories were inferred by PSMC (with the parameters: psmc -N25 -t15 -r5 -p “4+25*2+4+6”), and these results were then scaled to absolute time and population sizes using generation times and estimated per generation mutation rates by running “psmc2history.pl” and “history2ms.pl” with the default parameters. To visualize the result with “psmc_plot.pl” with the parameters: -g <estimates generation time> -u <neutral mutation rates>. The generation time (g) of *Danio* species were obtained from previous studies. The per year mutation rates were estimated by PAML. The per generation mutation rate is estimated by multiplying the per year mutation rates with the generation time.

**Biogeographic Analysis**

We retrieved hydrological basin data from Aquastat, the Food and Agricultural Organization of the United Nations’s global water information system (<http://www.fao.org/nr/water/aquamaps/>). We downloaded information of the locality of *Danio* species from the Global Biodiversity Information Facility (gbif.org) using the R package “rgbif”^45^ and imported them into ArcGIS (version 10.3) for visualization and comparison.

**SM Text 4. Genomic features related to danio evolution**

**Identification of positively selected genes (PSGs)**

We used a conserved genome synteny methodology to establish a high-confidence orthologous gene set that included *Lctalurus punctatus, Astyanax mexicanus, Oryzias latipes, Lepisosteus oculatus*, and these *Danio* species. Briefly, pairwise WGAs were constructed for relevant genomes using LAST^24^, with the GRCz11 serving as the reference genome. To minimize the effect of annotation, sequencing and assembly errors, pseudogenes, non-orthologous alignments, and non-conserved gene structures on subsequent evolutionary rate analyses, a series of rigorous filtering criteria were adopted: (1) the genes mapped to the reference genome via a single chain of sequence alignments including at least 80 % of its coding sequence (CDS), and met the alignment length/score thresholds required for inclusion in the MULTIZ alignments (Blanchette et al. 2004); (2) frame-shift indels in CDSs were prohibited; (3) CDSs with premature stop codons were excluded and (4) genes with Ks values (synonymous substitutions per synonymous site) between each species and cattle larger than two were excluded.

Based on the filtered orthologous gene set, we estimated the lineage-specific evolutionary rate for each branch. The Codeml program in the PAML package (version 4.8) with the free-ratio model (model=1) was run for each ortholog. Positive selection signals on genes along specific lineages were detected using the optimized branch-site model following the author's recommendation^46^. A likelihood ratio test (LRT) was conducted to compare a model that allowed sites to be under positive selection on the foreground branch with the null model in which sites could evolve either neutrally and under purifying selection. The *p*-values were computed based on Chi-square statistics, and genes with *p*-value less than 0.05 were treated as candidates that underwent positive selection. KEGG and GO enrichment analyses were applied in KOBAS (Wu et al. 2006) for the expanded and contracted gene families.

**Gene family expansion and contraction in danios**

***Gene family contraction***

Reference protein sequences of *Lepisosteus oculatus, Oryzias latipes, Ictalurus punctatus, Astyanax mexicanus* and *Danio rerio* were downloaded from Ensembl release 92. Then, the protein sequences of other danio species were extracted from our annotated genomes. Protein sequences, fewer than 30 amino acids or containing premature codons, were removed. This filtering resulted protein sequences were passed to OrthoFinder^47^ for protein clustering.

**Identification of expanded/contracted gene families with CAFÉ**

The gene family expansion or contraction analysis was performed using CAFÉ (parameter with -p 0.05 -r 100 –filter)^48^. In CAFÉ, a random birth-and-death model is used to study expansion and contraction in gene families across a user-specified divergence time tree, for which we obtained by r8s. KEGG and GO enrichment analyses were applied in KOBAS for the expanded and contracted gene families**.**

Several of expanded gene families are classified into important functions related with pigment pattern.

**SM Text 5. Genomic variations related to *Danio* characteristics**

To explore the difference of pigment trait of *Danio* species, we selected 38 pigment-related genes and investigated the distribution of these genes in 5 chromosome-level or super scaffold-level genomes of *Danio* species using MCScanX python (<https://github.com/tanghaibao/jcvi/wiki/MCscan-(Python-version))>.

**Commands/parameters used:**

**Data prepare:**

python -m jcvi.formats.gff bed --type=mRNA --key=Name speciesA.gff3 -o speciesA.bed

python -m jcvi.formats.fasta format speciesA.cds.fa speciesA.cds

python -m jcvi.formats.gff bed --type=mRNA --key=Name speciesB.gff3 -o speciesB.bed

python -m jcvi.formats.fasta format speciesA.cds.fa speciesB.cds

python -m jcvi.formats.gff bed --type=mRNA --key=Name speciesC.gff3 -o speciesC.bed

python -m jcvi.formats.fasta format speciesA.cds.fa speciesC.cds

**Pairwise synteny search:**

python -m jcvi.compara.catalog ortholog speciesA speciesB

python -m jcvi.compara.catalog ortholog speciesB speciesC

python -m jcvi.compara.synteny screen --minspan=30 --simple speciesA.speciesB.anchors speciesA.speciesB.anchors.new

python -m jcvi.compara.synteny screen --minspan=30 --simple speciesB.speciesC.anchors speciesB.speciesC.anchors.new

**Macrosynteny visualization:**

python -m jcvi.graphics.karyotype seqids layout

Then, in chromosome/ super scaffold 9, 22 and 23, each trait-related gene and 10 upstream/downstream neighbor genes were chosen to further research the difference of traits related genes in different species.

**Commands/parameters used:**

**Microsynteny visualization:**

python -m jcvi.compara.synteny mcscan speciesA.bed speciesA.speciesB.lifted.anchors --iter=1 -o speciesA.speciesB.i1.blocks

python -m jcvi.compara.synteny mcscan speciesA.bed speciesA.speciesC.lifted.anchors --iter=1 -o speciesA.speciesC.i1.blocks

python -m jcvi.formats.base join speciesA.speciesB.i1.blocks speciesA.speciesC.i1.blocks --noheader | cut -f1,2,4,6 > Danio.blocks

cat speciesA.bed speciesB.bed speciesC.bed > speciesA_speciesB_speciesC.bed

python -m jcvi.graphics.synteny blocks2 speciesA_speciesB_speciesC.bed blocks2.layout

Depend on these results, some traits related genes display the difference among these 5 *Danio* genomes. And these differences were classified into 3 types: present-absence type (PAT), insertion type (IT) and different length type (DLT). For present-absence type, after we rechecked the results by BLAST, 6 PAT genes were identified in chromosome/ super scaffold 9 and chromosome/ super scaffold 23, including 2 pigment related genes. And most of insertion type genes because of chromosome rearrangement events.

**SM Figures**

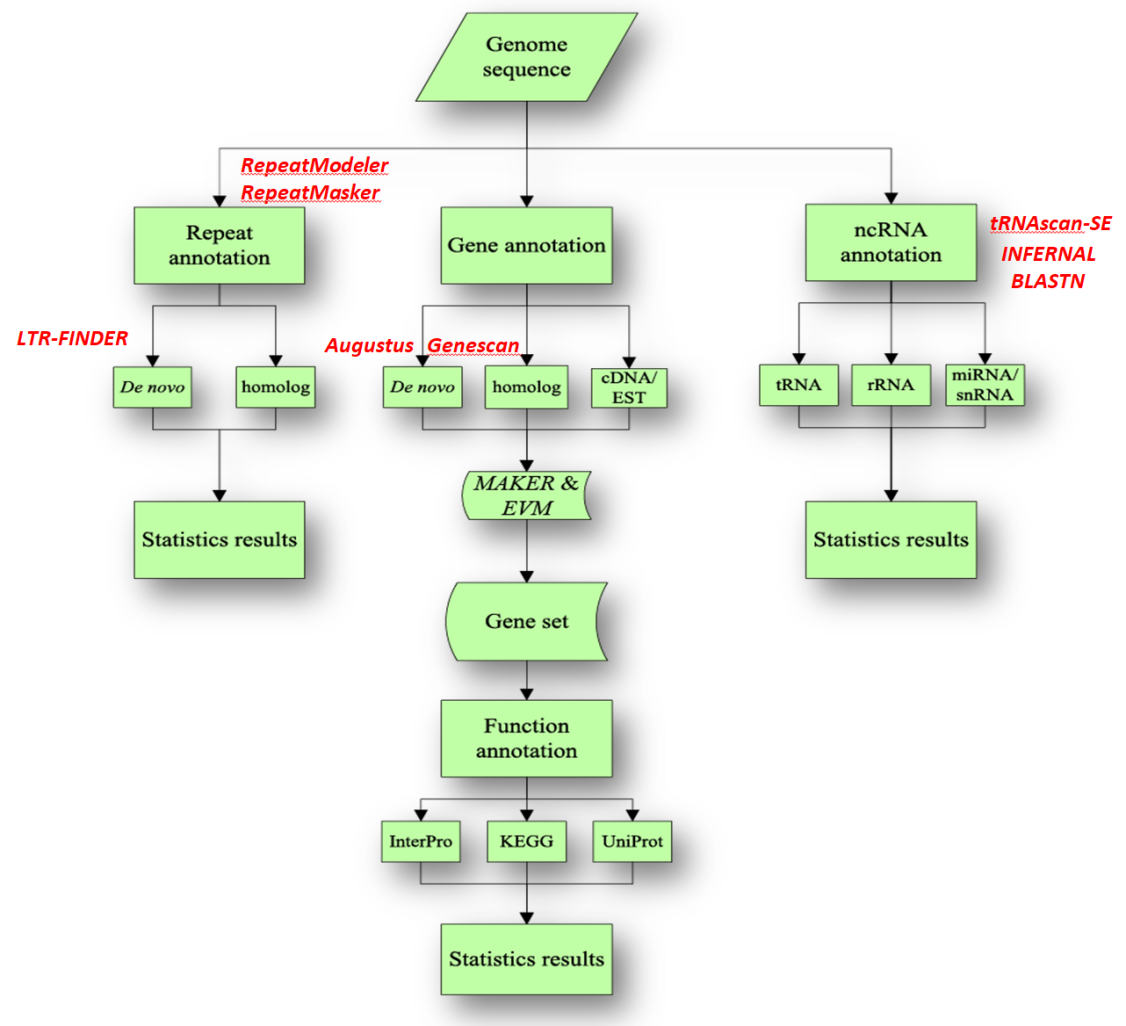

**Figure S1 | Pipeline of genome annotation analysis.** Repeat elements, genes and ncRNAs of danios were annotated with different software.

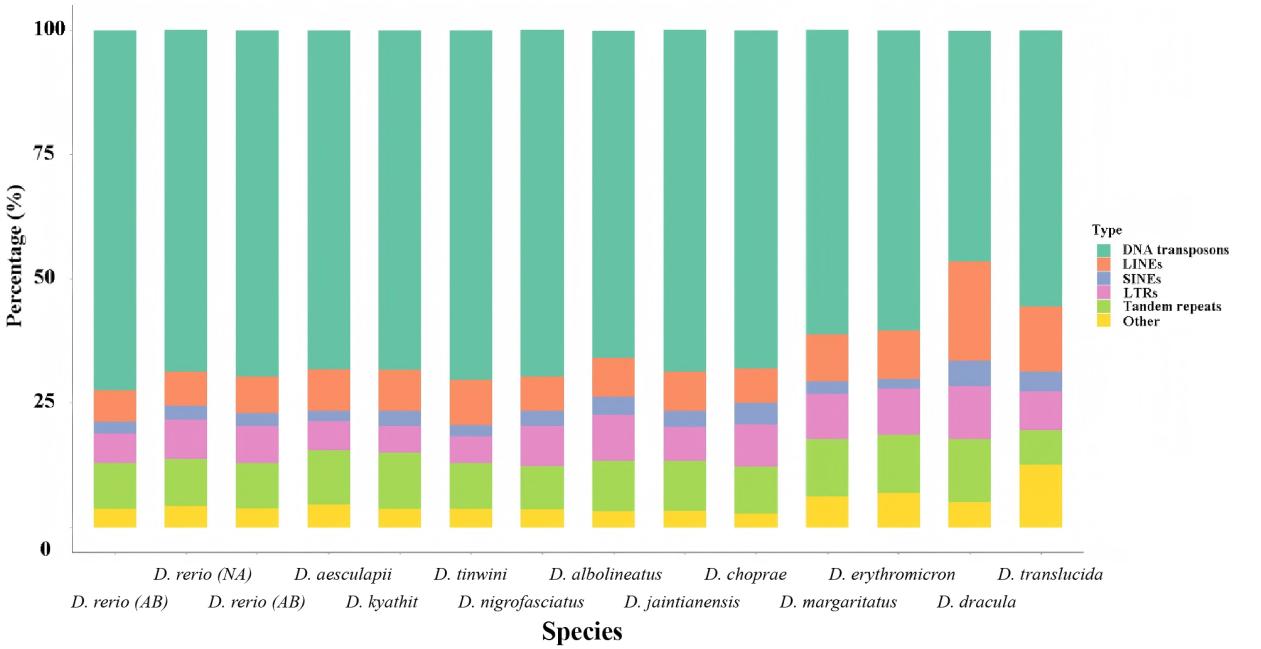

**Figure S2 | Summary of repeat sequences from 15 Danio genomes.** Different categories of repeat sequences are shown in this figure, including DNA transposons, LINEs, SINEs, LTRs, tandem repeats and others.

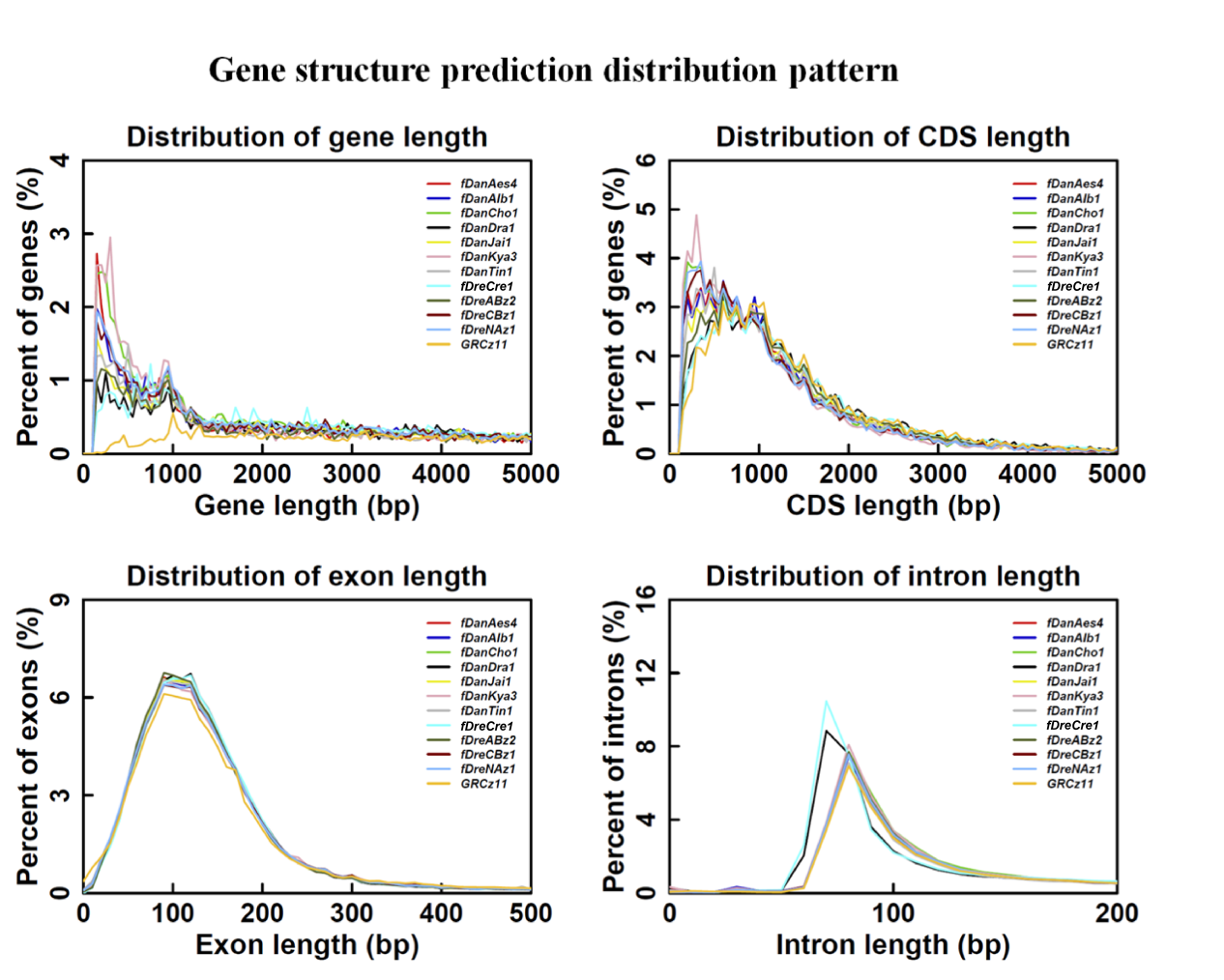

**Figure S3 | Gene structure prediction distribution pattern.** *fDanAes4: D. aesculapii; fDanAlb1: D. albolineatus; fDanCho1: D. choprae; fDanDra1: Danionella dracula; fDanJai1: Danio jaintianensis; fDanKya3: D. kyathit; fDanTin1: D. tinwini; fDanTra1: Danionella cerebrum; fDreABz2: D. rerio (AB); fDreCBz1: D. rerio (CB); fDreNAz1: D. rerio (NA); GRCz11: D. rerio*.

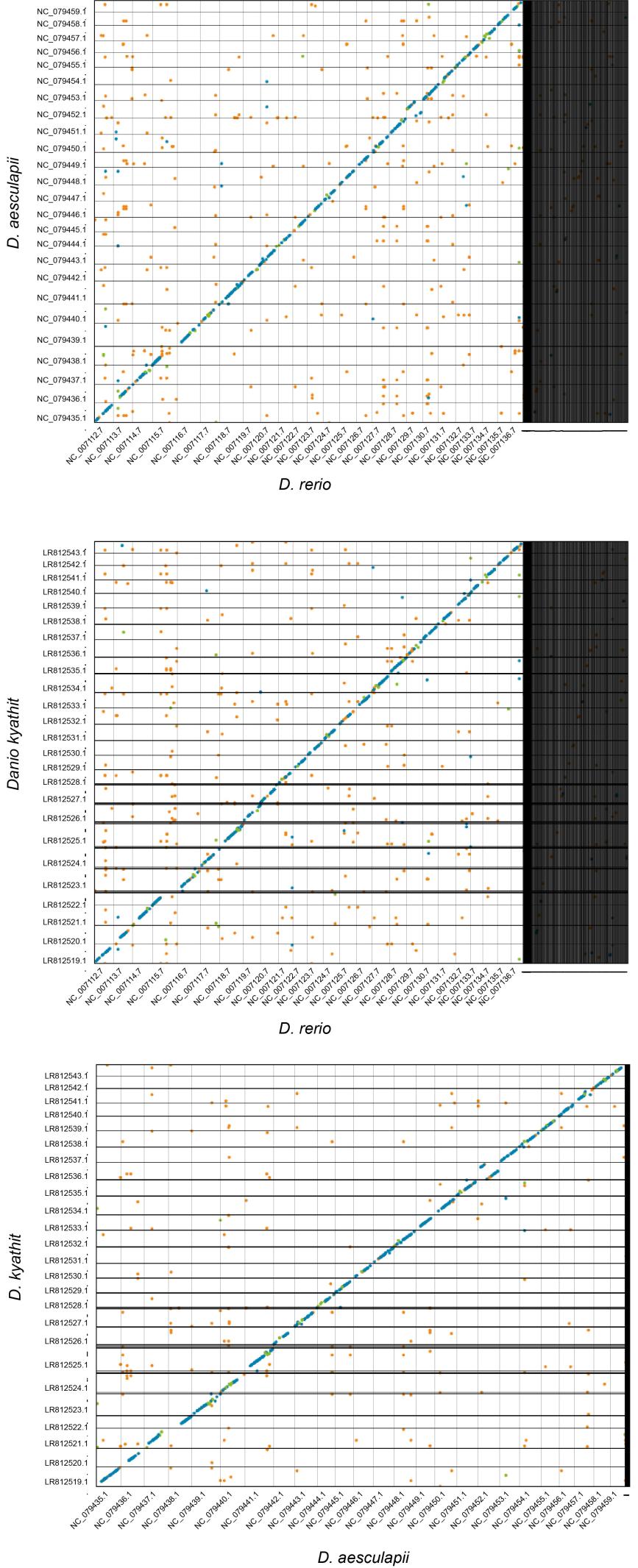

**Figure S4 | Whole genome synteny relationship among *Danio aesculapii (Aes), Danio kyathit (Kya) and Danio rerio (Rer).*** In the pairwise synteny plots, the X- and Y-axes represent chromosomes of the respective species. Blue dots represent unique forward alignments; green dots represent unique reverse alignments; orange dots represent repetitive alignments. Both chromosomes and scaffolds are included in the collinearity analysis. Black grid lines separate the chromosomes or scaffolds. Some of the thicker-appearing black lines in the figure may be due to the shorter length of the scaffolds and the smaller intervals between them.

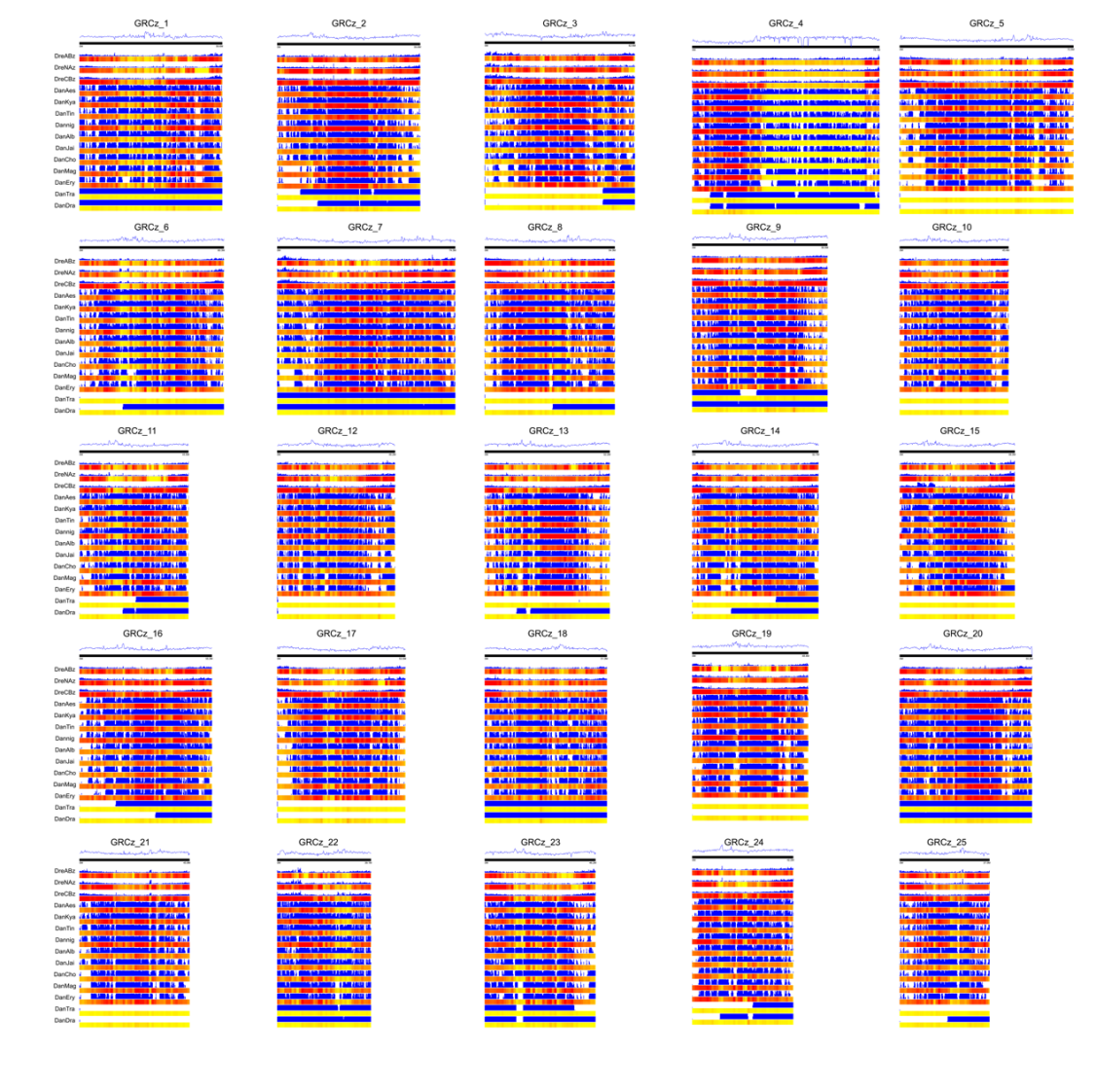

**Figure S5 | The liner maps of variants called from 14 danios relative to the reference.** The curve graph indicates the repeats content of each chromosome. Heatmap displays the density of SNP of each chromosome. PAVs are shown in the figure as blue blocks.

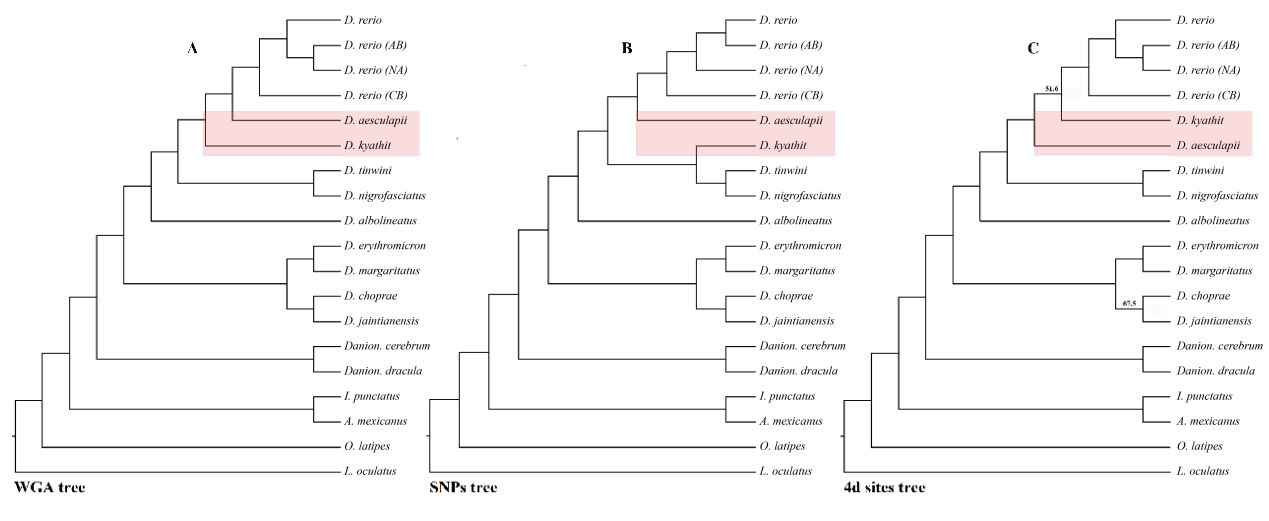

**Figure S6 | Incongruences among WGA, SNPs and 4d sites trees of the Danios.** A. WGA tree. The maximum likelihood phylogenetic tree from whole-genome sequences of 13 Danio species and 6 outgroup species. To compute the node supports, 100 standard bootstraps were used, and all nodes have 100 % support. B. SNPs tree. SNPs tree was constructed based on multispecies coalescent method by SVDquartets as implemented in PAUP*. To compute the node supports, 100 standard bootstraps were used, and all nodes have 100 % support. C. 4d sites tree. 4d sites of orthologous genes were extracted and used for inferring phylogenetic trees. Only two nodes had <100 % support. Labeled nodes give the support for the node in the ML tree. ML tree was constructed by PhyML based on HKY85 model. Bayesian (BI) tree was constructed by Mrbayes based on GTR+Gamma model. Topological conflicts between *D. aesculapii* and *D. kyathit* are highlighted in red.

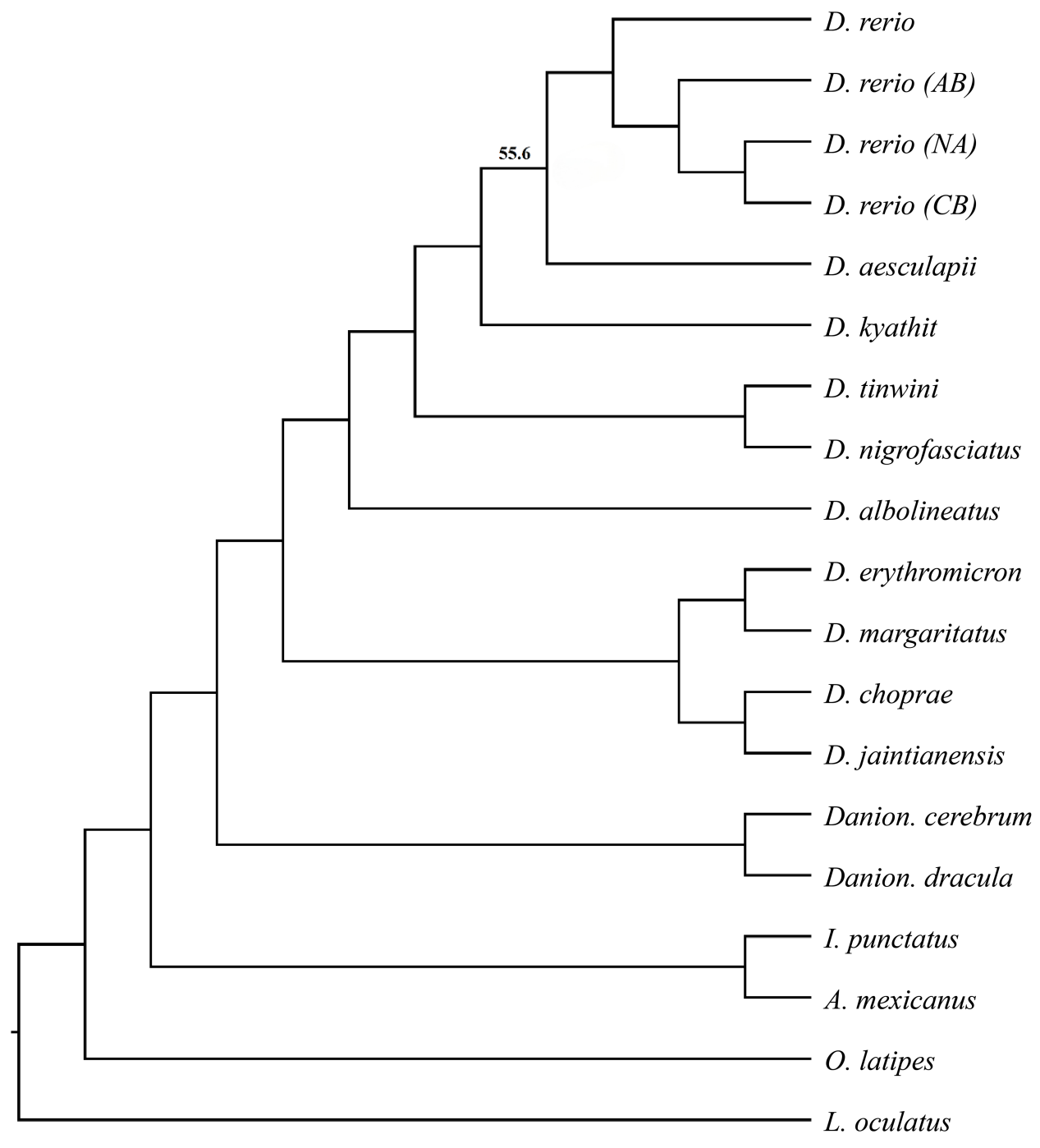

**Figure S7 | ML and BI phylogenetic tree inferred using 1st codon nucleotide of orthologous genes.** Only one node had < 100 % support. Labeled nodes give the support for the node in the ML tree. ML tree was constructed by PhyML based on HKY85 model. Bayesian (BI) tree was constructed by Mrbayes based on GTR+Gamma model.

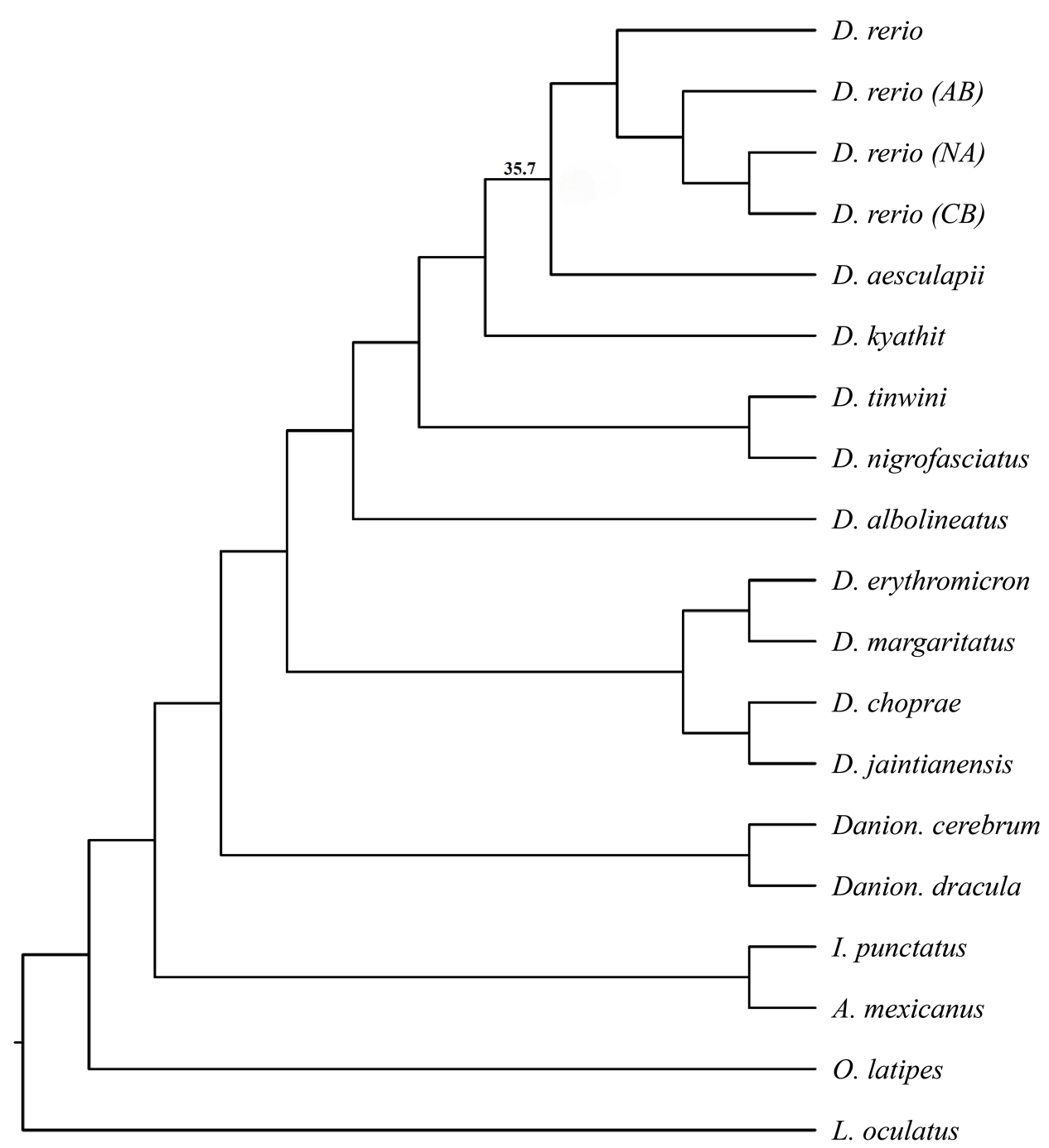

**Figure S8 | ML and BI phylogenetic tree inferred using 1st - 2nd codon nucleotide of orthologous genes.** Only one node had < 100 % support. Labeled nodes give the support for the node in the ML tree. ML tree was constructed by PhyML based on HKY85 model. Bayesian (BI) tree was constructed by Mrbayes based on GTR+Gamma model.

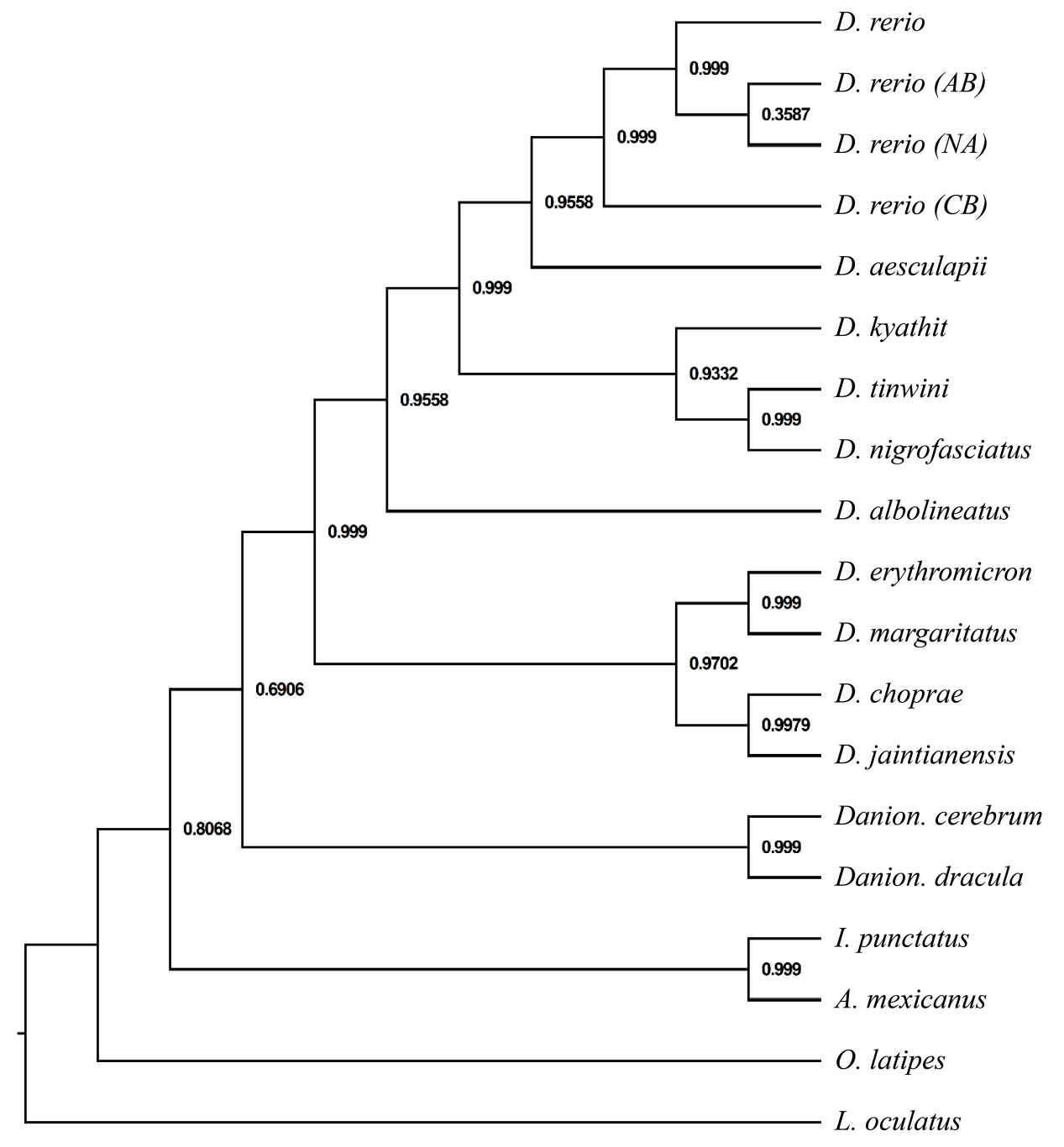

**Figure S9 | SNAPP SNPs tree.** SNPs tree was constructed based on multispecies coalescent method by SNAPP as implemented in BEAST. The local posterior probability is labeled on each node.

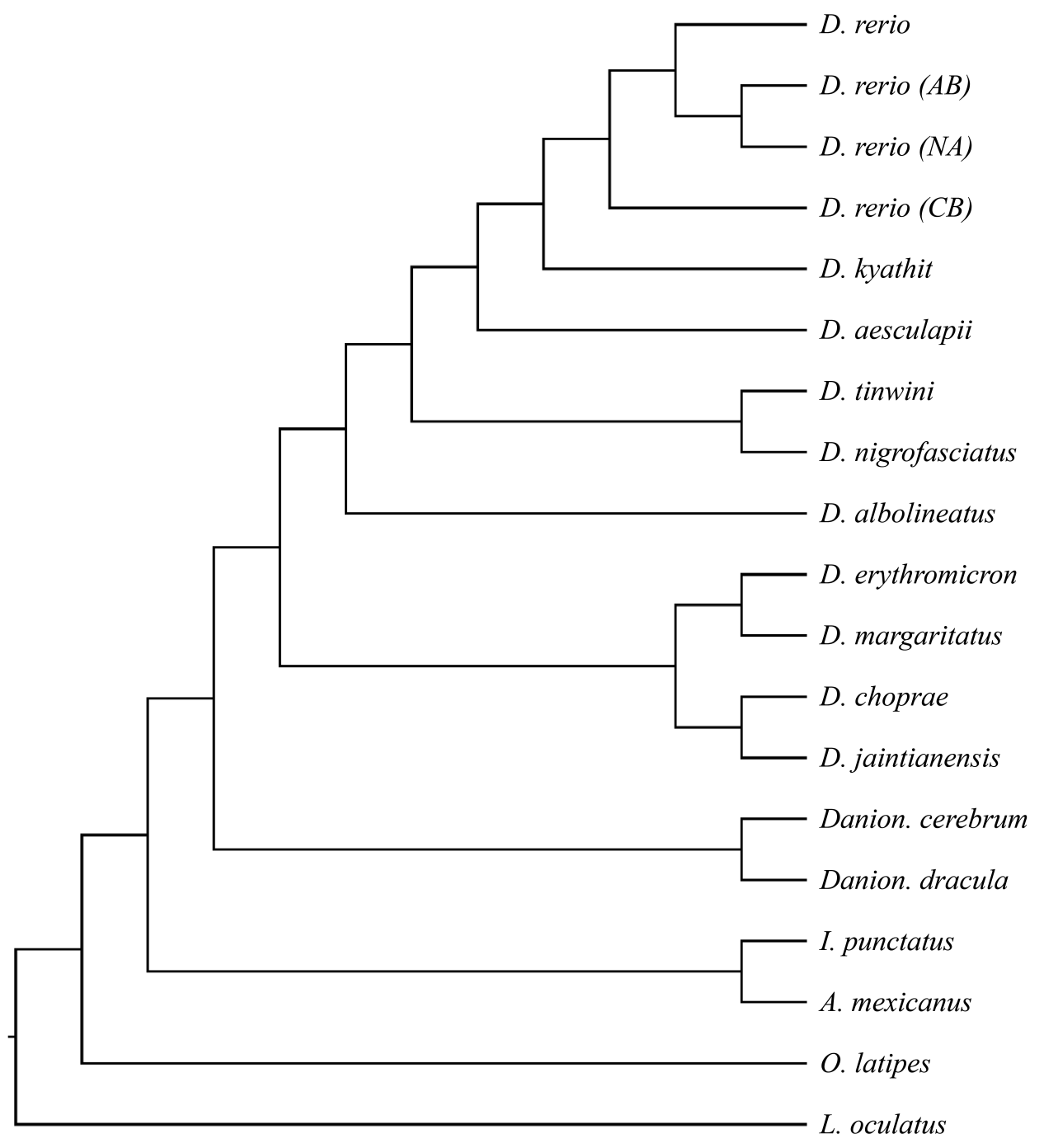

**Figure S10 | ASTRAL tree inferred from window-based gene trees by ASTRAL.** All nodes have 100 % support.

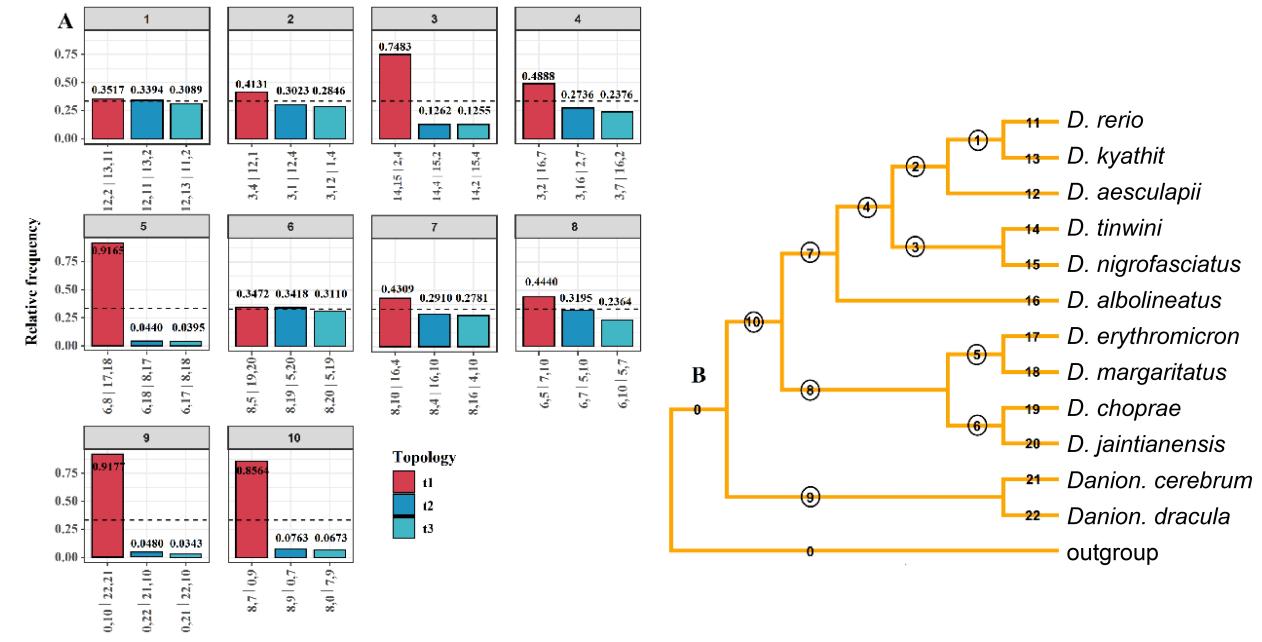

**Figure S11 | Quartet frequencies of branches based on WGTs.** Each internal branch with four neighboring branches would present three possible topologies. The frequency of the three topologies around focal internal branches of ASTRAL trees were computed using DiscoVista from the WGTs. A. The frequency of the main topology (found in the ASTRAL tree) is t1, and the other two alternative topologies are shown with blue bars. The dotted line indicates the 1/3 threshold expected at random. The number of each subfigure indicates the circled number label of the corresponding branch on the tree in B. In the x-axis, the exact definition of each quartet topology is provided using the neighboring branch labels separated by “|”. B. Each internal branch has four neighboring branches which could be used to represent quartet topologies. Branches are collapsed at the species level and marked with numbers.

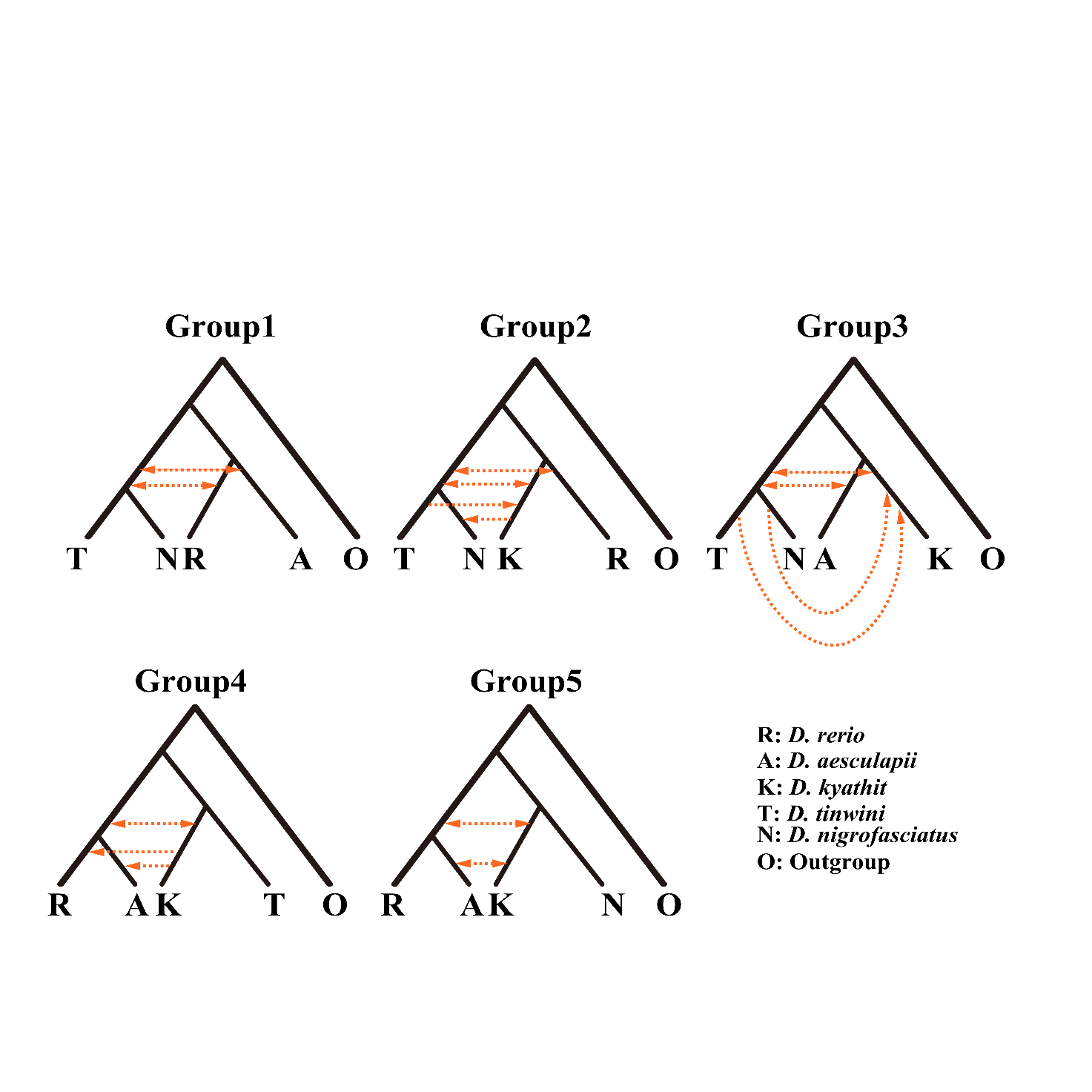

**Figure S12 | Gene flow was tested with the five-taxon topology approach using D_FOIL_.** These showed the gene flow signals between different species of *D. rerio* subclade. The orange arrow indicates the direction of gene flow.

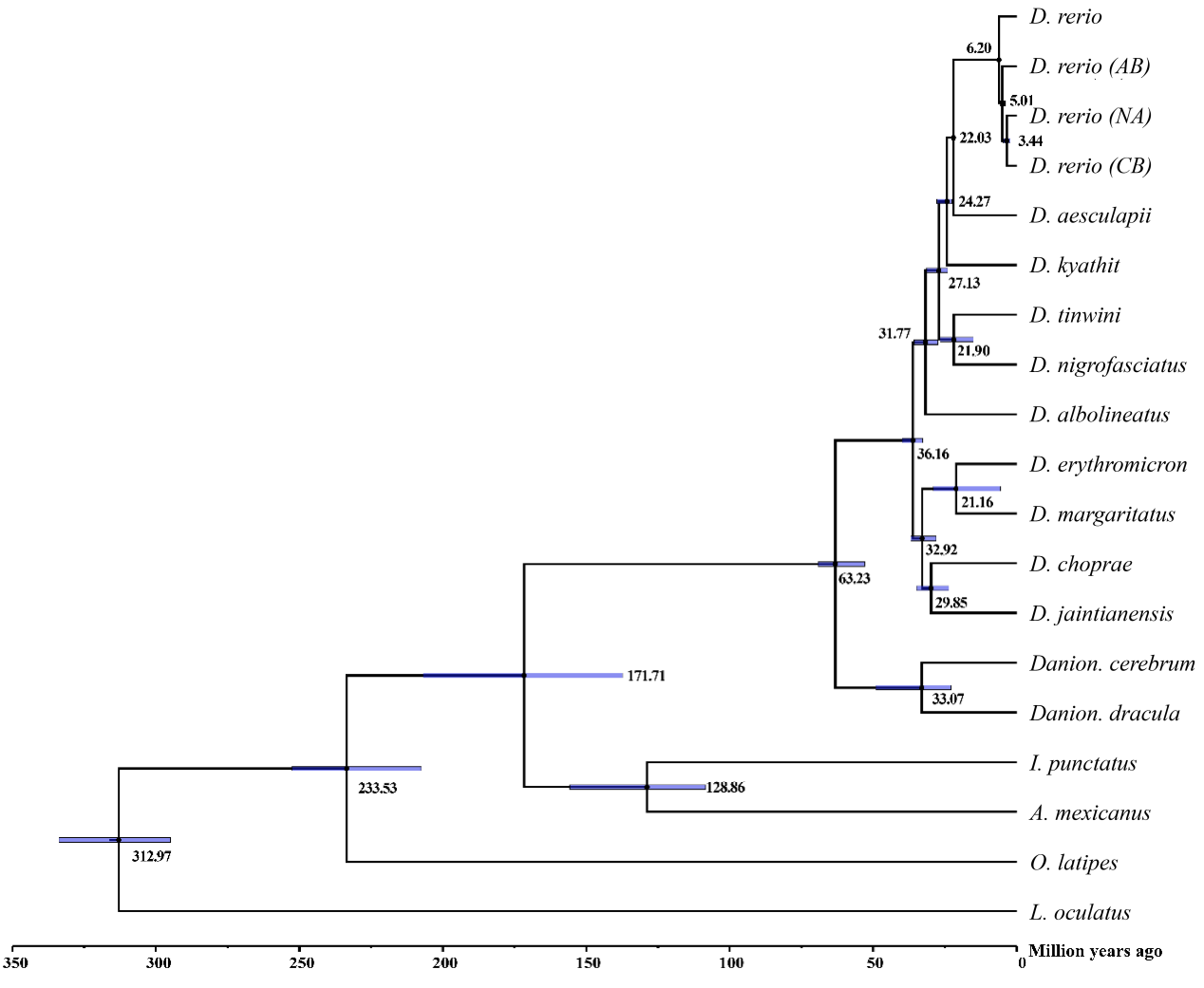

**Figure S13 | Divergence times was inferred by MCMCTREE based on 1&2 codon sites.** The estimated divergence time (the unit is million years) is labeled on each node, and the blue bars represent the 95 % confidence intervals.

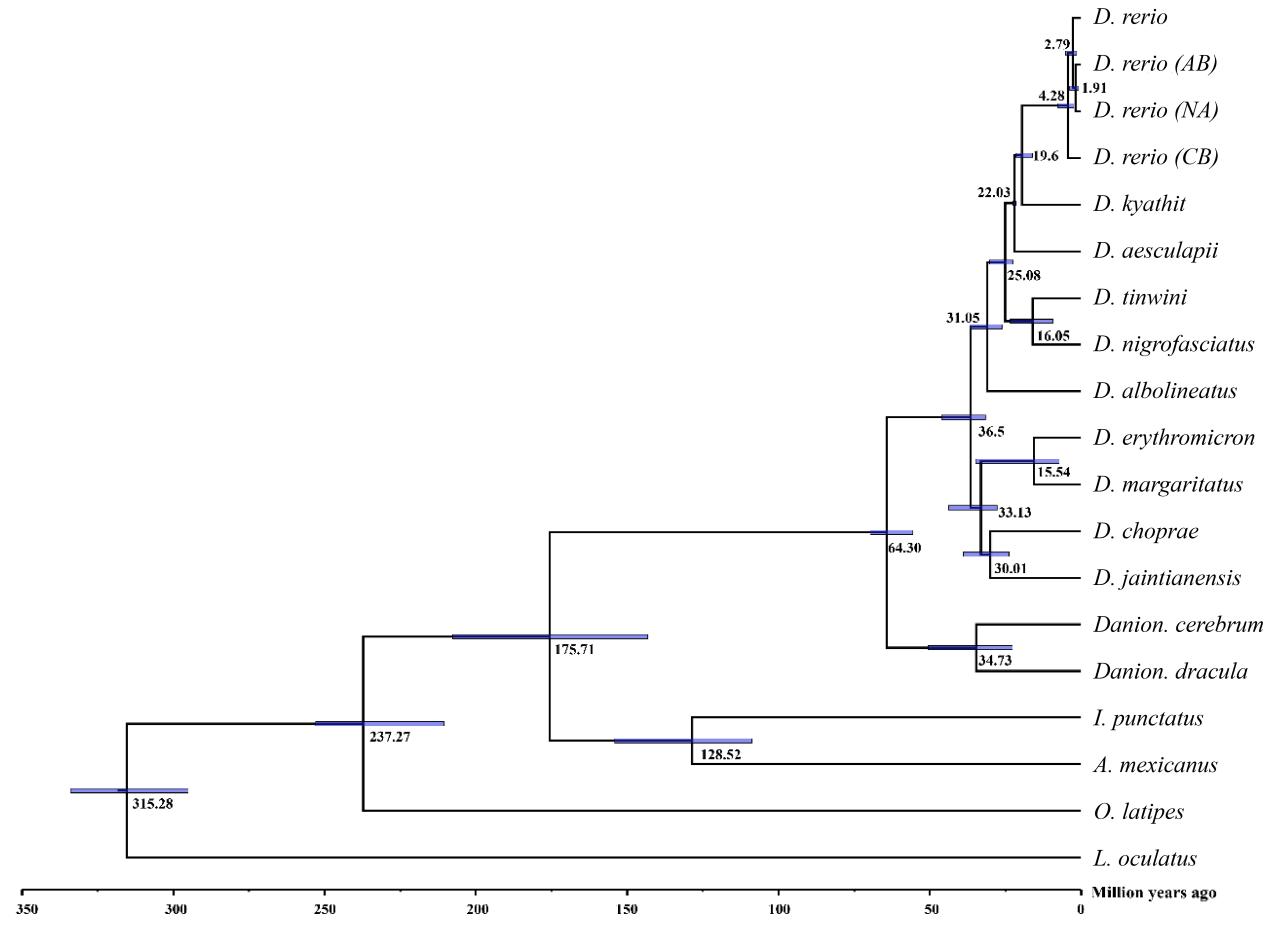

**Figure S14 | Divergence times was inferred by MCMCTREE based on 4d sites.** The estimated divergence time (the unit is million years) is labeled on each node, and the blue bars represent the 95 % confidence intervals.

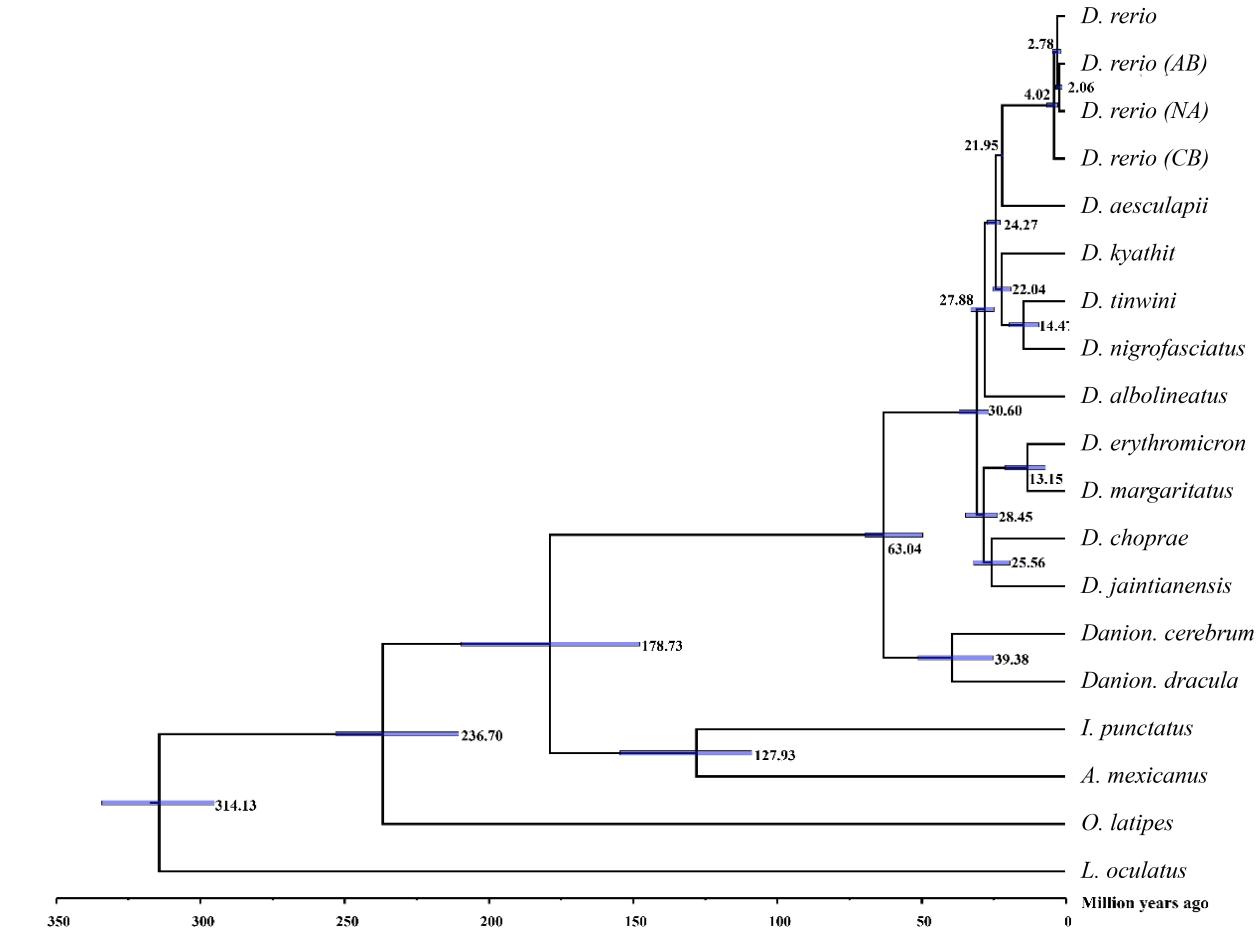

**Figure S15 | Divergence times was inferred by MCMCTREE based on SNAPP SNPs tree.** The estimated divergence time (the unit is million years) is labeled on each node, and the blue bars represent the 95 % confidence intervals.

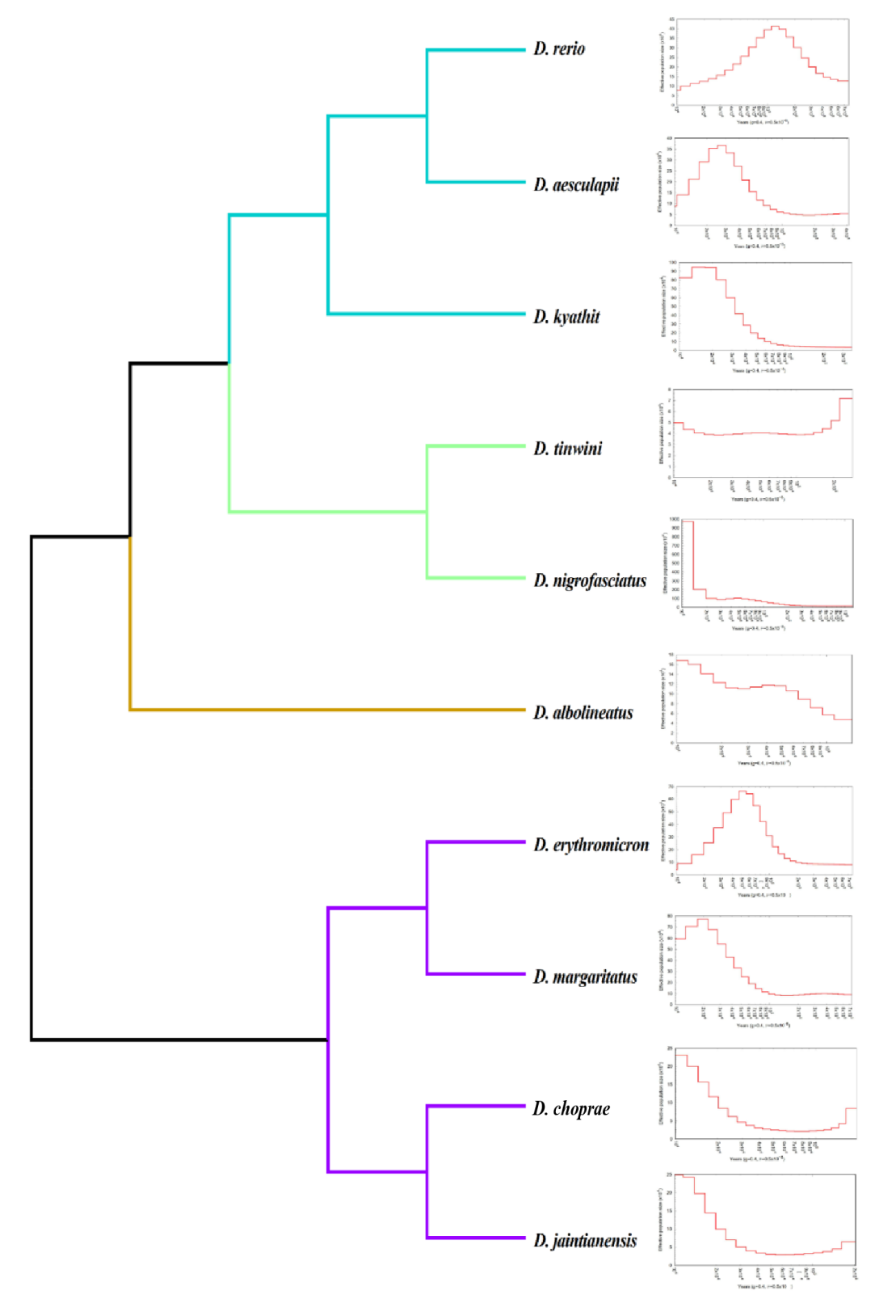

**Figure S16 | Population size history of Danios.** The dynamic changes of the effective population size are performed using PSMC software. The parameter “g” represents the generation length, and the parameter “μ” means the per generation mutation rate.

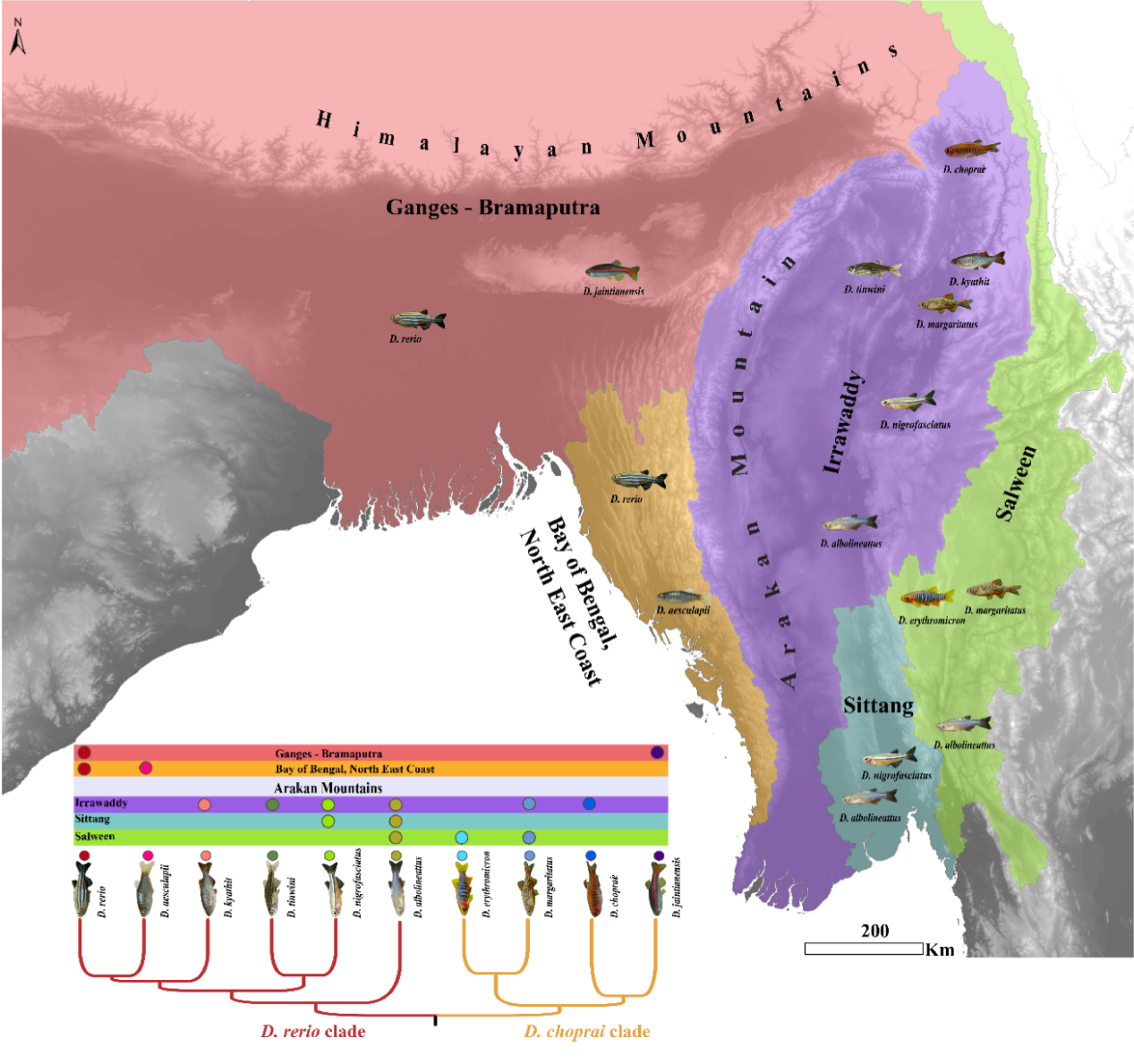

**Figure S17 | Phylogeography of Danios.** The Arakan Mountains of Myanmar separate *D. rerio* and *D. aesculapii* from all other Danios in this study.

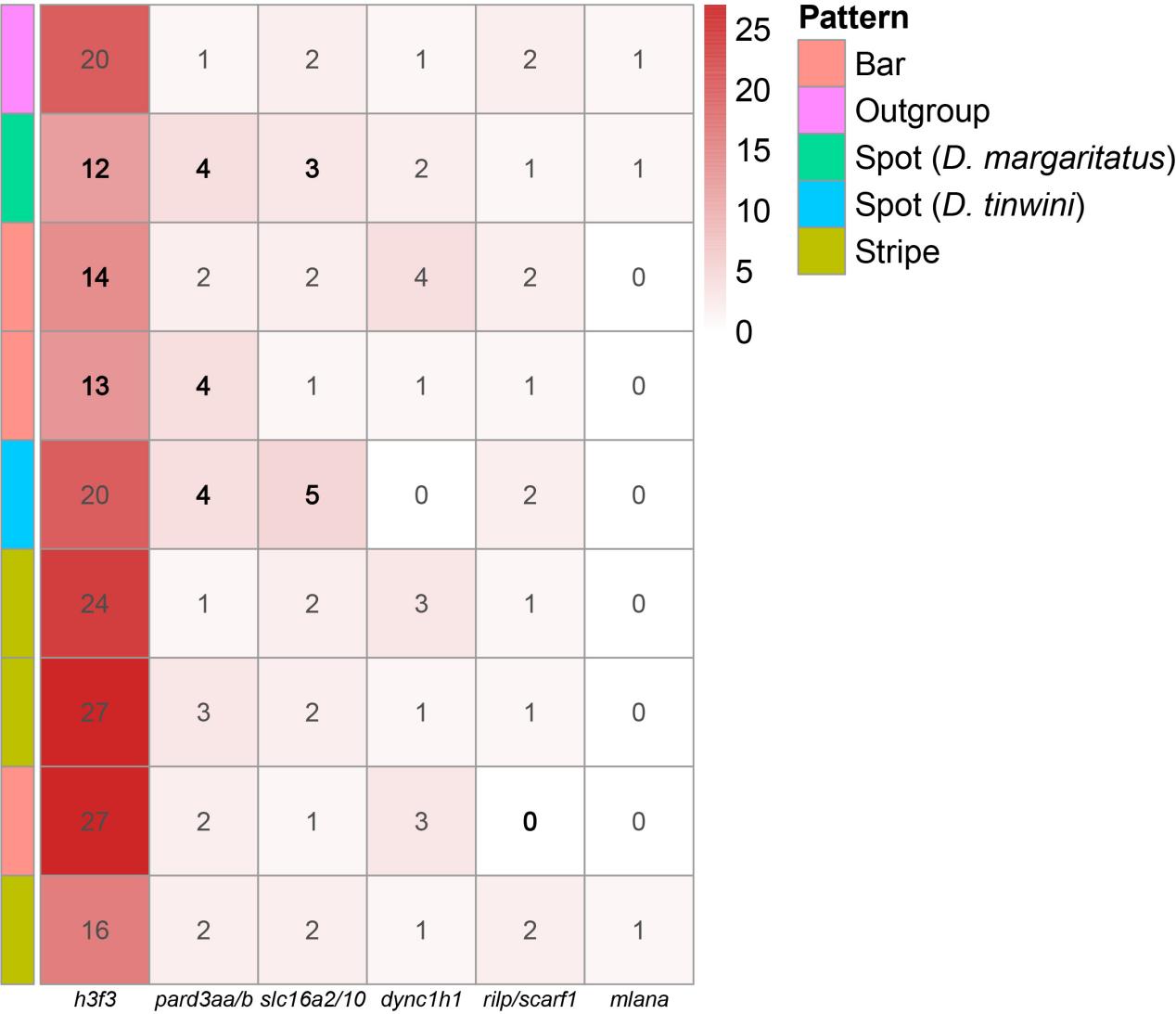

**Figure S18 | Heatmap of member number in the expanded / contracted pigment-related orthogroups.** Species were classified into five groups according to their pigment pattern: striped (*D. nigrofasciatus*, *D. kyathit*, *D. rerio*), spotted (*D. tinwini*), spotted (*D. margaritatus*), barred (*D. erythromicron, D. choprae, D. aesculapii*) and outgroup (*Ictalurus punctatus*).

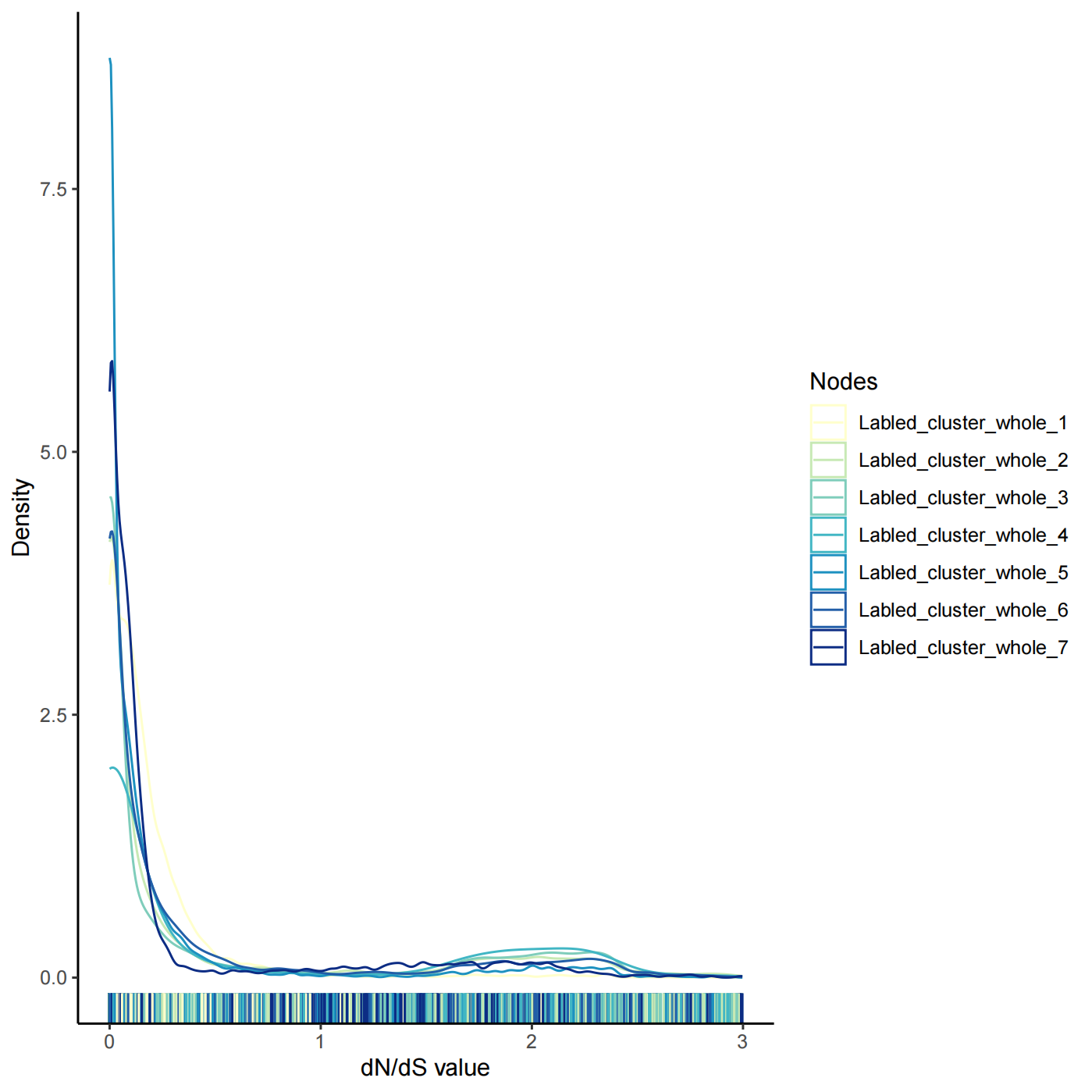

**Figure S19 | Density curves of the dN/dS value of all single-copy genes at each node within Danio.** Node order was the same as Fig. 4.

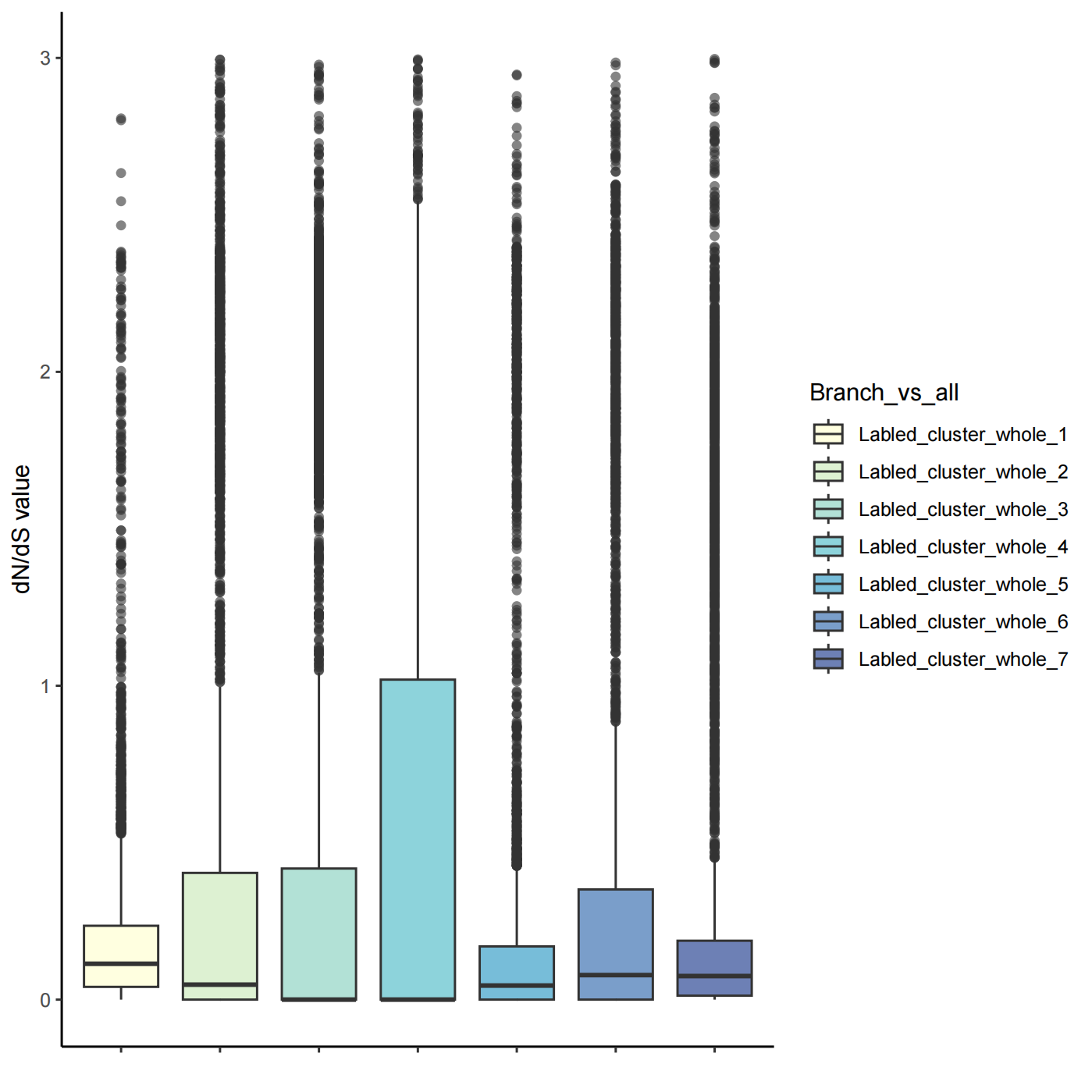

**Figure S20 | Box plot of the dN/dS value of all single-copy genes at each node within Danio.** Node order was the same as Fig. 4.

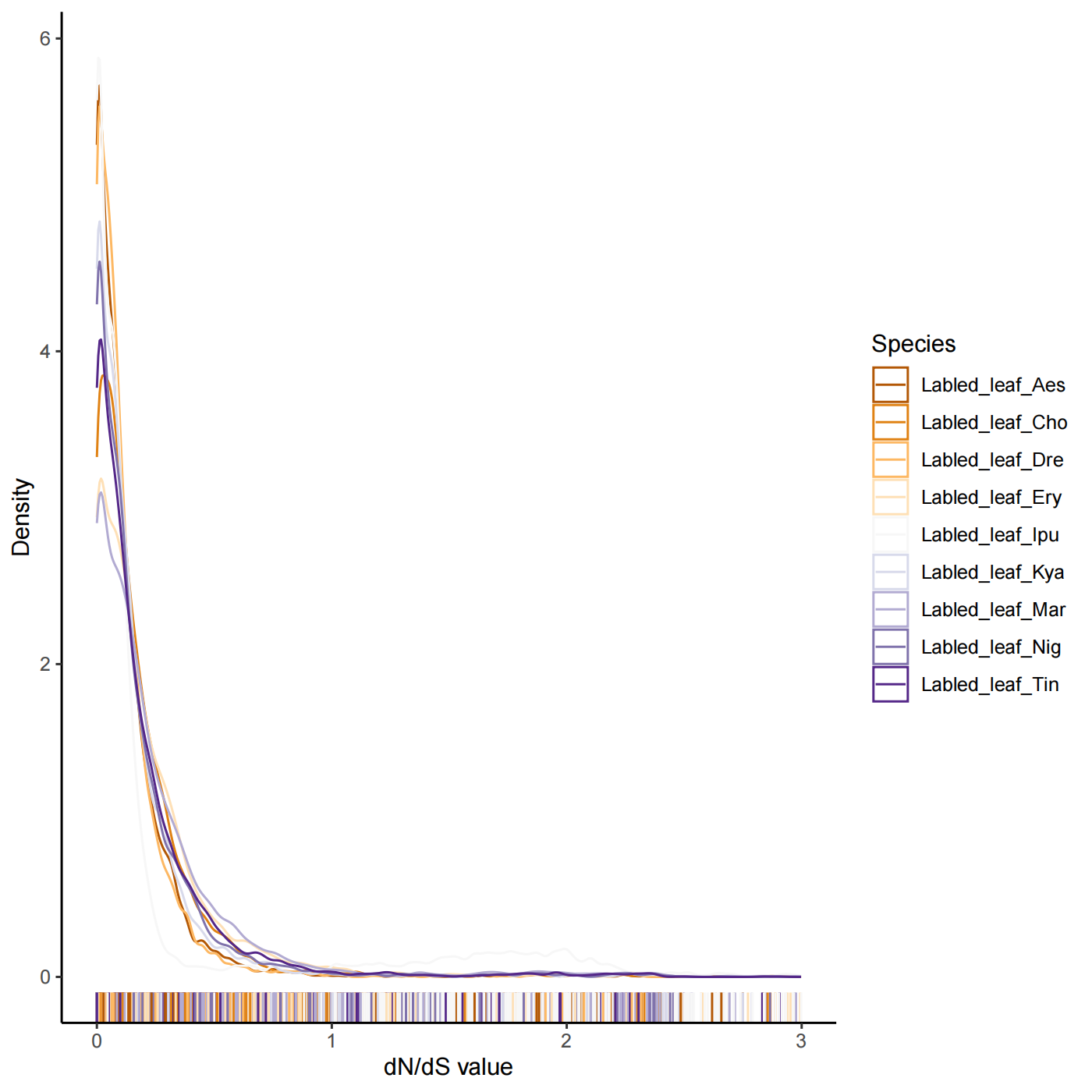

**Figure S21 | Density curves of the dN/dS value of all single-copy genes at each leaf.**

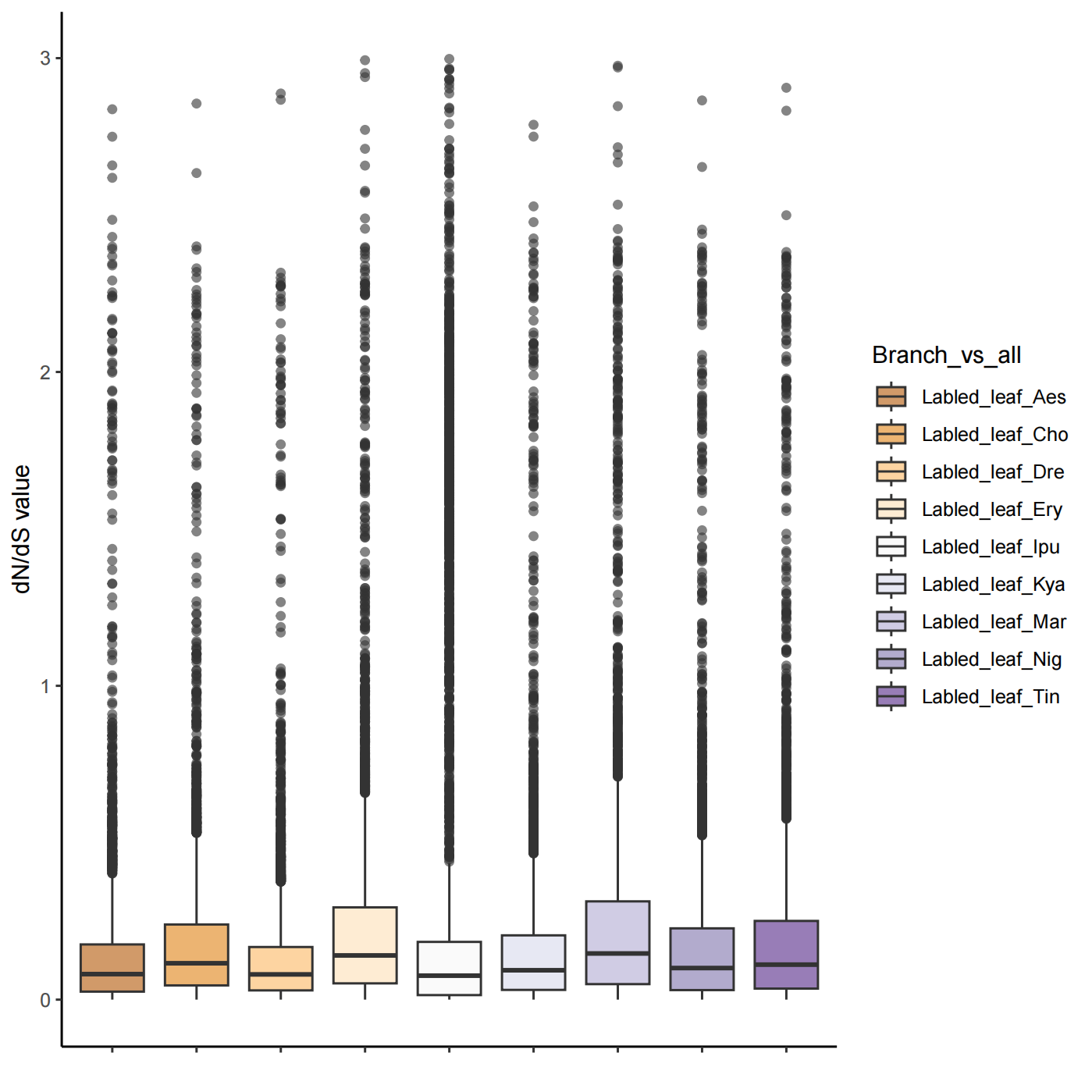

**Figure S22 | Box plot of the dN/dS value of all single-copy genes at each leaf.**

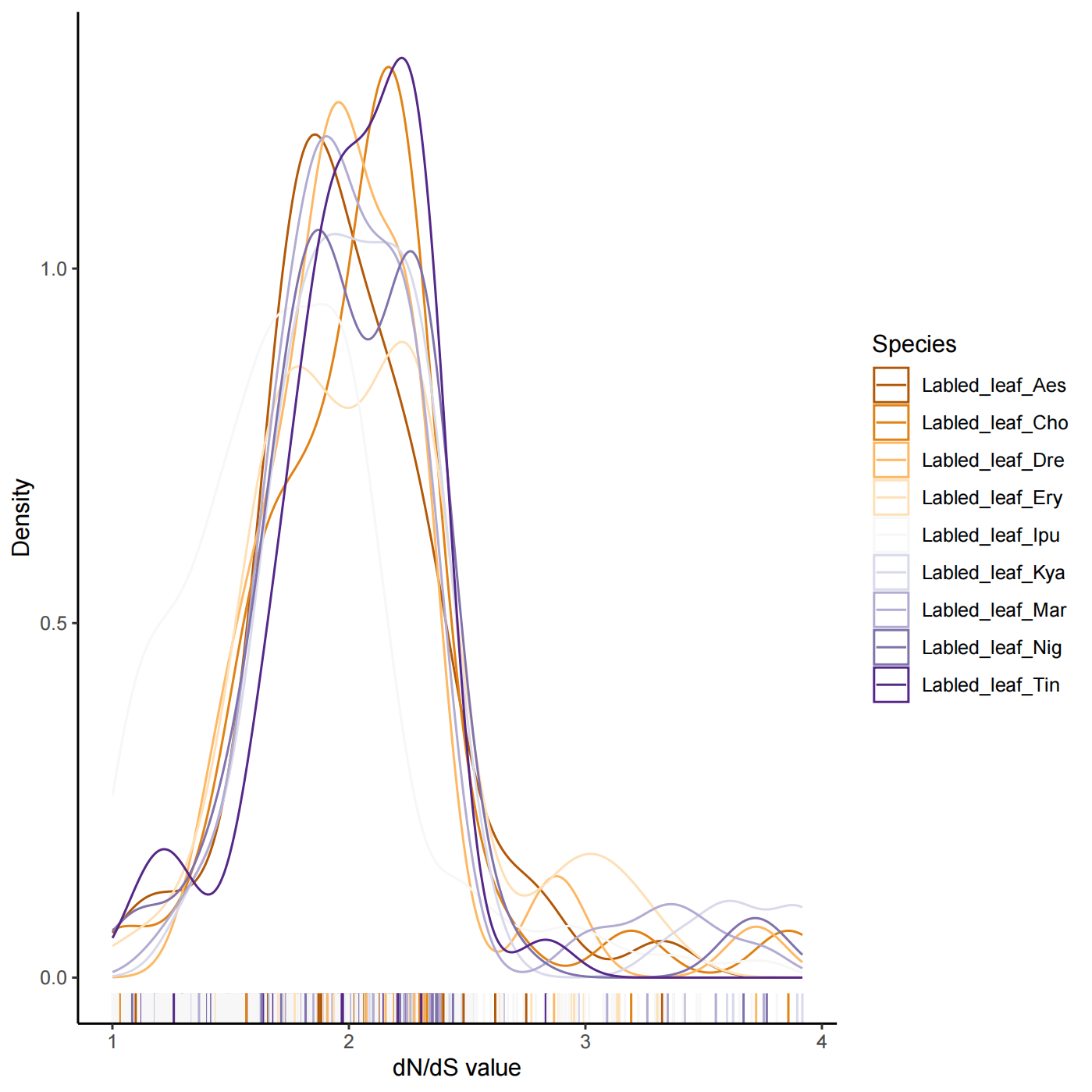

**Figure S23 | Density curves of the dN/dS value of all positive selected genes at each leaf.**

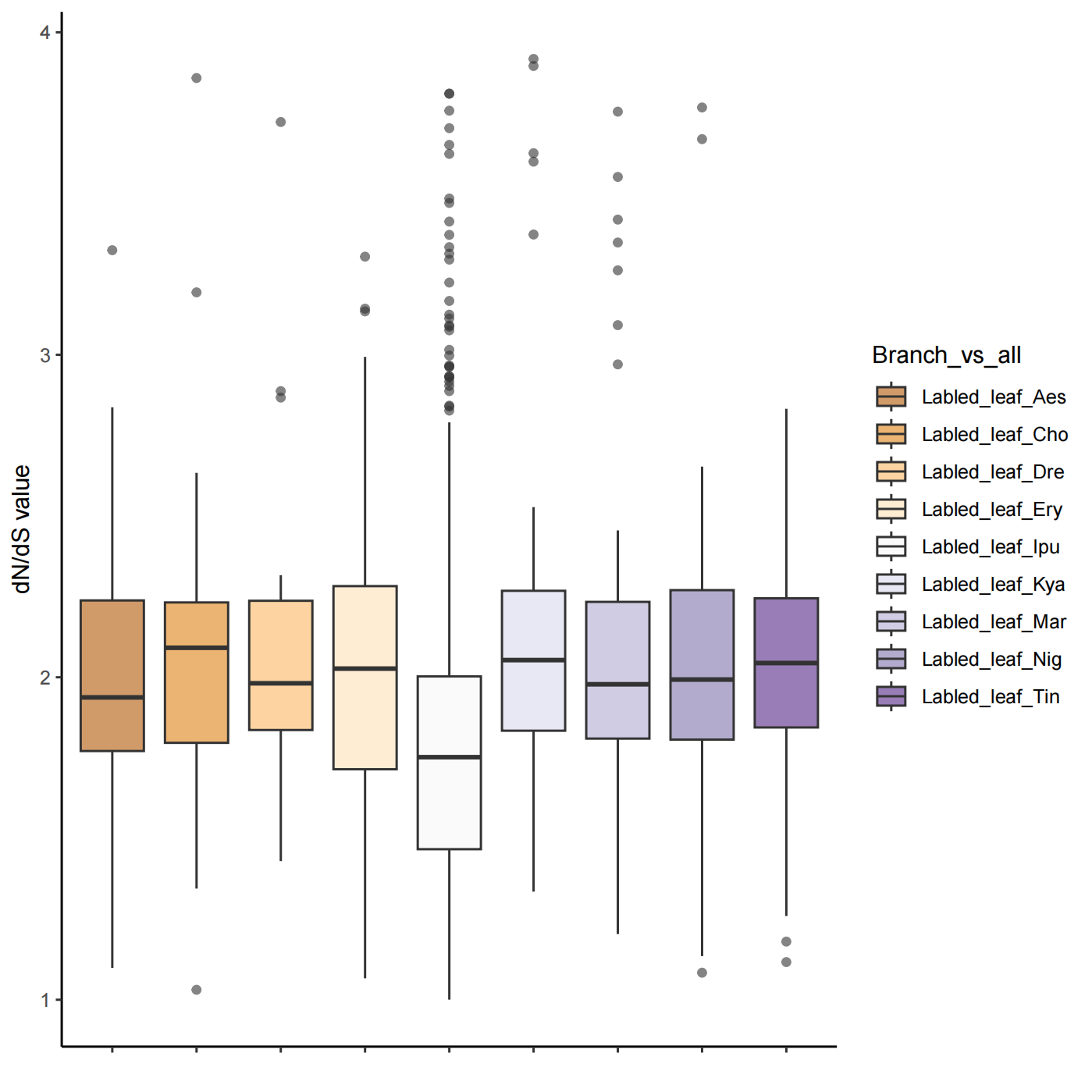

**Figure S24 | Box plot of the dN/dS value of all positive selected genes at each leaf.**

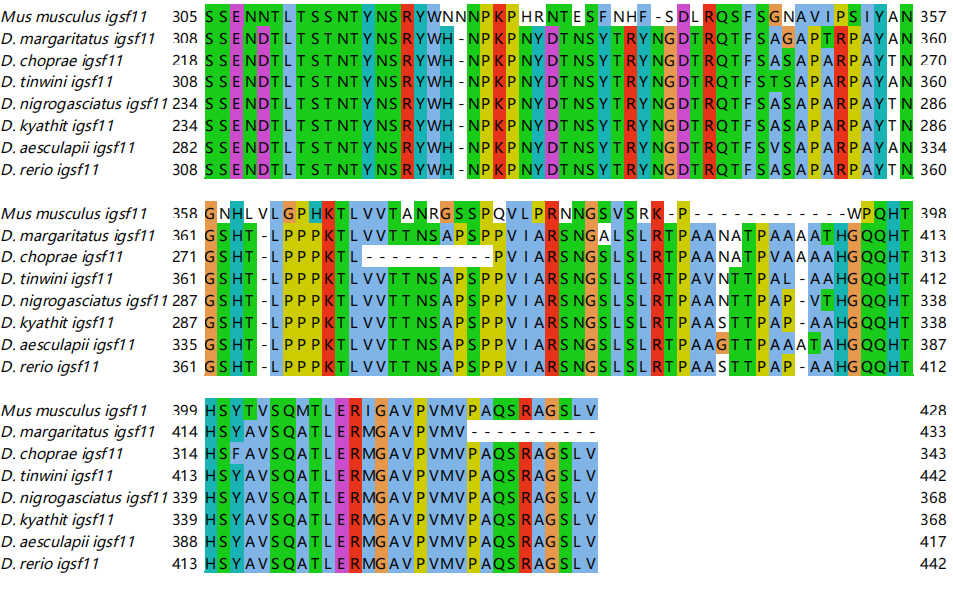

**Figure S25 | Amino acid alignment of the terminal region of Igsf11.**

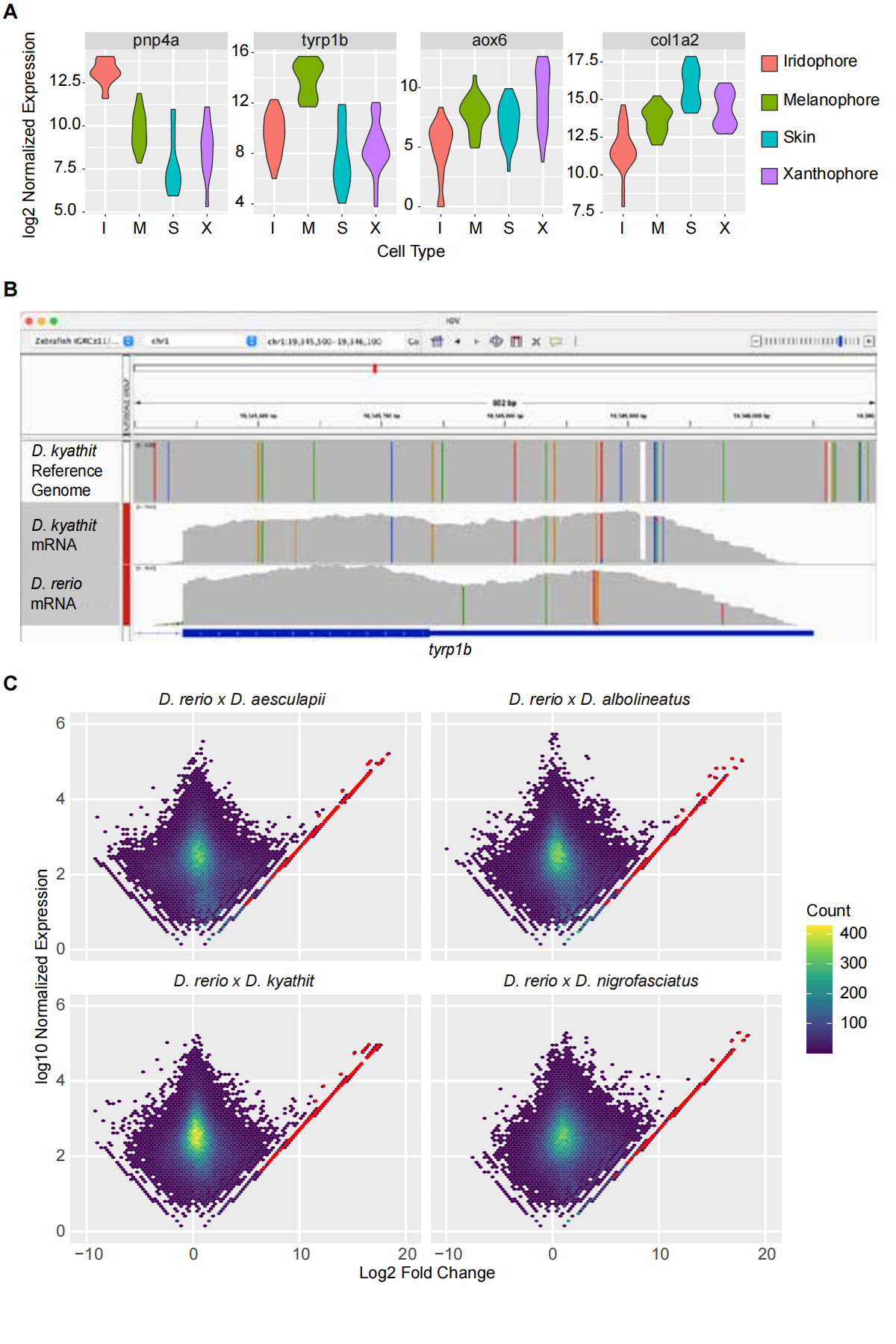

**Figure S26 | Allele-specific RNA-seq results recapitulate known biology.** A) FACS-sorted populations show increased expression of known marker genes. B) RNA-seq reads from *D. rerio* x *D. kyathit* hybrid melanophores show SNPs and an indel identified in the *D. kyathit* genome (top) are present in the RNA-seq reads from assigned to the *D. kyathit* allele (middle), but not the *D. rerio* allele (bottom). C) Density plot of allelic bias across all samples (*D. rerio* bias indicated as positive). Most genes display limited allelic bias as evidenced by the density of points near the center. Bins with maternally-expressed mitochondrial genes are shown in red.

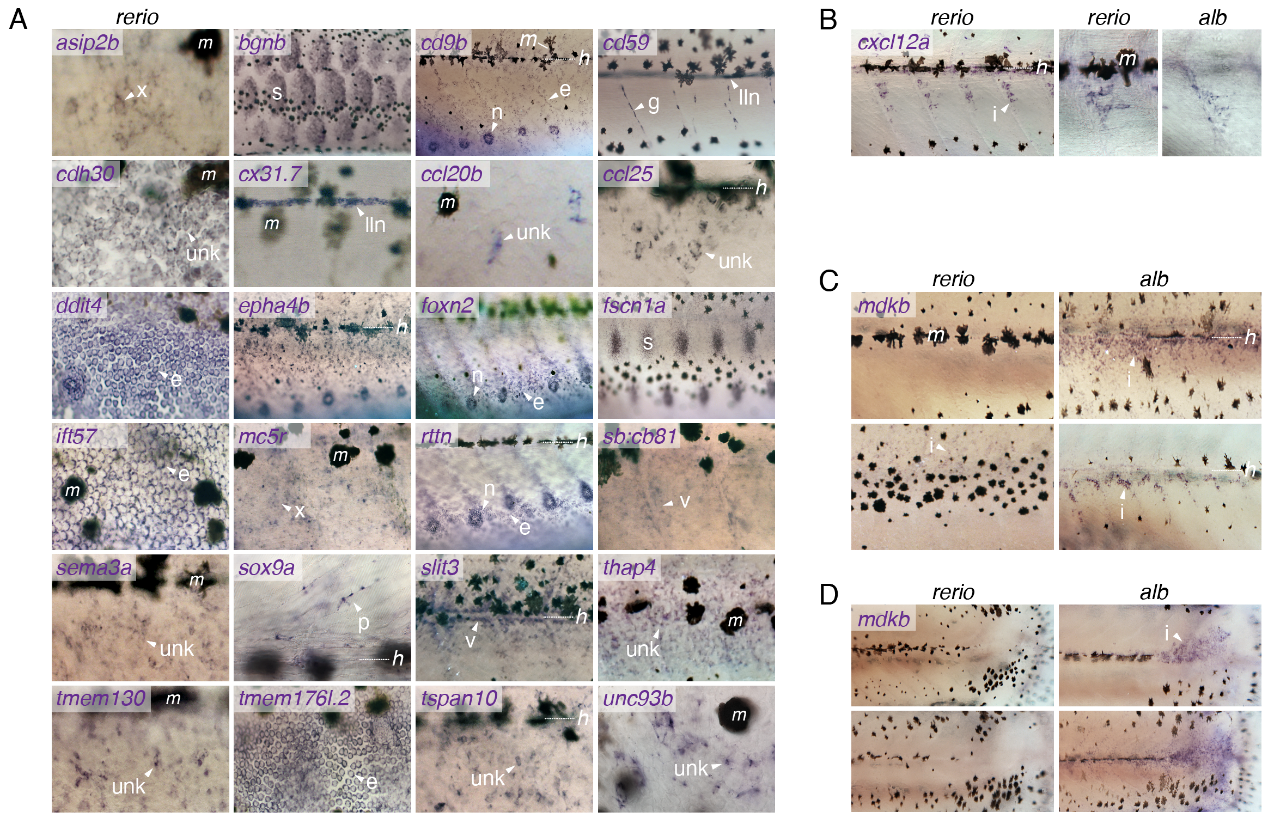

**Figure S27 | *In situ* hybridization of genes with differential allelic expression in hybrids.** A) Screen for gene expression during larva-to-adult transformation in *D. rerio* (6.2–8.5 mm standard length). A variety of presumptive cell types and tissues were observed across loci: e, epidermal cell; g, glia; I, iridophore; lln, lateral line nerve; n, neuromast; p, neural crest derived progenitor; x, xanthophore; unk, unknown; v, vasculature. m, indicates melanized melanophores (unstained) and ---h indicates level of horizontal myoseptum when present. B) Expression of *cxcl12a* in the prospective light interstripe region of *D. rerio* (left) and at higher magnification in *D. rerio* and *D. albolineatus* (right). C) Expression of *mdkb* in presumptive iridophores. At the same stage of the larva-to-adult transformation, staining is intense in *D. albolineatus* but not detected in *D. rerio* (upper). Later, weak staining was observed in a subset of *D. rerio* iridophores (lower left); staining was apparent in *D. albolineatus* at early stages as well (lower right). D) *mdkb* expression in region of iridophores at caudal penduncle in *D. albolineatus* but not *D. rerio* at early (upper) and later (lower) stages of the larva-to-adult transformation.

**SM Tables**

**Table S1. An overview of variants called from 14 *Danio* genomes.**

| **Species/Strain** | **SNPs** | **Indels** | **PAVs (total length, Mb)** | **PAVs (nubmer)** |
| --- | --- | --- | --- | --- |
| *Danio rerio*（*AB*） | 14,008,956 | 3,264,862 | 310.34 | 88,912 |
| *Danio rerio*（*NA*） | 14,743,056 | 3,387,552 | 215.62 | 99,873 |
| *Danio rerio*（*CB*） | 19,234,281 | 4,346,281 | 249.43 | 111,529 |
| *Danio aesculapii* | 13,718,792 | 3,196,155 | 1036.19 | 20,480 |
| *Danio kyathit* | 14,838,116 | 3,526,451 | 1036.79 | 22,352 |
| *Danio tinwini* | 11,519,329 | 2,684,451 | 994.96 | 115,518 |
| *Danio nigrofasciatus* | 16,190,093 | 4,130,600 | 969.42 | 14,249 |
| *Danio albolineatus* | 9,777,039 | 2,313,604 | 975.94 | 13,651 |
| *Danio jaintianensis* | 9,376,249 | 2,293,119 | 964.51 | 13,360 |
| *Danio choprae* | 10,639,665 | 2,569,840 | 929.91 | 11,884 |
| *Danio margaritatus* | 13,069,079 | 3,471,552 | 888.31 | 9,043 |
| *Danio erythromicron* | 12,945,277 | 3,459,055 | 625.78 | 8,876 |
| *Danionella cerebrum* | 1,014,099 | 414,496 | 562.07 | 758 |
| *Danionella dracula* | 1,426,272 | 465,956 | 705.50 | 864 |

**Table S2 | Frequency of ASTRAL region trees with different topologies among different positions of chromosomes, with a total of 14 chromosomes.** 50 % - 10 % represents the position from chromosome center to end. Type1, (((*D. rerio*, *D. kyathit*), *D. aesculapii*), (*D. tinwini*, *D. nigrofasciatus*)); Type2, (((*D. rerio*, *D. aesculapii*), *D. kyathit*), (*D. tinwini*, *D. nigrofasciatus*)); Type3, ((*D. rerio*, *D. aesculapii*), (*D. kyathit*, (*D. tinwini*, *D. nigrofasciatus*))); Type4, ((*D. rerio*, (*D. aesculapii*, *D. kyathit*)), (*D. tinwini*, *D. nigrofasciatus*)); Type5, ((*D. rerio*, *D. aesculapii*), ((*D. kyathit*, *D. tinwini*), *D. nigrofasciatus*)).

| **Frequency (%)** | **Type1** | **Type2** | **Type3** | **Type4** | **Type5** | **Others** | **Total** |
| --- | --- | --- | --- | --- | --- | --- | --- |
| Genome | 37.14 | 18.57 | 12.86 | 11.43 | 5.71 | 14.29 | 100 |
| 50 | 78.57 | 0 | 0 | 21.43 | 0 | 0 | 100 |
| 40 | 57.14 | 21.43 | 7.14 | 7.14 | 0 | 7.14 | 100 |
| 30 | 21.43 | 50 | 0 | 21.43 | 0 | 7.14 | 100 |
| 20 | 28.57 | 21.43 | 21.43 | 7.14 | 0 | 21.43 | 100 |
| 10 | 0 | 0 | 35.71 | 0 | 28.57 | 35.71 | 100 |

**Table S3 | Frequency of ASTRAL region trees with different topologies among different positions of chromosomes, with a total of 11 chromosomes.** 50 % - 10 % represents the position from chromosome center to end. Type1, (((*D. rerio*, *D. kyathit*), *D. aesculapii*), (*D. tinwini*, *D. nigrofasciatus*)); Type2, (((*D. rerio*, *D. aesculapii*), *D. kyathit*), (*D. tinwini*, *D. nigrofasciatus*)); Type3, ((*D. rerio*, *D. aesculapii*), (*D. kyathit*, (*D. tinwini*, *D. nigrofasciatus*))); Type4, ((*D. rerio*, (*D. aesculapii*, *D. kyathit*)), (*D. tinwini*, *D. nigrofasciatus*)); Type5, ((*D. rerio*, *D. aesculapii*), ((*D. kyathit*, *D. tinwini*), *D. nigrofasciatus*)).

| **Frequency (%)** | **Type1** | **Type2** | **Type3** | **Type4** | **Type5** | **Others** | **Total** |
| --- | --- | --- | --- | --- | --- | --- | --- |
| Genome | 40 | 20 | 16.36 | 7.27 | 3.64 | 12.73 | 100 |
| 50 | 100 | 0 | 0 | 0 | 0 | 0 | 100 |
| 40 | 54.55 | 27.27 | 9.09 | 9.09 | 0 | 0 | 100 |
| 30 | 18.18 | 45.45 | 0 | 27.27 | 0 | 9.09 | 100 |
| 20 | 27.27 | 27.27 | 27.27 | 0 | 0 | 18.18 | 100 |
| 10 | 0 | 0 | 45.45 | 0 | 18.18 | 36.36 | 100 |

**Table S4 | Frequency of ASTRAL region trees with different topologies among different positions of chromosomes, with a total of 11 chromosomes.** 50 % - 10 % represents the position from chromosome center to end. Type1, ((*D. rerio*, *D. kyathit*), *D. aesculapii*); Type2, ((*D. rerio*, *D. aesculapii*), *D. kyathit*); Type3, (*D. rerio*, (*D. aesculapii*, *D. kyathit*)).

| **Frequency (%)** | **Type1** | **Type2** | **Type3** | **Total** |
| --- | --- | --- | --- | --- |
| genome | 47.27 | 40 | 12.73 | 100 |
| 50 | 100 | 0 | 0 | 100 |
| 40 | 54.55 | 36.36 | 9.09 | 100 |
| 30 | 18.18 | 45.45 | 36.36 | 100 |
| 20 | 45.45 | 54.55 | 0 | 100 |
| 10 | 18.18 | 63.64 | 18.18 | 100 |

**Table S5 | Introgression analysis of *D. rerio* subclade using D_FOIL_.** Five groups with symmetrical five-taxon topology are used: (1) (((*D. tinwini, D. nigrofasciatus*), (*D. rerio*, *D. aesculapii*)), *D. albolineatus*); (2)(( (*D. tinwini*, *D. nigrofasciatus*), (*D. rerio*, *D. kyathit*)), *D. albolineatus*); (3) (((*D. tinwini*, *D. nigrofasciatus*), (*D. aesculapii*, *D. kyathit*)), *D. albolineatus*); (4) (((*D. rerio*, *D. aesculapii*), (*D. kyathit*, *D. tinwini*)), *D. albolineatus*);(5) (((*D. rerio*, *D. aesculapii*), (*D. kyathit*, *D. nigrofasciatus*)), *D. albolineatus*).

| **Group_id** | **Donor** | **Acceptor** | **Introgressed windows** |
| --- | --- | --- | --- |
| Group1 | *D. tinwini + D. nigrofasciatus* | *D. aesculapii* | 23 |
| Group1 | *D. aesculapii* | *D. tinwini + D. nigrofasciatus* | 23 |
| Group1 | *D. tinwini + D. nigrofasciatus* | *D. rerio* | 4 |
| Group1 | *D. rerio* | *D. tinwini + D. nigrofasciatus* | 4 |
| Group2 | *D. tinwini + D. nigrofasciatus* | *D. kyathit* | 121 |
| Group2 | *D. kyathit* | *D. tinwini + D. nigrofasciatus* | 121 |
| Group2 | *D. tinwini* | *D. kyathit* | 7 |
| Group2 | *D. kyathit* | *D. nigrofasciatus* | 1 |
| Group2 | *D. tinwini + D. nigrofasciatus* | *D. rerio* | 1 |
| Group2 | *D. rerio* | *D. tinwini + D. nigrofasciatus* | 1 |
| Group3 | *D. tinwini + D. nigrofasciatus* | *D. kyathit* | 75 |
| Group3 | *D. kyathit* | *D. tinwini + D. nigrofasciatus* | 75 |
| Group3 | *D. tinwini* | *D. kyathit* | 3 |
| Group3 | *D. tinwini + D. nigrofasciatus* | *D. aesculapii* | 4 |
| Group3 | *D. aesculapii* | *D. tinwini + D. nigrofasciatus* | 4 |
| Group3 | *D. nigrofasciatus* | *D. kyathit* | 1 |
| Group4 | *D. rerio + D. aesculapii* | *D. kyathit* | 49 |
| Group4 | *D. kyathit* | *D. rerio + D. aesculapii* | 49 |
| Group4 | *D. kyathit* | *D. rerio* | 1 |
| Group4 | *D. kyathit* | *D. aesculapii* | 1 |
| Group5 | *D. rerio + D. aesculapii* | *D. kyathit* | 68 |
| Group5 | *D. kyathit* | *D. rerio + D. aesculapii* | 68 |
| Group5 | *D. kyathit* | *D. aesculapii* | 3 |
| Group5 | *D. aesculapii* | *D. kyathit* | 2 |

**Table S6 | Introgression analysis of *Danio* species using D_FOIL_.** Three combinations (P3, P4) are used: (1) (((P1, P2), (*D. tinwini*, *D. nigrofasciatus*)), *L. oculatus*); (2) (((P1, P2), (*D. erythromicron*, *D. margaritatus*)), *L. oculatus*); (3) (((P1, P2), (*D. choprae*, *D. jaintianensis*)), *L. oculatus*).

| **Group_id** | **P1** | **P2** | **Donor** | **Acceptor** | **Introgressed**  **windows** |
| --- | --- | --- | --- | --- | --- |
| group1 | *D. rerio* | *D. aesculapii* | *D. rerio + D. aesculapii* | *D. nigrofasciatus* | 5 |
| group1 | *D. rerio* | *D. aesculapii* | *D. rerio* | *D. nigrofasciatus* | 1 |
| group1 | *D. rerio* | *D. aesculapii* | *D. rerio + D. aesculapii* | *D. tinwini* | 6 |
| group1 | *D. rerio* | *D. kyathit* | *D. rerio + D. kyathit* | *D. nigrofasciatus* | 6 |
| group1 | *D. rerio* | *D. kyathit* | *D. tinwini* | *D. kyathit* | 2 |
| group1 | *D. rerio* | *D. kyathit* | *D. rerio + D. kyathit* | *D. tinwini* | 6 |
| group1 | *D. rerio* | *D. kyathit* | *D. kyathit* | *D. tinwini* | 1 |
| group1 | *D. rerio* | *D. albolineatus* | *D. rerio* | *D. nigrofasciatus* | 2 |
| group1 | *D. rerio* | *D. albolineatus* | *D. rerio + D. albolineatus* | *D. nigrofasciatus* | 3 |
| group1 | *D. rerio* | *D. albolineatus* | *D. rerio + D. albolineatus* | *D. tinwini* | 3 |
| group1 | *D. rerio* | *D. albolineatus* | *D. tinwini* | *D. rerio* | 1 |
| group1 | *D. rerio* | *D. erythromicron* | *D. rerio + D. erythromicron* | *D. nigrofasciatus* | 5 |
| group1 | *D. rerio* | *D. erythromicron* | *D. rerio + D. erythromicron* | *D. tinwini* | 4 |
| group1 | *D. rerio* | *D. erythromicron* | *D. tinwini* | *D. rerio* | 1 |
| group1 | *D. rerio* | *D. margaritatus* | *D. rerio + D. margaritatus* | *D. nigrofasciatus* | 6 |
| group1 | *D. rerio* | *D. margaritatus* | *D. rerio* | *D. nigrofasciatus* | 1 |
| group1 | *D. rerio* | *D. margaritatus* | *D. tinwini* | *D. rerio* | 1 |
| group1 | *D. rerio* | *D. margaritatus* | *D. rerio + D. margaritatus* | *D. tinwini* | 3 |
| group1 | *D. rerio* | *D. choprae* | *D. rerio + D. choprae* | *D. nigrofasciatus* | 5 |
| group1 | *D. rerio* | *D. choprae* | *D. rerio + D. choprae* | *D. tinwini* | 4 |
| group1 | *D. rerio* | *D. jaintianensis* | *D. rerio + D. jaintianensis* | *D. nigrofasciatus* | 8 |
| group1 | *D. rerio* | *D. jaintianensis* | *D. rerio + D. jaintianensis* | *D. tinwini* | 3 |
| group1 | *D. aesculapii* | *D. albolineatus* | *D. aesculapii* | *D. nigrofasciatus* | 2 |
| group1 | *D. aesculapii* | *D. albolineatus* | *D. aesculapii + D. albolineatus* | *D. nigrofasciatus* | 4 |
| group1 | *D. aesculapii* | *D. albolineatus* | *D. tinwini* | *D. aesculapii* | 2 |
| group1 | *D. aesculapii* | *D. albolineatus* | *D. aesculapii + D. albolineatus* | *D. tinwini* | 4 |
| group1 | *D. aesculapii* | *D. erythromicron* | *D. aesculapii + D. erythromicron* | *D. nigrofasciatus* | 2 |
| group1 | *D. aesculapii* | *D. erythromicron* | *D. aesculapii* | *D. nigrofasciatus* | 3 |
| group1 | *D. aesculapii* | *D. erythromicron* | *D. tinwini* | *D. aesculapii* | 1 |
| group1 | *D. aesculapii* | *D. erythromicron* | *D. aesculapii + D. erythromicron* | *D. tinwini* | 3 |
| group1 | *D. aesculapii* | *D. erythromicron* | *D. nigrofasciatus* | *D. aesculapii* | 2 |
| group1 | *D. aesculapii* | *D. margaritatus* | *D. tinwini* | *D. aesculapii* | 1 |
| group1 | *D. aesculapii* | *D. margaritatus* | *D. aesculapii + D. margaritatus* | *D. nigrofasciatus* | 5 |
| group1 | *D. aesculapii* | *D. margaritatus* | *D. nigrofasciatus* | *D. aesculapii* | 1 |
| group1 | *D. aesculapii* | *D. margaritatus* | *D. aesculapii + D. margaritatus* | *D. tinwini* | 1 |
| group1 | *D. aesculapii* | *D. margaritatus* | *D. aesculapii* | *D. nigrofasciatus* | 1 |
| group1 | *D. aesculapii* | *D. choprae* | *D. aesculapii + D. choprae* | *D. nigrofasciatus* | 5 |
| group1 | *D. aesculapii* | *D. choprae* | *D. aesculapii + D. choprae* | *D. tinwini* | 4 |
| group1 | *D. aesculapii* | *D. jaintianensis* | *D. aesculapii + D. jaintianensis* | *D. nigrofasciatus* | 7 |
| group1 | *D. aesculapii* | *D. jaintianensis* | *D. aesculapii + D. jaintianensis* | *D. tinwini* | 4 |
| group1 | *D. kyathit* | *D. aesculapii* | *D. kyathit + D. aesculapii* | *D. nigrofasciatus* | 5 |
| group1 | *D. kyathit* | *D. aesculapii* | *D. tinwini* | *D. kyathit* | 3 |
| group1 | *D. kyathit* | *D. aesculapii* | *D. kyathit + D. aesculapii* | *D. tinwini* | 5 |
| group1 | *D. kyathit* | *D. albolineatus* | *D. kyathit + D. albolineatus* | *D. tinwini* | 4 |
| group1 | *D. kyathit* | *D. albolineatus* | *D. kyathit + D. albolineatus* | *D. nigrofasciatus* | 4 |
| group1 | *D. kyathit* | *D. albolineatus* | *D. tinwini* | *D. kyathit* | 4 |
| group1 | *D. kyathit* | *D. erythromicron* | *D. kyathit* | *D. nigrofasciatus* | 3 |
| group1 | *D. kyathit* | *D. erythromicron* | *D. tinwini* | *D. kyathit* | 9 |
| group1 | *D. kyathit* | *D. erythromicron* | *D. kyathit + D. erythromicron* | *D. nigrofasciatus* | 3 |
| group1 | *D. kyathit* | *D. erythromicron* | *D. kyathit + D. erythromicron* | *D. tinwini* | 3 |
| group1 | *D. kyathit* | *D. erythromicron* | *D. nigrofasciatus* | *D. kyathit* | 1 |
| group1 | *D. kyathit* | *D. margaritatus* | *D. kyathit* | *D. nigrofasciatus* | 3 |
| group1 | *D. kyathit* | *D. margaritatus* | *D. margaritatus* | *D. kyathit* | 8 |
| group1 | *D. kyathit* | *D. margaritatus* | *D. kyathit + D. margaritatus* | *D. nigrofasciatus* | 2 |
| group1 | *D. kyathit* | *D. margaritatus* | *D. kyathit + D. margaritatus* | *D. tinwini* | 2 |
| group1 | *D. kyathit* | *D. choprae* | *D. tinwini* | *D. kyathit* | 5 |
| group1 | *D. kyathit* | *D. choprae* | *D. kyathit + D. choprae* | *D. nigrofasciatus* | 2 |
| group1 | *D. kyathit* | *D. choprae* | *D. kyathit + D. choprae* | *D. tinwini* | 1 |
| group1 | *D. kyathit* | *D. jaintianensis* | *D. kyathit + D. jaintianensis* | *D. nigrofasciatus* | 4 |
| group1 | *D. kyathit* | *D. jaintianensis* | *D. tinwini* | *D. kyathit* | 4 |
| group1 | *D. kyathit* | *D. jaintianensis* | *D. kyathit + D. jaintianensis* | *D. tinwini* | 5 |
| group1 | *D. albolineatus* | *D. erythromicron* | *D. albolineatus + D. erythromicron* | *D. nigrofasciatus* | 7 |
| group1 | *D. albolineatus* | *D. erythromicron* | *D. albolineatus* | *D. nigrofasciatus* | 3 |
| group1 | *D. albolineatus* | *D. erythromicron* | *D. albolineatus + D. erythromicron* | *D. tinwini* | 3 |
| group1 | *D. albolineatus* | *D. margaritatus* | *D. albolineatus* | *D. nigrofasciatus* | 4 |
| group1 | *D. albolineatus* | *D. margaritatus* | *D. albolineatus + D. margaritatus* | *D. tinwini* | 3 |
| group1 | *D. albolineatus* | *D. margaritatus* | *D. albolineatus + D. margaritatus* | *D. nigrofasciatus* | 2 |
| group1 | *D. albolineatus* | *D. choprae* | *D. albolineatus + D. choprae* | *D. tinwini* | 3 |
| group1 | *D. albolineatus* | *D. choprae* | *D. albolineatus + D. choprae* | *D. nigrofasciatus* | 3 |
| group1 | *D. albolineatus* | *D. choprae* | *D. tinwini* | *D. choprae* | 1 |
| group1 | *D. albolineatus* | *D. choprae* | *D. choprae* | *D. tinwini* | 1 |
| group1 | *D. albolineatus* | *D. jaintianensis* | *D. albolineatus + D. jaintianensis* | *D. nigrofasciatus* | 7 |
| group1 | *D. albolineatus* | *D. jaintianensis* | *D. albolineatus + D. jaintianensis* | *D. tinwini* | 5 |
| group1 | *D. albolineatus* | *D. jaintianensis* | *D. jaintianensis* | *D. tinwini* | 1 |
| group1 | *D. erythromicron* | *D. margaritatus* | *D. erythromicron + D. margaritatus* | *D. nigrofasciatus* | 8 |
| group1 | *D. erythromicron* | *D. margaritatus* | *D. erythromicron + D. margaritatus* | *D. tinwini* | 6 |
| group1 | *D. erythromicron* | *D. choprae* | *D. erythromicron + D. choprae* | *D. nigrofasciatus* | 7 |
| group1 | *D. erythromicron* | *D. choprae* | *D. erythromicron + D. choprae* | *D. tinwini* | 7 |
| group1 | *D. erythromicron* | *D. jaintianensis* | *D. erythromicron + D. jaintianensis* | *D. nigrofasciatus* | 9 |
| group1 | *D. erythromicron* | *D. jaintianensis* | *D. erythromicron + D. jaintianensis* | *D. tinwini* | 9 |
| group1 | *D. erythromicron* | *D. jaintianensis* | *D. tinwini* | *D. jaintianensis* | 1 |
| group1 | *D. margaritatus* | *D. choprae* | *D. margaritatus + D. choprae* | *D. nigrofasciatus* | 4 |
| group1 | *D. margaritatus* | *D. choprae* | *D. margaritatus + D. choprae* | *D. tinwini* | 4 |
| group1 | *D. margaritatus* | *D. jaintianensis* | *D. margaritatus + D. jaintianensis* | *D. tinwini* | 6 |
| group1 | *D. margaritatus* | *D. jaintianensis* | *D. margaritatus + D. jaintianensis* | *D. nigrofasciatus* | 3 |
| group1 | *D. margaritatus* | *D. jaintianensis* | *D. nigrofasciatus* | *D. jaintianensis* | 1 |
| group1 | *D. choprae* | *D. jaintianensis* | *D. choprae + D. jaintianensis* | *D. nigrofasciatus* | 6 |
| group1 | *D. choprae* | *D. jaintianensis* | *D. choprae + D. jaintianensis* | *D. tinwini* | 6 |
| group1 | *D. choprae* | *D. jaintianensis* | *D. tinwini* | *D. choprae* | 1 |
| group2 | *D. rerio* | *D. aesculapii* | *D. rerio + D. aesculapii* | *D. erythromicron* | 1 |
| group2 | *D. rerio* | *D. aesculapii* | *D. rerio + D. aesculapii* | *D. margaritatus* | 6 |
| group2 | *D. rerio* | *D. kyathit* | *D. rerio + D. kyathit* | *D. margaritatus* | 7 |
| group2 | *D. rerio* | *D. kyathit* | *D. rerio + D. kyathit* | *D. erythromicron* | 2 |
| group2 | *D. rerio* | *D. tinwini* | *D. rerio + D. tinwini* | *D. margaritatus* | 4 |
| group2 | *D. rerio* | *D. tinwini* | *D. rerio + D. tinwini* | *D. erythromicron* | 2 |
| group2 | *D. rerio* | *D. nigrofasciatus* | *D. rerio + D. nigrofasciatus* | *D. margaritatus* | 2 |
| group2 | *D. rerio* | *D. nigrofasciatus* | *D. rerio + D. nigrofasciatus* | *D. erythromicron* | 1 |
| group2 | *D. rerio* | *D. albolineatus* | *D. rerio + D. albolineatus* | *D. erythromicron* | 2 |
| group2 | *D. rerio* | *D. albolineatus* | *D. rerio + D. albolineatus* | *D. margaritatus* | 6 |
| group2 | *D. rerio* | *D. choprae* | *D. rerio + D. choprae* | *D. margaritatus* | 3 |
| group2 | *D. rerio* | *D. choprae* | *D. rerio + D. choprae* | *D. erythromicron* | 1 |
| group2 | *D. rerio* | *D. jaintianensis* | *D. rerio + D. jaintianensis* | *D. margaritatus* | 4 |
| group2 | *D. rerio* | *D. jaintianensis* | *D. rerio + D. jaintianensis* | *D. erythromicron* | 2 |
| group2 | *D. aesculapii* | *D. tinwini* | *D. aesculapii + D. tinwini* | *D. margaritatus* | 3 |
| group2 | *D. aesculapii* | *D. tinwini* | *D. aesculapii + D. tinwini* | *D. erythromicron* | 1 |
| group2 | *D. aesculapii* | *D. tinwini* | *D. aesculapii* | *D. margaritatus* | 2 |
| group2 | *D. aesculapii* | *D. nigrofasciatus* | *D. aesculapii + D. nigrofasciatus* | *D. erythromicron* | 1 |
| group2 | *D. aesculapii* | *D. nigrofasciatus* | *D. aesculapii + D. nigrofasciatus* | *D. margaritatus* | 3 |
| group2 | *D. aesculapii* | *D. nigrofasciatus* | *D. aesculapii* | *D. margaritatus* | 1 |
| group2 | *D. aesculapii* | *D. albolineatus* | *D. aesculapii + D. albolineatus* | *D. margaritatus* | 5 |
| group2 | *D. aesculapii* | *D. albolineatus* | *D. aesculapii + D. albolineatus* | *D. erythromicron* | 2 |
| group2 | *D. aesculapii* | *D. albolineatus* | *D. aesculapii* | *D. margaritatus* | 1 |
| group2 | *D. aesculapii* | *D. choprae* | *D. aesculapii + D. choprae* | *D. margaritatus* | 5 |
| group2 | *D. aesculapii* | *D. choprae* | *D. aesculapii + D. choprae* | *D. erythromicron* | 1 |
| group2 | *D. aesculapii* | *D. jaintianensis* | *D. aesculapii + D. jaintianensis* | *D. margaritatus* | 8 |
| group2 | *D. aesculapii* | *D. jaintianensis* | *D. aesculapii + D. jaintianensis* | *D. erythromicron* | 1 |
| group2 | *D. aesculapii* | *D. jaintianensis* | *D. jaintianensis* | *D. erythromicron* | 1 |
| group2 | *D. kyathit* | *D. aesculapii* | *D. kyathit + D. aesculapii* | *D. margaritatus* | 6 |
| group2 | *D. kyathit* | *D. aesculapii* | *D. kyathit + D. aesculapii* | *D. erythromicron* | 1 |
| group2 | *D. kyathit* | *D. tinwini* | *D. kyathit + D. tinwini* | *D. margaritatus* | 8 |
| group2 | *D. kyathit* | *D. tinwini* | *D. kyathit + D. tinwini* | *D. erythromicron* | 1 |
| group2 | *D. kyathit* | *D. nigrofasciatus* | *D. kyathit + D. nigrofasciatus* | *D. erythromicron* | 2 |
| group2 | *D. kyathit* | *D. nigrofasciatus* | *D. kyathit + D. nigrofasciatus* | *D. margaritatus* | 4 |
| group2 | *D. kyathit* | *D. nigrofasciatus* | *D. margaritatus* | *D. kyathit* | 1 |
| group2 | *D. kyathit* | *D. nigrofasciatus* | *D. kyathit* | *D. margaritatus* | 2 |
| group2 | *D. kyathit* | *D. albolineatus* | *D. kyathit* | *D. margaritatus* | 1 |
| group2 | *D. kyathit* | *D. albolineatus* | *D. kyathit + D. albolineatus* | *D. margaritatus* | 5 |
| group2 | *D. kyathit* | *D. albolineatus* | *D. kyathit + D. albolineatus* | *D. erythromicron* | 1 |
| group2 | *D. kyathit* | *D. choprae* | *D. kyathit + D. choprae* | *D. margaritatus* | 4 |
| group2 | *D. kyathit* | *D. choprae* | *D. kyathit + D. choprae* | *D. erythromicron* | 1 |
| group2 | *D. kyathit* | *D. jaintianensis* | *D. kyathit + D. jaintianensis* | *D. margaritatus* | 10 |
| group2 | *D. kyathit* | *D. jaintianensis* | *D. kyathit + D. jaintianensis* | *D. erythromicron* | 2 |
| group2 | *D. tinwini* | *D. nigrofasciatus* | *D. tinwini + D. nigrofasciatus* | *D. margaritatus* | 7 |
| group2 | *D. tinwini* | *D. nigrofasciatus* | *D. margaritatus* | *D. tinwini* | 1 |
| group2 | *D. tinwini* | *D. choprae* | *D. tinwini + D. choprae* | *D. margaritatus* | 4 |
| group2 | *D. tinwini* | *D. jaintianensis* | *D. tinwini + D. jaintianensis* | *D. margaritatus* | 5 |
| group2 | *D. tinwini* | *D. jaintianensis* | *D. tinwini + D. jaintianensis* | *D. erythromicron* | 2 |
| group2 | *D. nigrofasciatus* | *D. choprae* | *D. margaritatus* | *D. choprae* | 1 |
| group2 | *D. nigrofasciatus* | *D. choprae* | *D. nigrofasciatus + D. choprae* | *D. margaritatus* | 2 |
| group2 | *D. nigrofasciatus* | *D. choprae* | *D. nigrofasciatus + D. choprae* | *D. erythromicron* | 1 |
| group2 | *D. nigrofasciatus* | *D. jaintianensis* | *D. nigrofasciatus + D. jaintianensis* | *D. margaritatus* | 5 |
| group2 | *D. nigrofasciatus* | *D. jaintianensis* | *D. nigrofasciatus + D. jaintianensis* | *D. erythromicron* | 2 |
| group2 | *D. nigrofasciatus* | *D. jaintianensis* | *D. margaritatus* | *D. jaintianensis* | 2 |
| group2 | *D. albolineatus* | *D. tinwini* | *D. albolineatus + D. tinwini* | *D. margaritatus* | 8 |
| group2 | *D. albolineatus* | *D. tinwini* | *D. albolineatus + D. tinwini* | *D. erythromicron* | 1 |
| group2 | *D. albolineatus* | *D. nigrofasciatus* | *D. albolineatus + D. nigrofasciatus* | *D. erythromicron* | 2 |
| group2 | *D. albolineatus* | *D. nigrofasciatus* | *D. albolineatus + D. nigrofasciatus* | *D. margaritatus* | 5 |
| group2 | *D. albolineatus* | *D. choprae* | *D. choprae* | *D. margaritatus* | 1 |
| group2 | *D. albolineatus* | *D. choprae* | *D. albolineatus + D. choprae* | *D. erythromicron* | 1 |
| group2 | *D. albolineatus* | *D. choprae* | *D. albolineatus + D. choprae* | *D. margaritatus* | 6 |
| group2 | *D. albolineatus* | *D. jaintianensis* | *D. albolineatus + D. jaintianensis* | *D. erythromicron* | 1 |
| group2 | *D. albolineatus* | *D. jaintianensis* | *D. albolineatus + D. jaintianensis* | *D. margaritatus* | 6 |
| group2 | *D. choprae* | *D. jaintianensis* | *D. choprae + D. jaintianensis* | *D. margaritatus* | 5 |
| group2 | *D. choprae* | *D. jaintianensis* | *D. choprae + D. jaintianensis* | *D. erythromicron* | 2 |
| group3 | *D. rerio* | *D. aesculapii* | *D. rerio + D. aesculapii* | *D. jaintianensis* | 8 |
| group3 | *D. rerio* | *D. aesculapii* | *D. rerio + D. aesculapii* | *D. choprae* | 2 |
| group3 | *D. rerio* | *D. kyathit* | *D. rerio + D. kyathit* | *D. jaintianensis* | 12 |
| group3 | *D. rerio* | *D. kyathit* | *D. rerio + D. kyathit* | *D. choprae* | 4 |
| group3 | *D. rerio* | *D. tinwini* | *D. rerio + D. tinwini* | *D. choprae* | 6 |
| group3 | *D. rerio* | *D. tinwini* | *D. rerio + D. tinwini* | *D. jaintianensis* | 9 |
| group3 | *D. rerio* | *D. nigrofasciatus* | *D. rerio + D. nigrofasciatus* | *D. jaintianensis* | 10 |
| group3 | *D. rerio* | *D. nigrofasciatus* | *D. rerio + D. nigrofasciatus* | *D. choprae* | 2 |
| group3 | *D. rerio* | *D. albolineatus* | *D. rerio + D. albolineatus* | *D. choprae* | 3 |
| group3 | *D. rerio* | *D. albolineatus* | *D. rerio + D. albolineatus* | *D. jaintianensis* | 8 |
| group3 | *D. rerio* | *D. albolineatus* | *D. rerio* | *D. jaintianensis* | 1 |
| group3 | *D. rerio* | *D. erythromicron* | *D. rerio + D. erythromicron* | *D. choprae* | 2 |
| group3 | *D. rerio* | *D. erythromicron* | *D. rerio + D. erythromicron* | *D. jaintianensis* | 3 |
| group3 | *D. rerio* | *D. margaritatus* | *D. jaintianensis* | *D. margaritatus* | 1 |
| group3 | *D. rerio* | *D. margaritatus* | *D. rerio + D. margaritatus* | *D. jaintianensis* | 4 |
| group3 | *D. rerio* | *D. margaritatus* | *D. rerio + D. margaritatus* | *D. choprae* | 1 |
| group3 | *D. aesculapii* | *D. tinwini* | *D. aesculapii* | *D. jaintianensis* | 1 |
| group3 | *D. aesculapii* | *D. tinwini* | *D. aesculapii + D. tinwini* | *D. jaintianensis* | 10 |
| group3 | *D. aesculapii* | *D. tinwini* | *D. aesculapii + D. tinwini* | *D. choprae* | 4 |
| group3 | *D. aesculapii* | *D. tinwini* | *D. jaintianensis* | *D. aesculapii* | 1 |
| group3 | *D. aesculapii* | *D. nigrofasciatus* | *D. aesculapii + D. nigrofasciatus* | *D. jaintianensis* | 8 |
| group3 | *D. aesculapii* | *D. nigrofasciatus* | *D. aesculapii + D. nigrofasciatus* | *D. choprae* | 4 |
| group3 | *D. aesculapii* | *D. albolineatus* | *D. aesculapii + D. albolineatus* | *D. jaintianensis* | 7 |
| group3 | *D. aesculapii* | *D. albolineatus* | *D. aesculapii + D. albolineatus* | *D. choprae* | 3 |
| group3 | *D. aesculapii* | *D. albolineatus* | *D. aesculapii* | *D. jaintianensis* | 2 |
| group3 | *D. aesculapii* | *D. erythromicron* | *D. aesculapii + D. erythromicron* | *D. jaintianensis* | 2 |
| group3 | *D. aesculapii* | *D. erythromicron* | *D. aesculapii + D. erythromicron* | *D. choprae* | 1 |
| group3 | *D. aesculapii* | D. margaritatus | *D. aesculapii + D. margaritatus* | *D. jaintianensis* | 2 |
| group3 | *D. aesculapii* | D. margaritatus | *D. aesculapii + D. margaritatus* | *D. choprae* | 1 |
| group3 | *D. kyathit* | *D. aesculapii* | *D. kyathit + D. aesculapii* | *D. jaintianensis* | 10 |
| group3 | *D. kyathit* | *D. aesculapii* | *D. kyathit* | *D. jaintianensis* | 2 |
| group3 | *D. kyathit* | *D. aesculapii* | *D. kyathit + D. aesculapii* | *D. choprae* | 4 |
| group3 | *D. kyathit* | *D. tinwini* | *D. kyathit* | *D. jaintianensis* | 1 |
| group3 | *D. kyathit* | *D. tinwini* | *D. kyathit + D. tinwini* | *D. jaintianensis* | 11 |
| group3 | *D. kyathit* | *D. tinwini* | *D. kyathit + D. tinwini* | *D. choprae* | 3 |
| group3 | *D. kyathit* | *D. nigrofasciatus* | *D. kyathit + D. nigrofasciatus* | *D. jaintianensis* | 9 |
| group3 | *D. kyathit* | *D. nigrofasciatus* | *D. kyathit + D. nigrofasciatus* | *D. choprae* | 4 |
| group3 | *D. kyathit* | *D. nigrofasciatus* | *D. kyathit* | *D. jaintianensis* | 2 |
| group3 | *D. kyathit* | *D. albolineatus* | *D. kyathit + D. albolineatus* | *D. jaintianensis* | 9 |
| group3 | *D. kyathit* | *D. albolineatus* | *D. kyathit + D. albolineatus* | *D. choprae* | 3 |
| group3 | *D. kyathit* | *D. albolineatus* | *D. choprae* | *D. kyathit* | 1 |
| group3 | *D. kyathit* | *D. erythromicron* | *D. jaintianensis* | *D. kyathit* | 1 |
| group3 | *D. kyathit* | *D. erythromicron* | *D. kyathit + D. erythromicron* | *D. jaintianensis* | 4 |
| group3 | *D. kyathit* | *D. erythromicron* | *D. kyathit* | *D. jaintianensis* | 1 |
| group3 | *D. kyathit* | *D. margaritatus* | *D. jaintianensis* | *D. kyathit* | 1 |
| group3 | *D. kyathit* | *D. margaritatus* | *D. kyathit + D. margaritatus* | *D. jaintianensis* | 5 |
| group3 | *D. kyathit* | *D. margaritatus* | *D. kyathit + D. margaritatus* | *D. choprae* | 1 |
| group3 | *D. tinwini* | *D. nigrofasciatus* | *D. tinwini + D. nigrofasciatus* | *D. jaintianensis* | 10 |
| group3 | *D. tinwini* | *D. nigrofasciatus* | *D. tinwini + D. nigrofasciatus* | *D. choprae* | 4 |
| group3 | *D. tinwini* | *D. nigrofasciatus* | *D. tinwini* | *D. jaintianensis* | 3 |
| group3 | *D. tinwini* | *D. nigrofasciatus* | *D. choprae* | *D. tinwini* | 1 |
| group3 | *D. tinwini* | *D. erythromicron* | *D. tinwini + D. erythromicron* | *D. jaintianensis* | 3 |
| group3 | *D. tinwini* | *D. erythromicron* | *D. tinwini + D. erythromicron* | *D. choprae* | 1 |
| group3 | *D. tinwini* | *D. margaritatus* | *D. tinwini + D. margaritatus* | *D. jaintianensis* | 5 |
| group3 | *D. tinwini* | *D. margaritatus* | *D. tinwini* | *D. jaintianensis* | 1 |
| group3 | *D. tinwini* | *D. margaritatus* | *D. tinwini + D. margaritatus* | *D. choprae* | 1 |
| group3 | *D. nigrofasciatus* | *D. erythromicron* | *D. nigrofasciatus + D. erythromicron* | *D. jaintianensis* | 4 |
| group3 | *D. nigrofasciatus* | *D. erythromicron* | *D. nigrofasciatus + D. erythromicron* | *D. choprae* | 2 |
| group3 | *D. nigrofasciatus* | *D. margaritatus* | *D. nigrofasciatus + D. margaritatus* | *D. jaintianensis* | 4 |
| group3 | *D. nigrofasciatus* | *D. margaritatus* | *D. nigrofasciatus + D. margaritatus* | *D. choprae* | 1 |
| group3 | *D. albolineatus* | *D. tinwini* | *D. albolineatus + D. tinwini* | *D. jaintianensis* | 6 |
| group3 | *D. albolineatus* | *D. tinwini* | *D. albolineatus + D. tinwini* | *D. choprae* | 1 |
| group3 | *D. albolineatus* | *D. nigrofasciatus* | *D. albolineatus + D. nigrofasciatus* | *D. jaintianensis* | 10 |
| group3 | *D. albolineatus* | *D. nigrofasciatus* | *D. albolineatus + D. nigrofasciatus* | *D. choprae* | 2 |
| group3 | *D. albolineatus* | *D. erythromicron* | *D. albolineatus + D. erythromicron* | *D. jaintianensis* | 9 |
| group3 | *D. albolineatus* | *D. erythromicron* | *D. albolineatus + D. erythromicron* | *D. choprae* | 3 |
| group3 | *D. albolineatus* | *D. margaritatus* | *D. albolineatus + D. margaritatus* | *D. jaintianensis* | 8 |
| group3 | *D. albolineatus* | *D. margaritatus* | *D. albolineatus + D. margaritatus* | *D. choprae* | 1 |
| group3 | *D. erythromicron* | *D. margaritatus* | *D. erythromicron + D. margaritatus* | *D. choprae* | 2 |
| group3 | *D. erythromicron* | *D. margaritatus* | *D. erythromicron + D. margaritatus* | *D. jaintianensis* | 7 |

**Table S7 | Summary of gene family analysis (9 species).**

| **Species** | **Gene number** | **Unassigned genes** | **Orthogroup number** | **Species-specific orthogroup (gene) number** |
| --- | --- | --- | --- | --- |
| *Danio aesculapii* | 25,193 | 801 | 18,360 | 29 (101) |
| *Danio choprae* | 25,059 | 1,691 | 18,065 | 52 (117) |
| *Danio rerio* | 26,511 | 503 | 18,807 | 60 (313) |
| *Danio erythromicron* | 22,669 | 1,178 | 17,553 | 14 (32) |
| *Ictalurus punctatus* | 23,201 | 811 | 16,743 | 181 (1133) |
| *Danio kyathit* | 26,574 | 704 | 18,661 | 47 (193) |
| *Danio margaritatus* | 23,041 | 1,067 | 17,914 | 8 (21) |
| *Danio nigrofasciatus* | 26,390 | 1,779 | 18,553 | 47 (123) |
| *Danio tinwini* | 24,374 | 1,049 | 18,226 | 30 (64) |

**Table S8 | dN/dS value of pigment patterning genes (w = dN/dS*, p*-value < 0.05).**

| **Gene** | **Branch** | **w null** | **w branch** | **w background** | ***p*-value** |
| --- | --- | --- | --- | --- | --- |
| *gja4* | Node 2 | 0.18797 | 1.76388 | 0.18797 | 0 |
| *gja4* | Node 4 | 0.18797 | 1.97993 | 0.18797 | 0 |
| *gja4* | Node 5 | 0.18797 | 1.99478 | 0.18797 | 0 |
| *gja4* | Node 6 | 0.18797 | 0.1972 | 0.18767 | 0.038671964 |
| *gja4* | *D. rerio* | 0.18797 | 0.18815 | 0.18796 | 0.001128379 |
| *igsf11* | Node 2 | 0.06265 | 1.8059 | 0.06254 | 0 |
| *igsf11* | Node 3 | 0.06265 | 1.65617 | 0.06124 | 0 |
| *igsf11* | Node 4 | 0.06265 | 2.25818 | 0.06218 | 0 |
| *kcnj13* | Node 2 | 0.13054 | 1.00E-04 | 0.13054 | 0.006432681 |
| *kcnj13* | Node 4 | 0.13054 | 1.99799 | 0.13054 | 0 |
| *kcnj13* | Node 5 | 0.13054 | 2.10628 | 0.13054 | 0 |
| *kcnj13* | Node 6 | 0.13054 | 1.41484 | 0.13054 | 0 |
| *kcnj13* | *D. nigrofasciatus* | 0.13054 | 1.65223 | 0.13054 | 0 |
